# Benchmarking precision matrix estimation methods for differential co-expression network analysis

**DOI:** 10.64898/2026.04.13.716081

**Authors:** Matthias Overmann, Gordon Grabert, Tim Kacprowski

## Abstract

**Background:** Gene expression profiling is widely used to investigate disease mechanisms, but classical approaches such as differential expression or pairwise correlation analyses provide limited interpretability. Network-based differential co-expression methods that model conditional dependencies through partial correlations offer richer insights, yet their application in high-dimensional settings requires estimation of precision matrices. Numerous precision matrix estimation methods (PMEMs) have been proposed, but their relative performance under various conditions remains unclear.

**Results:** Simulated gene expression datasets with known ground truth correlation structures were used to benchmark a broad set of PMEMs. Performance was strongly affected by data characteristics, including covariance structure, matrix density, covariance values, sample size-to-dimension ratio, and sampling distribution. Among the evaluated methods, *GLassoElnetFast* consistently showed the highest accuracy in recovering differential edges, although high signal-to-noise ratios and sufficient sample sizes remain essential for reliable inference.

**Conclusions:** Evaluation across diverse simulation conditions demonstrated that no single metric or condition was sufficient to assess PMEM performance. Therefore, previous less extensive evaluations risked misleading conclusions. Our simulation and benchmarking framework supports future method development and ensures reproducible evaluation of newly developed approaches.

## 1 Introduction

The transcriptome of cells is constantly responding to changing conditions or developmental stages, allowing the cells to functionally adapt [1]. This variable gene expression is profiled using two key technologies: microarrays or RNA sequencing (RNA-seq) [2]. The most common approach for analysing transcriptomic data is differential gene expression (DGE) analysis [3, 4]. Its main objective is to identify differentially expressed (DE) genes, which show changes in expression levels between conditions or are associated with relevant predictors or responses [3, 5–7]. To this end, a wide range of statistical pipelines and software tools have been developed [8–11], including DESeq2 [12], edgeR [13], and limma/voom [14, 15]. The typical output of such analyses is a list of significantly up- and downregulated genes, which serves as the basis for downstream interpretation, including pathway analysis and biomarker discovery [4, 16, 17]. Additionally, the relationship between the activity of two genes is often summarized using Pearson’s correlation, which assumes linear dependence and quantifies its strength and direction [6, 18–20]. The comparison of the expression of a single gene or the relationship of a gene pair across two conditions poses a simple statistical question, a two-sample comparison. However, analysing each gene or relationship in isolation neglects advantages that could be gained if the entirety of the gene expression profiles were taken into account [5]. Differential co-expression analysis addresses this by constructing networks that capture condition-specific changes in gene–gene relationships. These approaches include module and hub gene identification, regulatory network construction, and guilt by association analysis, with the aim of identifying subnetworks of connected genes that operate abnormally across sample groups [21]. Using Gaussian graphical models (GGMs), partial correlations obtained from the precision matrix highlight direct dependencies while filtering out indirect effects, resulting in sparser and more interpretable networks [22]. While incorporating the entirety of gene expression profiles offers numerous advantages, it also introduces new challenges. Gene expression studies usually analyse the gene expression profiles, including more than 20,000 genes, from typically a few hundred samples [23, 24], leading to a high-dimensional setting [25]. In this setting, the sample covariance matrix becomes singular, and its inverse cannot be used to calculate the precision matrix. To overcome this, precision matrix estimation methods have been developed that impose regularization or structural assumptions, allowing the recovery of stable and interpretable partial correlation networks. However, the relative performance of existing precision matrix estimation methods for differential co-expression analysis has not been systematically evaluated across diverse high-dimensional data scenarios.

In this study, a simulation framework is introduced to evaluate a wide range of such methods across diverse data scenarios. Their performance is compared using multiple evaluation metrics, with the aim of identifying strengths, weaknesses, and practical considerations for differential co-expression network analysis.

### 2 Background

### 2.1 Networks

A network, or graph, specifies relationships between system components (nodes) via interactions (edges) [26, 27]. Nodes with high degrees are referred to as hubs [26]. Networks are often sparse and can be mathematically described using an adjacency matrix. Real-world networks, including biological, social, and technological systems, commonly display non-random structures, such as scale-free topologies, characterized by a power-law degree distribution: *p*(*k*) ≈ *k*^−*γ*^ [26, 28, 29]. This distribution results in few hubs and many sparsely connected nodes [27, 30], emerging from mechanisms like growth and preferential attachment [30]. In contrast, Erdős–Rényi random networks exhibit uniform degree distributions and lack hubs [30]. While differential gene expression analysis has revealed insights into transcriptional regulation and disease biomarkers [31–34], it tends to highlight generic or well-characterized genes [35]. Biological networks change in response to development, external stimuli, or disease [36]. Importantly, many diseases result from network-level disruptions rather than from changes in individual genes [37]. Thus, comparing network topologies under different conditions can reveal rewiring patterns and underlying molecular mechanisms, aiding in the understanding of disease processes and identification of therapeutic targets [38–40]. Network-based approaches, especially differential network analysis, address this by evaluating changes in interactions rather than individual components [36, 38, 39]. This allows such approaches to reveal disease-related modules and pathways even when gene expression itself remains unchanged. Differential network analysis relies on the comparison of condition-specific networks [38]. To obtain these networks, associations between variables must first be inferred within each condition. To this end, a wide range of differential network analysis tools have been proposed for transcriptomic and genomic data. The regression tree-based method BoostDiff infers differential networks by building adaptively boosted ensembles of differential trees [41]. DINGO focuses on identifying condition-specific network changes within predefined biological pathways [42]. Weighted gene co-expression network analysis (WGCNA), one of the most widely used approaches, constructs weighted co-expression networks using pairwise correlations and scale-free topology assumptions [21, 43]. Methods based on Gaussian graphical models explicitly capture changes in conditional dependencies between genes. One such method, the hierarCHical differential NETwork analysis model (chNet), tests for changes in partial correlations between gene pairs as well as changes in expression levels of individual genes [38]. Many approaches, such as WGCNA and chNet, use correlation-based metrics to evaluate the strength of associations between node pairs [38, 42, 43]. The following section provides a more detailed introduction to these metrics.

### 2.2 Correlation-based Metrics

Empirical quantitative data are often summarized using descriptive statistics such as the arithmetic mean 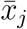 and variance 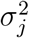, which measures data dispersion. In the following, we distinguish between population-level quantities defined for random variables and their empirical counterparts computed from observed data. Uppercase letters (e.g., *X*) denote random variables or random vectors, while lowercase letters (e.g., *x*_*ij*_) denote observed realizations. Population parameters are defined via expectations, whereas sample quantities are defined as finite sums over the data. Let *X*_*j*_ denote a random variable representing the *j*th variable in the population, and let *x*_*ij*_ be the observed value of variable *j* in observation *i*. The population mean of variable *j* is defined as

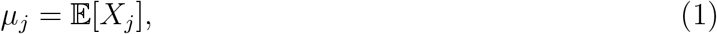

and the population variance is given by

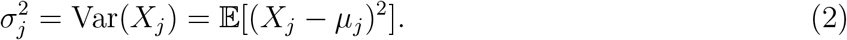

Covariance quantifies joint variability between two variables and is key to assessing linear relationships [44]. The covariance between two random variables *X*_1_ and *X*_2_ is defined as

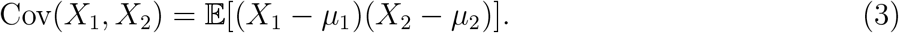

In practice, population parameters are unknown and must be estimated from a sample of size *n*. Given observed realizations *x*_1*j*_, …, *x*_*nj*_, the sample mean approximates *µ*_*j*_ and is defined as

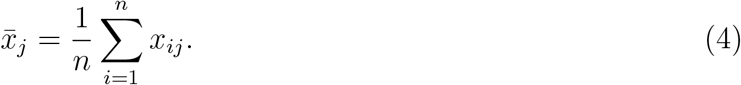

The sample variance and covariance are defined as

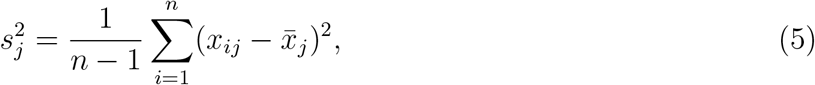

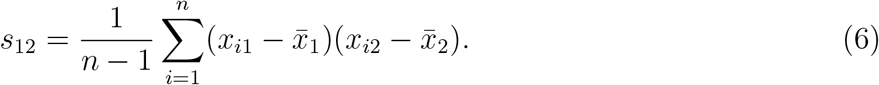

Covariance is unit-dependent and not standardized. The Pearson correlation coefficient normalizes it:

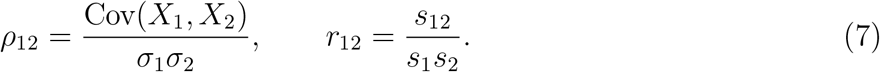

In multivariate analysis, let *X* = (*X*_1_, …, *X*_*p*_)^*T*^ ∈ ℝ^*p*^ denote a *p*-dimensional random vector with mean vector *µ* = 𝔼[*X*] and covariance matrix Σ = Cov(*X*) = 𝔼[(*X* − *µ*)(*X* − *µ*)^*T*^]. The (*i, j*)th element of Σ is *σ*_*ij*_ = Cov(*X*_*i*_, *X*_*j*_), so that the population covariance matrix can be written as

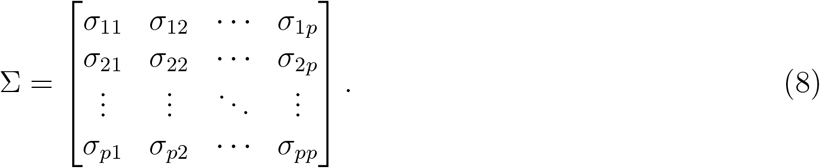

Suppose we observe *n* independent and identically distributed realizations

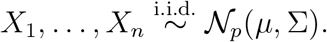

The observed data are stored in the matrix **X** ∈ ℝ^*n*×*p*^, where each row corresponds to one observation. The sample covariance matrix *S* = (*s*_*ij*_), which serves as an estimator of Σ, is defined as

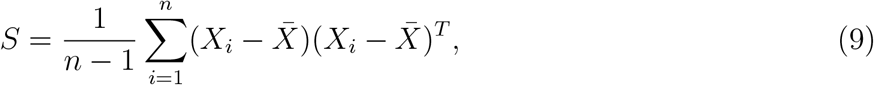

where 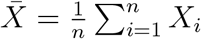 denotes the sample mean vector. The correlation matrix *R* = (*ρ*_*ij*_), where

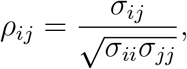

contains pairwise correlations and thus represents the normalized covariance matrix. The inverse of Σ, Σ^−1^, known as the precision matrix or concentration matrix Θ, reflects conditional dependencies:

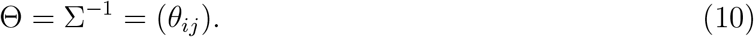

Under the assumption that *X* ∼ 𝒩_*p*_(*µ*, Σ), an element *θ*_*ij*_ = 0 implies that *X*_*i*_ and *X*_*j*_ are conditionally independent given all remaining variables {*X*_*k*_ : *k* ≠ *i, j*}. Conversely, *θ*_*ij*_ ≠ 0 indicates a direct conditional dependency between *X*_*i*_ and *X*_*j*_. This property allows the precision matrix to be interpreted as the adjacency structure of an undirected graphical model, where nodes represent variables and edges correspond to non-zero off-diagonal elements of Θ [45]. A matrix **A** ∈ ℝ^*p*×*p*^ is invertible if there exists a matrix **B** ∈ ℝ^*p*×*p*^ such that

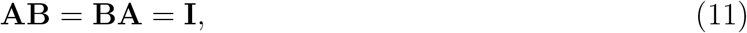

where **I** denotes the *p* × *p* identity matrix. The matrix **B** = **A**^−1^ is uniquely determined by **A** [46]. A symmetric matrix **A** is positive definite if all eigenvalues are strictly positive, or equivalently, if

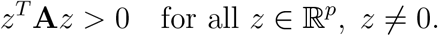

In classical multivariate statistical studies, asymptotic settings presume that the number of samples *n* increases indefinitely and is substantially greater than the number of variables *p*, which remains constant. In these settings, the sample covariance matrix *S* consistently estimates Σ and is positive definite with probability tending to one. Accordingly, *S*^−1^ can serve as a reliable estimator for the precision matrix [47, 48]. However, technological and computational advancements over the past decades have significantly increased the availability of high-dimensional data. In high-dimensional low sample size (HDLSS) settings (*p* ≫ *n*), the sample covariance matrix satisfies rank(*S*) ≤ *n* − 1, and is therefore singular whenever *p* ≥ *n*. Consequently, *S*^−1^ cannot be computed and therefore cannot serve as an estimator of the precision matrix [48]. These limitations of *S* in HDLSS settings have focused attention on alternative estimators for large covariance or precision matrices [49, 50]. Methods include banding [51], tapering [52], thresholding [50], modified Cholesky decomposition with regularization [53–55], and sparse precision matrix estimation using the graphical lasso [56]. Sparse precision matrices are particularly relevant in graphical models, as zero partial correlations correspond to the absence of edges between nodes in the inferred network [45, 50]. Sparsity therefore constrains the network structure and enables the identification of subnetworks or clusters of strongly connected variables that are conditionally independent of others.

### 2.3 Gaussian Graphical Models

Probabilistic graphical models (PGMs) combine probability theory and graph theory to represent joint probability distributions through graphs, assuming conditional independence among variables. This reduces computational complexity in inference and learning, making PGMs widely used for modelling uncertainty in fields such as computer vision, time-series analysis, and bioinformatics [57]. For large-scale biological data generated by high-throughput omics technologies, PGMs, and in particular Gaussian graphical models (GGMs), are commonly used to infer statistical dependencies among biological variables under the assumption of multivariate Gaussian distributions [38, 58–60]. GGMs estimate conditional dependencies using full-order partial correlations, enabling the distinction between direct and indirect associations. This contrasts with correlation-based approaches, which only capture marginal relationships and may conflate direct and indirect effects [58, 61]. Partial correlations adjust for the influence of all other variables, offering more accurate network representations [62].

In correlation networks, the absence of an edge between two nodes indicates that the nodes lack both direct and indirect associations. However, this is rare in biological data, and consequently correlation networks are typically dense [61]. Sparsity is typically enforced through thresholding, which introduces arbitrary cut-offs and interpretability challenges [61]. In contrast, GGMs naturally yield sparse networks by modelling conditional independencies directly. The resulting partial correlation networks represent variables as nodes, with edges indicating direct dependencies. Conditional independencies are encoded in the precision matrix Θ, where zeros correspond to the absence of edges and thus to conditional independence between variables [38, 58, 60].

### 2.4 Precision Matrix Estimation

The partial correlations, used by GGMs to quantify linear interrelationships after removing the effect of all remaining variables [63]. For a *p*-dimensional random vector *X* = (*X*_1_, …, *X*_*p*_)^*T*^, the partial correlation between two variables *X*_*i*_ and *X*_*j*_, given all other variables *X*_*k*_ : *k* ≠ *i, j*}, is defined as

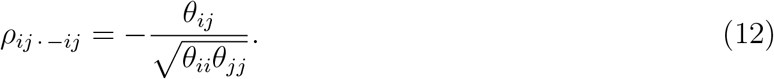

The partial correlation matrix Ψ = (*ρ*_*ij* · −*ij*_) therefore contains the normalized off-diagonal entries of the precision matrix. In matrix form, it can be written as

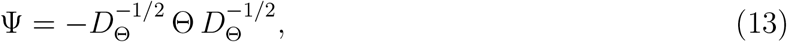

where *D*_Θ_ = diag(*θ*_11_, …, *θ*_*pp*_) denotes the diagonal matrix containing the diagonal elements of Θ [64, 65].

In high-dimensional conditions, *S*^−1^ cannot be calculated and therefore cannot be used as an estimator of Θ. As a result, Θ must be estimated directly to obtain Ψ [47, 66–68]. Consequently, directly estimating the precision matrix is a central problem in high-dimensional statistics that has received much attention in the literature [48, 69, 70]. A variety of strategies have been developed to address HDLSS conditions, including dimension reduction, feature selection, and standard fitting methods with regularization. In particular, regularization approaches such as *ℓ*_1_- and *ℓ*_2_-penalized regression are widely applied to address feature selection and overfitting. Least absolute shrinkage and selection operator (Lasso) regression, proposed by Tibshirani (1996) [71], uses *ℓ*_1_-penalized regularization and is able to shrink less crucial feature coefficients to zero, effectively eliminating some features. In contrast, ridge regression, introduced by Hoerl and Kennard (2000) [72], utilizes *ℓ*_2_-penalized regularization and helps prevent overfitting by penalizing large coefficients. The elastic net, a regularization and variable selection method introduced by Zou and Hastie (2005) [73], combines *ℓ*_1_ and *ℓ*_2_ penalties [74].

## 3 Data Description

In this study, datasets were simulated to allow evaluation against a known ground truth, in order to provide a reliable benchmark for comparing precision matrix estimation methods. The simulation design follows the intuition of differential gene co-expression. Specifically, two conditions were generated with the same marginal distributions of variables but differing in their underlying correlation structures. In this way, variation between conditions was not reflected in mean differences but in the structure of the condition-specific networks. This setup provides a suitable setting for assessing whether precision matrix estimation methods can recover the predefined structures and, more generally, whether they are suitable to infer partial correlations for differential co-expression analysis. In addition, the simulation design allows systematic variation of key data characteristics, including sample size, dimensionality, covariance structure, and sampling distribution. These factors are used in the subsequent analyses to examine how data properties influence the performance of different estimation methods. Accordingly, the simulation pipeline is designed to generate condition-specific covariance structures, simulate data, and evaluate estimation performance (see Figure 1 for an overview). In a single simulation run the simulation pipeline starts by generating a covariance matrix for the first condition Σ_1_, using a desired dimension *p* and one of the following covariance 1 methods: single block, multiple block, band network, scale free 1, scale free 2, single block icf, multiple block icf, random graph icf, and random hubs icf (see Methods: Covariance Matrix Generation). Furthermore, the methods: scale free 2, single block icf, multiple block icf, random graph icf, and random hubs icf were developed to generate the covariance matrix for the second condition Σ_2_ internally. Otherwise, Σ_2_ is created based on Σ_1_ using one of the two covariance 2 methods: *knockout* and *mutate* (see Methods: Covariance Matrix Alteration). Σ_1_ and Σ_2_ are generated to be positive definite. Thus, the inverse covariance matrices, 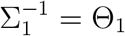 and 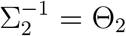, can be computed and used as ground truth in the subsequent analysis. Additionally, the precision matrices are used to calculate the differential partial correlation matrix Ψ_Δ_. Furthermore, Σ_1_ and Σ_2_ are used to generate the simulated gene expression data sets **X**_1_ and **X**_2_ with a specified sample count *n* (see Methods: Sampling). These data sets reflect the respective covariance structure between the variables. Either a multivariate normal distribution or Poisson distribution of the individual variable counts is simulated. The sample covariance matrices *S*_1_ and *S*_2_ are calculated based on the simulated data sets. **X**_1_ and **X**_2_ as well as *S*_1_ and *S*_2_ are used as input for the precision matrix estimation methods (Table 2). These estimate the precision matrices for both conditions 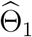 and 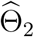. The performance of the PMEMs is evaluated by comparing the respective 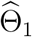 and 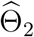 with Θ_1_ and Θ_2_ (see Methods: Evaluation). Additionally, the estimated differential partial correlation matrix 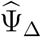 is compared with Ψ_Δ_.

**Figure 1:**
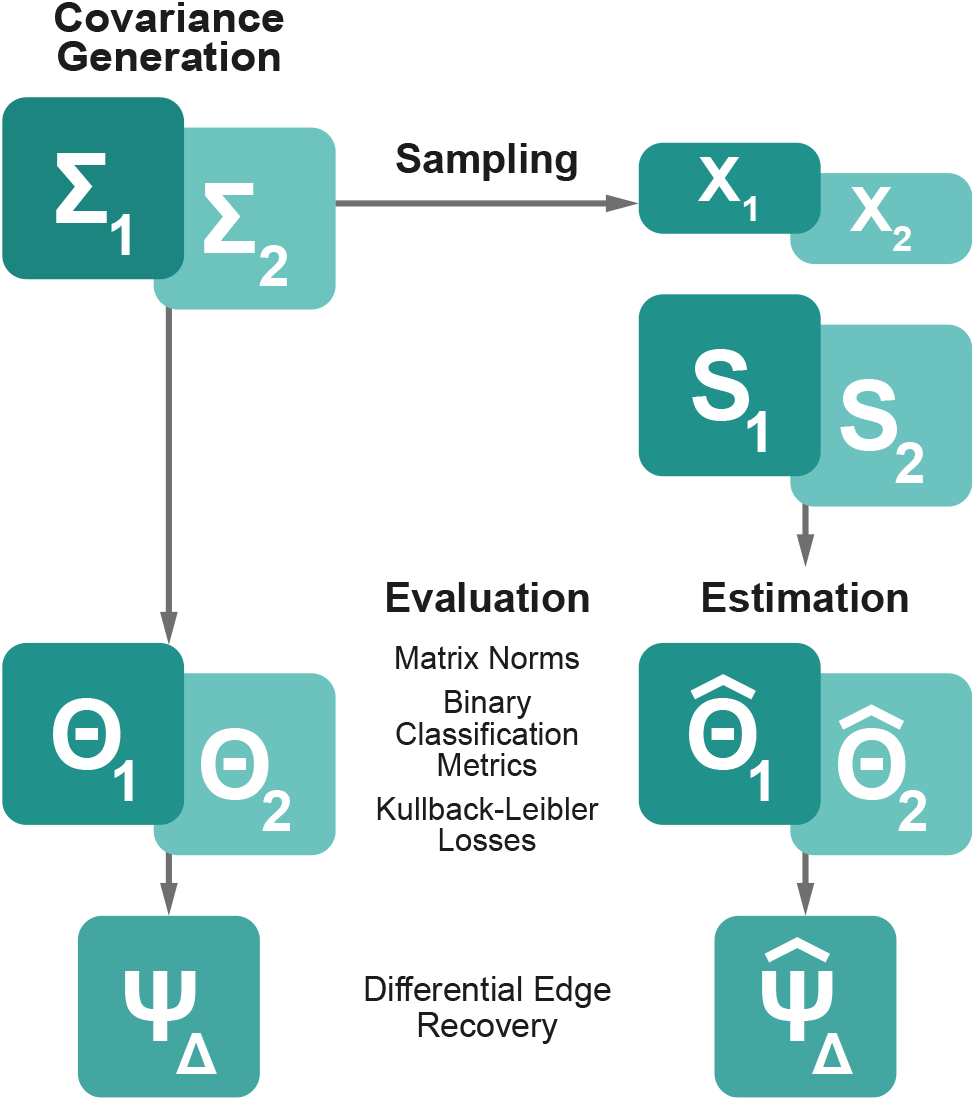
Flowchart, illustrating the general procedure of the simulation studies. Σ: covariance matrix, Θ: precision matrix, Ψ: partial correlation matrix, **X**: simulated data set, *S*: sample covariance matrix, 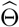: estimated precision matrix, PMEM: precision matrix estimation method.

## 4 Analyses

Table 1 provides an overview of all analyses performed in this study. A baseline scenario was defined using block-structured covariance matrices, with differences between conditions introduced at the level of the condition-specific covariance matrices through the knockout procedure (see Methods: Covariance Alteration). The dimensionality was set to *p* = 100 and the sample size to *n* = 50, with data sampled from a multivariate Gaussian distribution with a zero mean vector. Subsequent analyses systematically varied one parameter of this baseline scenario at a time, in order to assess the influence of individual data properties on the performance of precision matrix estimation methods. Further detailed figures and additional results can be found in the Supplementary Data. The analyses are presented in the following order: replicates, covariance structures, covariance densities, covariance alterations, covariance values, dimensionality, mean distributions, normalization, sample size, sampling distributions, and runtimes (see Table 1 for an overview).

**Table 1:**
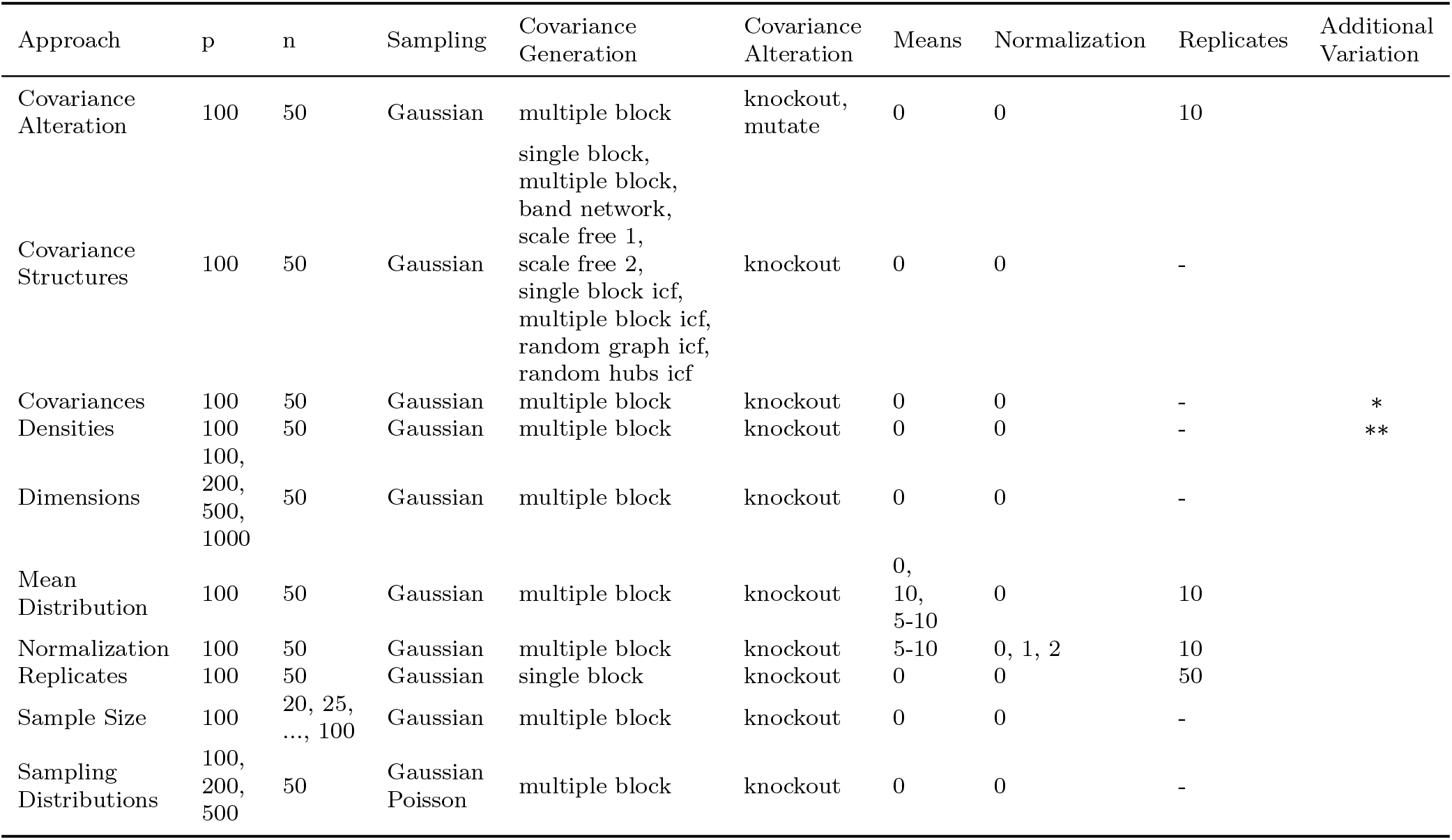
Overview of the analysed simulation settings. *p* denotes the dimensionality and *n* the sample size. Sampling indicates the distribution used for data generation. Covariance Generation specifies the methods used to construct Σ_1_, while Covariance Alteration describes how Σ_2_ is derived from Σ_1_. Means indicate the mean vectors used for sampling from Σ_1_ and Σ_2_. Normalization specifies whether no normalization (0), normalization of means and variances (1), or normalization of means only (2) was applied. Replicates denote the number of simulation replicates. Additional variation indicates settings with modified covariance structures: ∗ indicates settings in which the off-diagonal entries were fixed to *c* ∈ {1, …, 10} and the diagonal entries to *c* + 1 ∈ {2, …, 11}, with *c* varied across settings; ∗∗ corresponds to *k* non-zero blocks in Σ_1,2_ with *k* ∈ {3, …, 20}, resulting in precision matrix densities ranging from 0.05 to 0.33.

### 4.1 Replicates

Since some methods required substantial computation time, especially when estimating highdimensional precision matrices (e.g., several hours for *p* = 500), extensive replication was not always feasible. However, to obtain a general impression of baseline performance and consistency across methods, the PMEMs were first evaluated in 50 replicated simulations. Most PMEMs show stable performance across binary classification metrics, with the greatest variation observed in the F1 score (Figure S1 A). *GLassoElnetFast* achieves the highest values across all metrics, while *clime, rags2ridges*, and *rope* perform poorly, with normalized MCC values of 0.5 and reduced accuracy due to estimating fully dense precision matrices without thresholding. Matrix norm values vary only slightly for most PMEMs, with the Frobenius norm showing the largest spread (Figure S1 B). *Gdtrace* stands out with high variability across all norms, whereas *GLassoElnetFast* consistently yields the lowest values. Kullback-Leibler and reverse Kullback-Leibler losses show little variation between methods, with *GLassoElnetFast* again producing the lowest values (Figure S1 C). *Bigquic, scio*, and *tiger* consistently estimated empty precision matrices. *Bigquic* produced diagonal values fixed at 0.5, while *scio* and *tiger* showed varying diagonal entries ranging from 0.26 to 1.02 and 0.27 to 1.05, respectively. These outputs led to constant or near-constant results across all metrics, limiting their interpretability in this evaluation.

### 4.2 Covariance Structures

To test whether the underlying covariance structures of the simulated datasets influence the performance of PMEMs, nine covariance generation methods, or covariance 1 methods, were developed and tested, namely single block, multiple block, band network, scale free 1, scale free 2, single block icf, multiple block icf, random graph icf and random hubs icf (see Methods: Covariance Matrix Generation). The results are presented in the Supplementary Figures 2, 3, and 4. For most PMEMs, performance across all three metrics varies considerably depending on the covariance generation method used. In the binary classification metrics, the PMEMs *bigquic, scio*, and *tiger* show similar results (Figure S2). For almost all PMEMs a bad performance in band network, scale free 1, random graph icf, and random hubs icf approaches can be seen. *Clime, rags2ridges*, and *rope* show identical results, and in contrast to the other PMEMs result in high F1 score and accuracy in the band network, scale free 1, random graph icf, and random hubs icf approaches. However, the normalized MCC reveals that all PMEMs perform equally poorly in these approaches, with values consistently at 0.5 across all methods. These patterns highlight that the structure induced by different covariance 1 methods can significantly affect binary classification metrics.

In general, the selected covariance generation methods led to different values of the matrix norms for each PMEM (Figure S3). Most notably, all analysed PMEMs showed the highest values for the matrix 1-norm, Frobenius norm, and spectral norm in the approaches based on the scale free 1 method, consistently several orders of magnitude higher than in the other settings. Since all methods were affected, the explanation lies not in the estimation methods themselves but in the generation of the ground truth matrices Σ_1_ and Θ_1_. The scale free 1 method uses the `nearPD()` function to enforce positive definiteness, but the resulting Σ_1_ remained nearly singular, with eigenvalues close to zero. As a consequence, the inverse matrix Θ_1_ contained extremely large entries. In contrast, the PMEMs estimated precision matrices with more typical magnitudes, resulting in consistently large differences between the estimated precision matrix 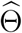 and the ground truth precision matrix Θ_1_, and thereby inflating the matrix norm values across all methods. Apart from this, the Σ_1_ generation method scale free 2 results in the highest values of the Frobenius norm for all PMEMs. The lowest values for the Frobenius norm are observed in approaches based on the single block or multiple block method. The highest matrix 1-norms are seen in approaches based on scale free 2, single block icf, and random hubs icf. Overall, a significant influence of the selected covariance 1 methods on the performance of the PMEMs can be seen on the basis of the presented matrix norms. In this context, the PMEMs display comparable performance trends across settings. This suggests a similar response to differences in the underlying covariance structures.

### 4.3 Covariance Densities

In the previous subsection, the influence of different covariance structures on PMEM performance was examined. However, the methods used to generate Σ_1_, and consequently Θ_1_, differ not only in their non-zero patterns but also in the resulting matrix densities (see Methods: Covariance Matrix Generation). In the approaches considered, the single block and multiple block methods yielded Θ_1_ densities of 0.12 and 0.10, respectively. The band network, random graph icf, and random hubs icf methods resulted in matrices with a density of 1. The scale free 1 and scale free 2 approaches produced densities of 0.98 and 0.55, respectively. The single block icf had a density of 0.26, while the multiple block icf matched the standard multiple block method with a density of 0.10.

These differences highlight the strong dependence of PMEM performance, as measured by binary classification metrics, on the density of the underlying precision matrix. Accordingly, this section further investigates how variation in the density of the true precision matrix Θ_1_ affects the performance of different PMEMs. The density of Θ_1_ was systematically increased from 0.05 to 0.33 by adjusting the number of non-zero blocks generated using the multiple block covariance method (Figure S5). Since the resulting precision matrices derived from this method consistently match the density of their corresponding covariance matrices, only the density of the true and estimated precision matrices is considered in the following analysis. An important property of the sample covariance matrix *S* is that, regardless of the density of the underlying true covariance matrix or the number of samples, *S* is always fully dense. Thus, PMEMs receive no explicit information about the true sparsity of the underlying structure. They must infer it entirely through their internal regularization, based solely on the signal present in *S*.

The binary classification metrics F1 score, accuracy, and normalized MCC declined for most PMEMs as the density of Θ_1_ increased (Figure S6). *GLassoElnetFast* was the exception, showing an increase in F1 score, a smaller decrease in accuracy compared to other PMEMs, and normalized MCC values that increased at lower densities and remained stable at higher ones. *Clime, rags2ridges*, and *rope* also showed increased F1 scores and accuracy, though their normalized MCC remained constant at 0.5. The matrix norm values generally increased with density for all methods except *rags2ridges*, which showed decreasing norm values across all density levels (Figure S7). Figure 2 reveals an interesting relationship between the density of 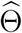 and the true density of Θ_1_. For most PMEMs, the estimated density remained largely unchanged regardless of the ground truth density, suggesting a lack of adaptivity in sparsity estimation. *GLassoElnetFast* was the only method to adjust its estimated density in response to the true structure, though it tended to overshoot. As a result, binary classification metrics are meaningfully interpretable only for *GLassoElnetFast* in this setting, since performance for all other methods is highly dependent on the mismatch between estimated and true density. Finally, it is worth noting that this was the only scenario in which *tiger* did not produce an empty matrix, beginning to yield non-zero estimates at densities of Θ_1_ ≥ 0.2.

**Figure 2:**
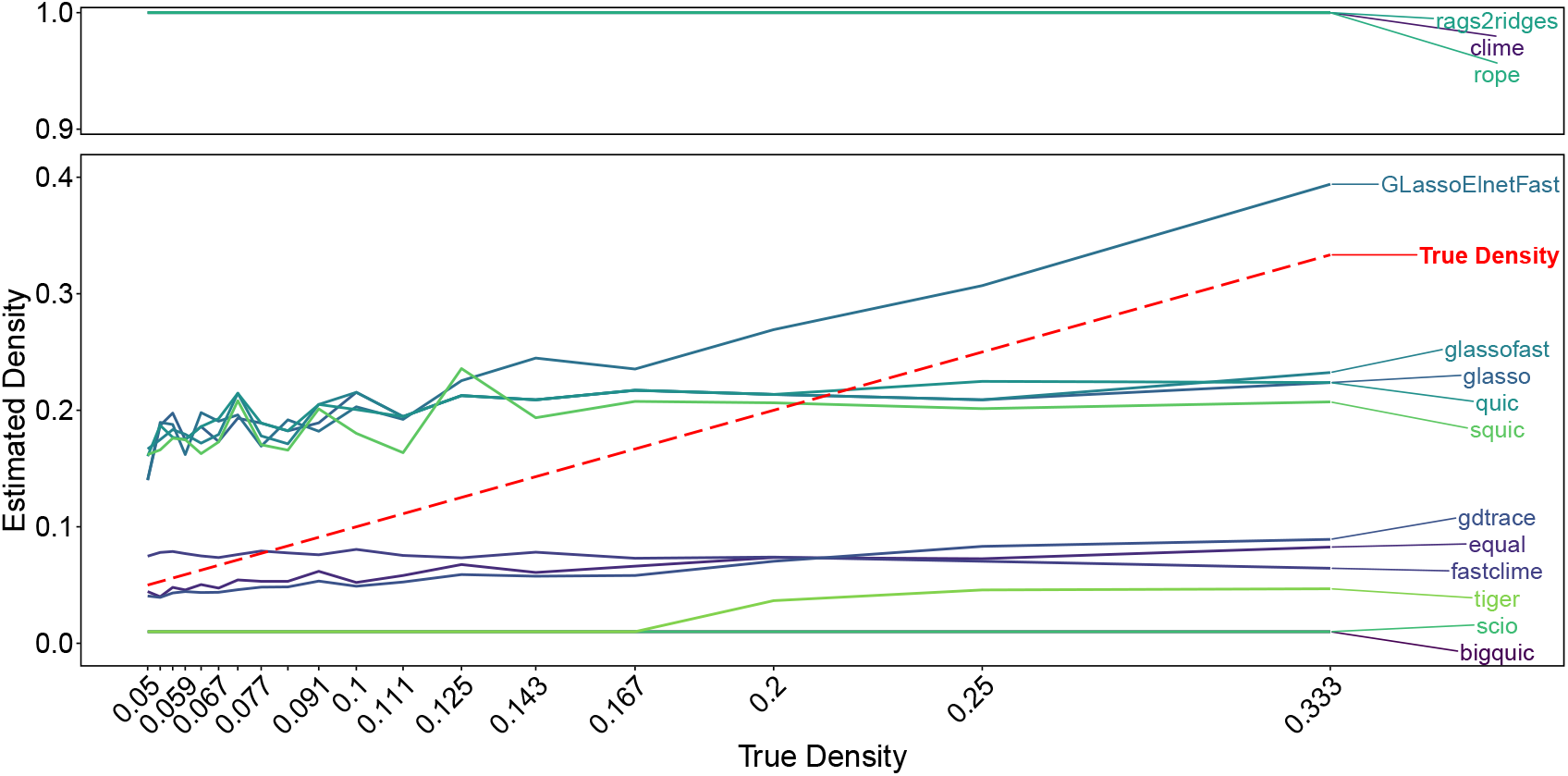
Line plot illustrating the influence of varying true densities of Θ_1_ on the densities of the estimated precision matrices for condition one 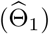. The PMEMs *clime, rags2ridges*, and *rope* consistently estimated the matrices with a density of 1 across all settings. In contrast, *bigquic, scio*, and *tiger* predominantly recovered only diagonal elements, resulting in an estimated density of 0.01. For densities greater than 0.2 in Θ_1_, *tiger* showed a slight increase in density in 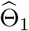. *Gdtrace* and *equal* exhibited similar behaviour, with only minimal increases in estimated density as the true density increased, while *fastclime* showed a slight decrease in the density of 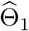 with increasing density of Θ_1_. Comparable patterns were observed for *glasso, glassofast, quic*, and *squic*, which displayed only modest sensitivity to changes in true density, generally overestimating density at low true densities (≤ 0.2) and underestimating density at higher true densities (0.25 and 0.333). *GLassoElnetFast* consistently overestimated the density of 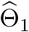, but was the only PMEM to exhibit a clear and systematic increase in estimated density with increasing true density of Θ_1_.

### 4.4 Covariance Alteration

To assess how the method of covariance alteration influences the performance of the PMEMs, the second condition was simulated using either the knockout or mutate approach to generate Σ_2_, from which the corresponding true precision matrix Θ_2_ was derived (see Methods: Covariance Matrix Alteration). The estimated precision matrices 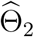 were then evaluated against Θ_2_ (Supplementary Figures 9, 10, 11, and 12). *Bigquic, scio*, and *tiger* again estimated completely empty matrices for 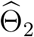. *Bigquic* produced diagonal entries fixed at 0.5, while *scio* and *tiger* yielded diagonal values ranging from 0.136 to 1.277 and 0.139 to 1.303, respectively. As before, *clime, rags2ridges*, and *rope* returned dense matrices, limiting the interpretability of binary classification metrics without additional thresholding.

The binary classification metrics indicate reduced performance for all PMEMs in the mutate approaches (Figure S9). *Clime, rope*, and *rags2ridges* appear to perform better in terms of F1 score and accuracy. It is important to note that in these approaches, the knockout method generated Θ_2_ with a density similar to Θ_1_, consistently around 0.08, while the mutate method resulted in considerably higher and more variable densities, ranging from 0.30 to 1.00. While this benefits the PMEMs *clime, rags2ridges*, and *rope* in F1 and accuracy, normalized MCC remains low. Matrix norm values mostly increased from knockout to mutate (Figure S10), although the Frobenius norm occasionally decreased depending on the PMEM. Kullback-Leibler and reverse Kullback-Leibler losses also tended to increase under the mutate condition (Figure S11). Differential edge recovery results are shown in Supplementary Figure S12. The highest recovery rates were achieved by *GLassoElnetFast* and *rags2ridges* in the knockout approaches. While *GLassoElnetFast* showed reduced recovery in the mutate approaches, performing similarly to *glasso, glassofast, quic*, and *squic. Rags2ridges* maintained higher recovery and achieved the best results across all PMEMs. *Bigquic, scio*, and *tiger* did not recover any differential edges, consistent with their empty matrix estimates.

### 4.5 Covariance Values

Additionally, we analysed whether the values of the covariances in the underlying covariance matrices (Σ_1_ and Σ_2_) have an influence on the performance of the PMEMs. This was approached by varying the off-diagonal values or covariances in Σ_1_ and thus also Σ_2_. In each approach, the diagonal values or variances were always set to the used covariance value plus one. This essentially increased the signal-to-noise ratio, while also increasing the variances (Figure S13). Overall, increasing the covariances had little to no effect on the performance of most analysed PMEMs in terms of F1 score, accuracy, and normalized MCC. The only notable exception was the PMEM *equal*, which showed a clear decline across all binary classification metrics as covariance values increased (Figure S14). Similarly, the performances of most PMEMs measured by the matrix norms were unaffected by varying covariance values (Figure S15). However, the PMEMs *clime, equal, fastclime, scio* and *tiger* result in higher values for Frobenius norm with an increase in the covariance values. *GLassoElnetFast* is shown to achieve the lowest values for all matrix norms in these settings. Additionally, a visible decrease in all matrix norms can be seen when the covariances increase from 2 to 3. In terms of differential edge recovery (see Methods: Differential Edge Recovery), most PMEMs did not substantially benefit from the increased signal-to-noise ratio associated with higher covariance values (Figure 3). *GLassoElnetFast* was the only method to consistently benefit from this effect, achieving the highest recovery rates across all tested covariance levels, with performance improving as covariances increased. *Rags2ridges* achieved the second-highest recovery rates overall but showed a decline at higher covariance values. A minimal increase in differential edge recovery was observed for *clime, rope, glasso, squic, quic*, and *glassofast. Gdtrace* and *fastclime* displayed largely stable recovery rates across all conditions, with only small variations. Again, *bigquic, scio*, and *tiger* failed to recover any differential edges. The performance of *equal* declined markedly, with recovery rates dropping to zero.

**Figure 3:**
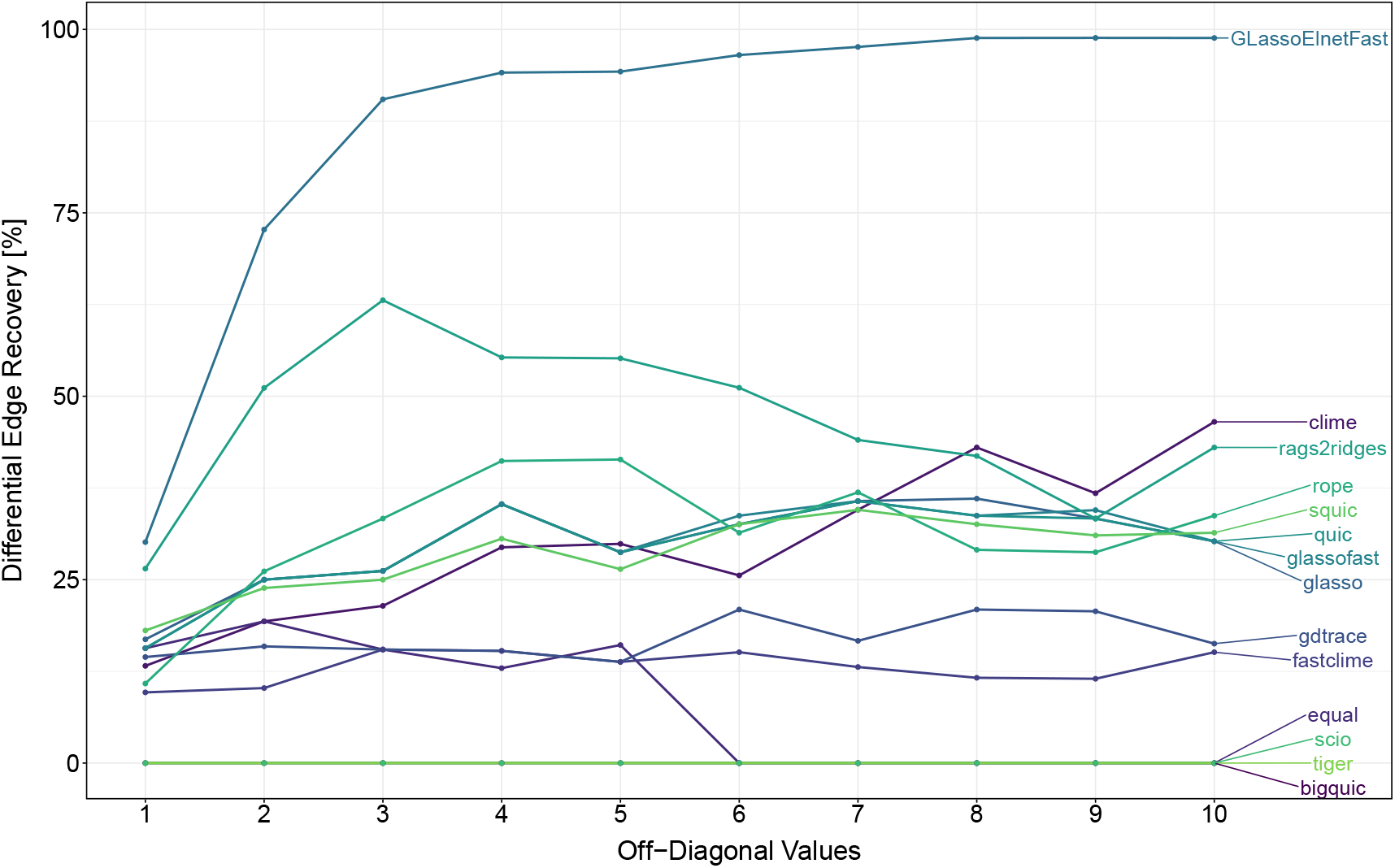
Line plot illustrating the influence of varying off-diagonal entry values (covariances) on the performance of precision matrix estimation methods, assessed by differential edge recovery rates. The PMEMs *scio, tiger*, and *bigquic* did not recover any differential edges across the considered settings. For *equal*, differential edge recovery decreased with increasing off-diagonal entry values and dropped to zero for values of 6 and higher. Recovery rates for *gdtrace* and *fastclime* were largely unaffected by variations in off-diagonal entry values. In contrast, *rags2ridges* initially recovered an increasing number of differential edges as off-diagonal entry values increased, followed by a decline in recovery rates for values greater than 3. Comparable increasing trends in differential edge recovery with increasing off-diagonal entry values were observed for *rope, squic, quic, glassofast, glasso*, and *clime*. Among all methods, *GLassoElnetFast* demonstrated the most pronounced increase in differential edge recovery as off-diagonal entry values increased.

### 4.6 Dimensions

This section examines how increasing dimensionality affects the performance of different PMEMs. This was analysed by increasing the number of variables while keeping the sample size constant, effectively decreasing the *n/p* ratio. The results are presented in Supplementary Figures 16, 17, and 18. The accuracy of all PMEMs remained largely unaffected by dimensionality. However, the F1 score and normalized MCC decreased with increasing *p* for most methods (Figure S16). *GLassoElnetFast* consistently outperformed all other PMEMs across binary classification metrics and was the least affected by increasing dimensionality. Matrix norm values and Kullback-Leibler losses increased with higher dimensionality for nearly all PMEMs (Figures S17 and S18). *GLassoElnetFast* mostly achieved the lowest values in all matrix norms and KL losses across the full range of dimensions. Occasionally, lower norm values were observed for *bigquic, scio*, and *tiger*. However, these methods again estimated empty precision matrices in all settings. Their diagonal values ranged from 0.26 to 1.23 for *tiger*, 0.26 to 1.21 for *scio*, and were constantly at 0.5 for *bigquic*. Differential edge recovery rates decreased with increasing dimensionality for most PMEMs. *GLassoElnetFast* again achieved the highest recovery rates across all settings, although differences between methods became less distinct at higher dimensions (Figure 4).

**Figure 4:**
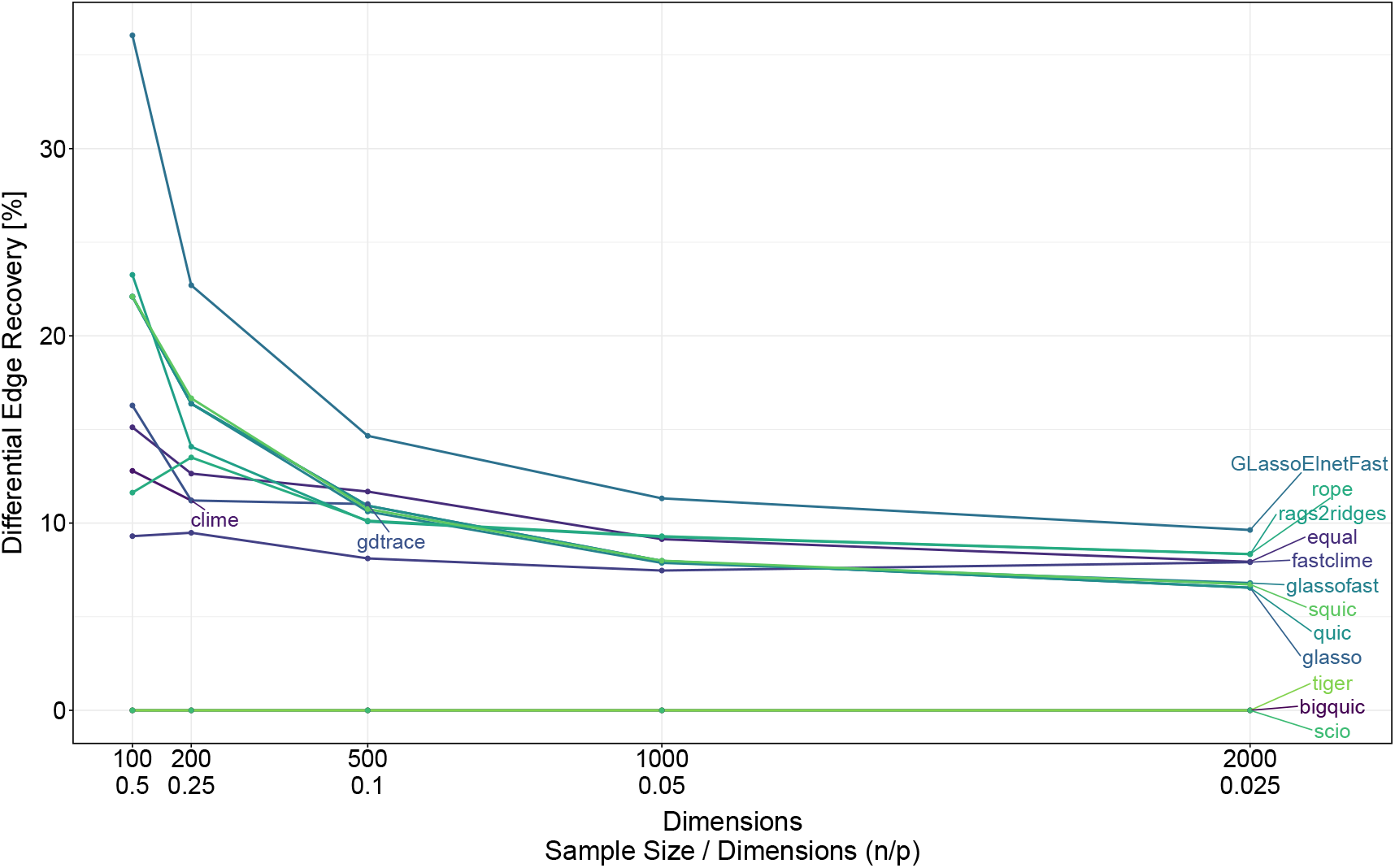
Line plot illustrating the impact of increasing the dimensions on the performances of precision matrix estimation methods, assessed by differential edge recovery rates. The PMEMs *scio, tiger*, and *bigquic* did not recover any differential edges across the considered dimensions. Differential edge recovery rates decreased with increasing dimensionality for the remaining PMEMs. However, *GLassoElnetFast* achieved the highest recovery rates across all settings, although the differences between PMEMs became less distinct at higher dimensions.

### 4.7 Mean Distribution

Next, the effect of shifting the mean values of the variable distributions in the simulated datasets was evaluated. While the covariance structure remained constant, the mean vectors were either set to zero, fixed at ten, or randomly sampled within the range of five to fifteen. Results are shown in Supplementary Figures 19, 20, and 21. Overall, no visible impact on performance was observed for any PMEM across the binary classification metrics, matrix norms, or Kullback-Leibler losses.

### 4.8 Normalization

In this approach, we assessed the impact of two normalization strategies using datasets, where variable means were randomly sampled in the range of five to fifteen, as in the previous analysis. Normalization strategy 1 standardizes the variable distributions to have both zero mean and unit variance. In the second normalization approach, variables are mean-centered only, retaining their original variances. The resulting performance differences are summarized in Supplementary Figures 22, 23, and 24. Binary classification metrics reveal a slight performance decrease for *GLassoElnet-Fast* when applying normalization strategy 1, while no notable differences are observed between the non-normalized data and normalization strategy 2 for any PMEM (Figure S22). The matrix norm values increase across nearly all PMEMs when applying normalization 1, indicating reduced performance. Only *scio* and *tiger* show decreased matrix norm values, suggesting improved estimates under this strategy. In contrast, normalization 2 has no visible impact on any of the matrix norms (Figure S23). Kullback-Leibler losses increase for all PMEMs under normalization 1, while reverse Kullback-Leibler losses tend to decrease. Again, normalization 2 does not influence performance (Figure S24).

### 4.9 Sample Size

This section examines how the sample size or *n/p* ratio influences the performance of different PMEMs. This was done by increasing the sample size while keeping the dimensionality fixed, thus increasing the *n/p* ratio (Figure S25). The results are presented in Figure 5 and Figures S26-S30. The F1 score and normalized MCC improved for most PMEMs as the sample size increased. *Equal* and *gdtrace* showed the strongest gains, outperforming all other methods in F1 score, accuracy, and normalized MCC as the *n/p* ratio increased. For *tiger, scio, rope, rags2ridges, bigquic*, and *clime* the F1 score remained constant at 0.18. The accuracy of all methods remained largely unchanged by different *n/p* ratios. The normalized MCC values for *tiger, scio*, and *bigquic* stayed constant at 0.65, and for *clime, rags2ridges*, and *rope* at 0.5. Matrix norm values generally decreased with increasing sample size, though high fluctuations were observed across most PMEMs. In contrast, the norm values increased for *rags2ridges*, reaching an matrix 1-norm of 37.5, Frobenius norm of 103, and spectral norm of 12.5 at *n* = 100. *GLassoElnetFast* initially showed decreasing norm values with larger sample sizes, but these increased again at *n >* 60 (Figures S27, S28). Kullback-Leibler losses followed a similar trend, decreasing for most PMEMs as sample size increased. *GLassoElnetFast* showed an initial decrease until *n* = 85, followed by an increase at higher sample sizes. As before, unlike all other methods, *rags2ridges* exhibited increasing KL losses. These losses reached values as high as 876 for KLL and 150 for reverse KLL at n = 100 (Figures S29 and S30). Differential edge recovery improved with increasing sample size for almost all PMEMs (Figure 5). Despite their strong performance in binary classification metrics, *equal* and *gdtrace* showed poor results in the differential edge recovery metric, with recovery rates among the lowest at *p* = 100. *GLassoElnetFast* and *rags2ridges* achieved the highest recovery rates. Overall, matrix norm values were not well-aligned with edge recovery performance in this setting. *Rags2ridges*, despite extremely high norm values, and *GLassoElnetFast*, with average norm values, both performed well in recovering the underlying differential structure.

**Figure 5:**
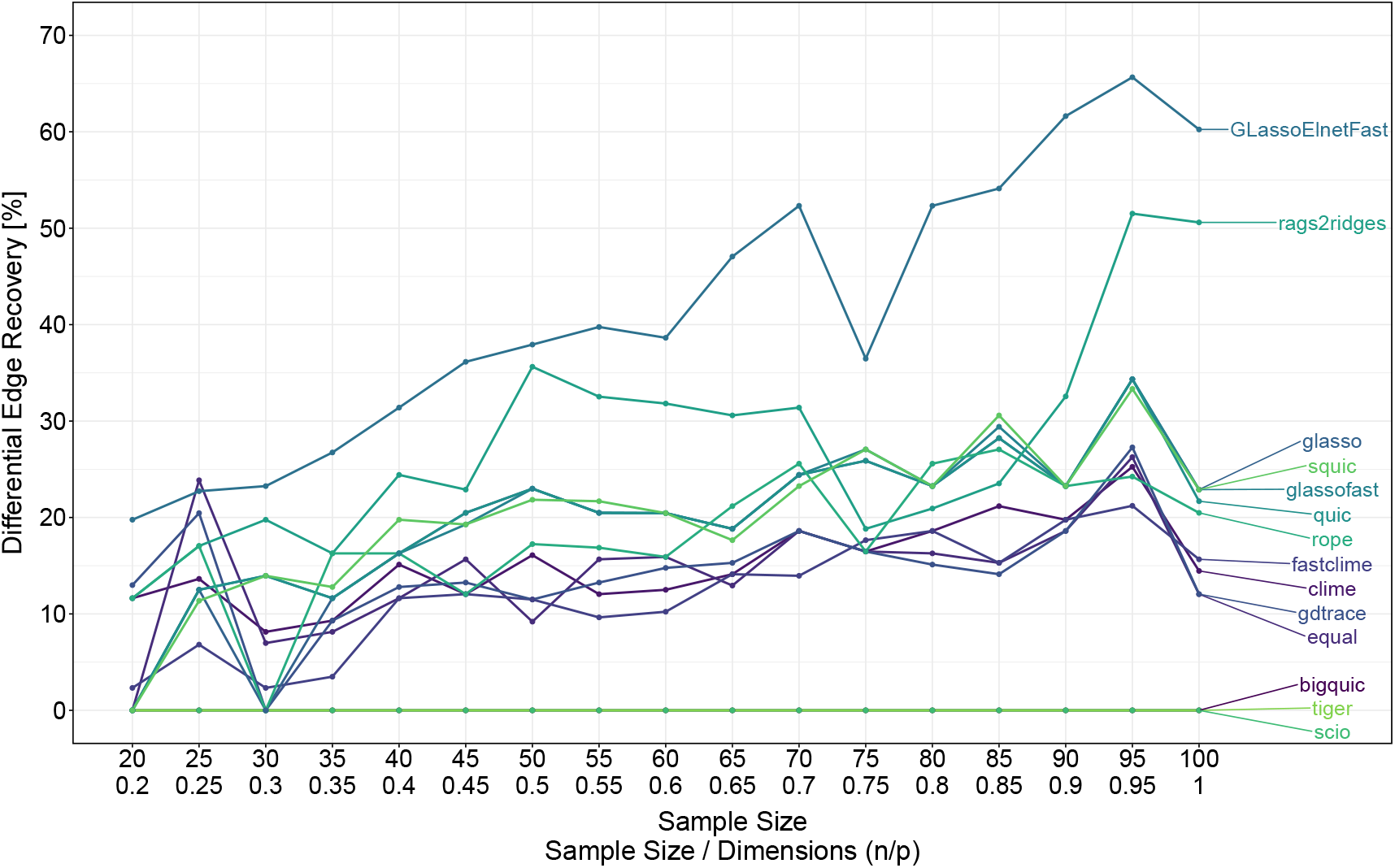
Line plot illustrating the influence of increasing sample size–to–dimension ratios on the performance of precision matrix estimation methods, assessed by differential edge recovery rates. The PMEMs *scio, tiger*, and *bigquic* did not recover any differential edges across the considered ratios. All remaining methods exhibited increasing differential edge recovery rates as the sample size–to–dimension ratio was increased. Comparable recovery patterns were observed for *fastclime, clime, equal*, and *gdtrace*, which showed similar, steadily increasing recovery rates. Slightly higher recovery rates, with otherwise comparable trends, were observed for *glasso, squic, glassofast*, and *rope. Rags2ridges* generally achieved higher recovery rates than these methods across ratios. Among all PMEMs, *GLassoElnetFast* consistently outperformed the other approaches in terms of differential edge recovery.

### 4.10 Runtimes

In addition to evaluating PMEM performance across different scenarios, estimation runtimes were recorded for each method in every approach. A representative subset of these results stems from the dimension increase scenario, in which PMEMs were applied to datasets of increasing dimensionality (*p* = 100, 200, 500, 1000, and 2000). Figure 6 visualizes the corresponding runtimes. It should be noted that *clime* and *gdtrace* were only included up to *p* = 200 and *p* = 500, respectively, due to their substantially longer runtimes compared to the other methods. Overall, runtime progression was consistent across dimensions, with the relative ranking of PMEMs from slowest to fastest remaining largely unchanged. At *p* = 2000, *scio* achieved the shortest runtime, followed by *rope, glassofast*, and *equal. glassofast* and *fastclime* showed clearly reduced runtimes compared to their original counterparts, *glasso* and *clime*, respectively, confirming their intended runtime advantages. Similarly, *squic* consistently outperformed *quic* up to *p* = 1000, but at *p* = 2000 it demonstrated the longest runtime among the remaining PMEMs, exceeding 1,710,000 seconds, or roughly 20 days. Finally, estimation times were highly consistent across replicates for all methods, with the largest variation observed in *quic* (Figure S31).

**Figure 6:**
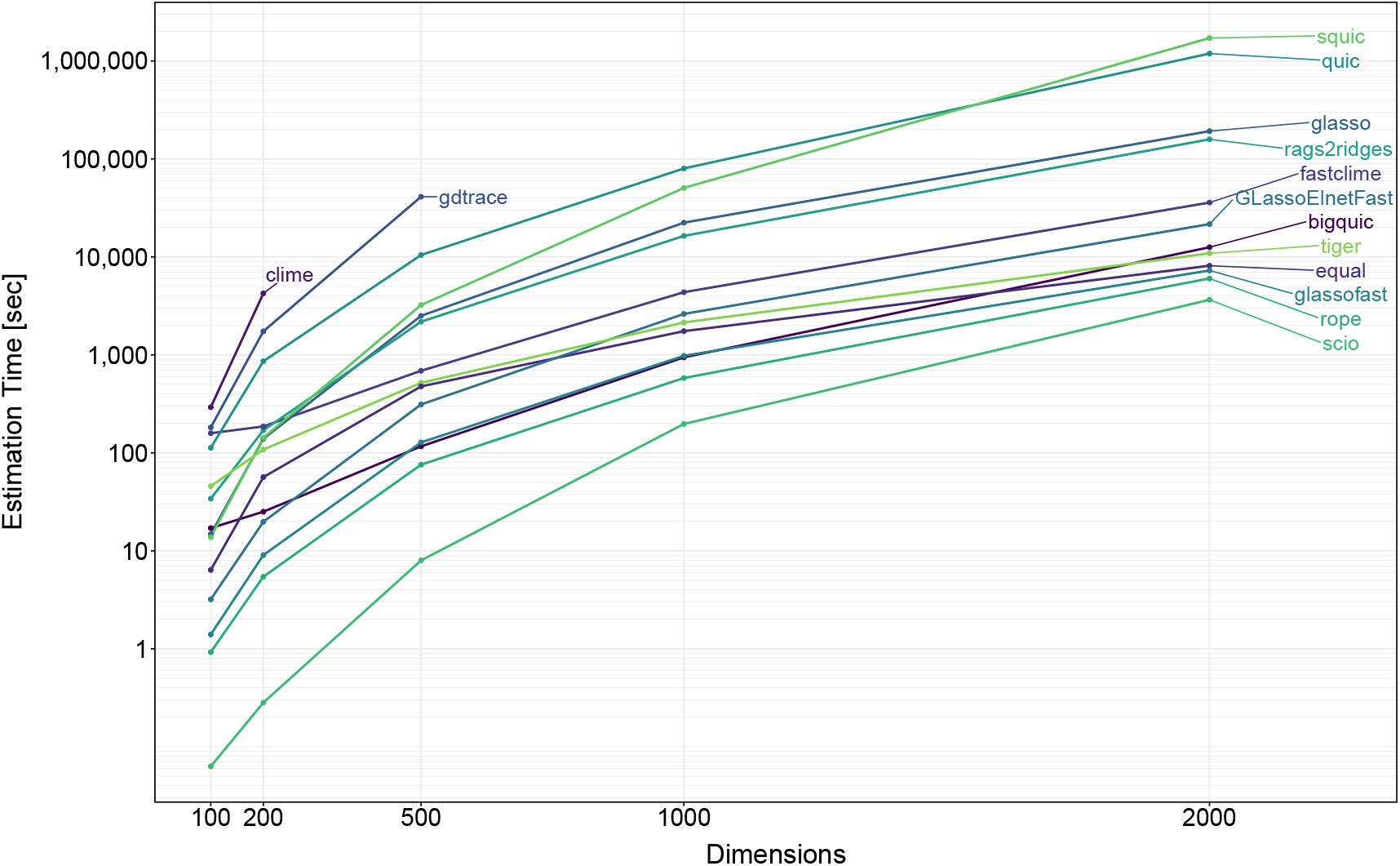
Line plot illustrating the impact of increasing the dimensions on the estimation times of precision matrix estimation methods.

### 4.11 Sampling Distributions

Until this point, all simulated datasets were generated using Gaussian-distributed samples, typically with a zero mean vector. While this reflects the assumptions underlying most estimation methods, RNA-seq data is count-based and, without transformation, is often more appropriately described by a Poisson distribution. This approach therefore examined how different sampling distributions affect PMEM performance. Datasets were generated from predefined covariance matrices using two Gaussian-based methods (mvrnorm and rmvnorm) and three Poisson-based methods (PoisNor, PoisBinOrdNonNor, and the approach proposed by Yahav and Shmueli [75]). The resulting distributions are shown in Supplementary Figure 32. The performance results are presented in Figures S33-S35. Binary classification metrics were largely unaffected by the sampling method, except for *GLassoElnetFast*, which showed decreased F1 score, accuracy, and normalized MCC when applied to Poisson-sampled datasets. Matrix norm values generally increased when moving from Gaussian to Poisson-based sampling. Some PMEMs, however, achieved slightly better norm values under Poisson-based conditions. This pattern was also observed for the Kullback-Leibler losses. Overall, Poisson-distributed data impacted PMEM performance in terms of KullbackLeibler losses and matrix norms, while different methods within each distribution type produced comparable results.

## 5 Discussion

In this work, a simulation pipeline was developed that enables the systematic comparison of a large number of precision matrix estimation methods. Both established methods, which often serve as a basis for comparison, and newer methods were analysed and compared with each other in a variety of simulation approaches. The precision matrix estimation methods were applied under consistent, untuned conditions and were not specifically adjusted to the different simulation settings. Consequently, the results presented here reflect a lower-bound estimate of each method’s performance. Parameter tuning was performed in each scenario, but the setup for this tuning, such as the range of regularization parameters and the cross-validation loss function, was defined once during the development of the simulation pipeline and not modified afterwards. This was mainly done to avoid overfitting the methods to individual scenarios. Furthermore, this also better mimics a case in which the ground truth is not known, such as when using real gene expression data. However, this approach also meant that the best performances of the methods may not have been observed, which might otherwise have been achieved with other settings, such as a different selection of regularization parameters or adapted parameter tuning methods.

Overall, the PMEM performance strongly depended on the simulation approaches, particularly on variations in individual parameters. In addition, the performance of the PMEMs could not be determined solely on the basis of individual evaluation metrics. This large number of influencing factors complicates the evaluation of the PMEMs. Accordingly, narrow simulation strategies, which are frequently used in the publication of new methods, are prone to consider the performance of the PMEMs only one-sidedly and can therefore be misleading. This also has a particular impact on the comparability of results between different publications. By incorporating various approaches and evaluation metrics, this study enabled a more objective evaluation of the analysed PMEMs. It was observed that the densities of the estimated matrices did not change over the different densities of the ground truth matrices. The exception was *GLassoElnetFast*, which always slightly overestimated the density, but more importantly resulted in an adjusted density of Θ as the density of Θ increased. This points to inherent weaknesses of both *ℓ*_1_- and *ℓ*_2_-regularized PMEMs. The *ℓ*_1_ regularization introduces a structural bias towards sparse solutions. This is often preferred, since zero partial correlations directly impose interpretable graph structures [50]. However, this potentially introduces an issue, if, for example, the networks are estimated too sparse, clusters or hub nodes in the estimated network may remain undetected. These hub nodes, in particular, could hold significant biological relevance [69]. This problem is only amplified when the precision matrices or partial correlation networks of two conditions are estimated to sparse and the differential edges are to be identified. This was observed, for example, with the PMEMs *equal, fastclime*, and *gdtrace*, which usually estimated the matrix too sparsely and also exhibited below-average differential edge recovery. Even if other regularization parameters can lead to denser estimates, the respective parameter tuning method only selects the parameters that, for example, reduce the likelihood loss, regardless of the density of the matrices. As for the *ℓ*_2_-regularized PMEMs, or overall the PMEMs that estimate dense matrices, *rags2ridges* demonstrated the poorest performance in terms of the matrix norms of the unfiltered estimated matrices. Furthermore, the F1 score and accuracy fail to evaluate the real performance if 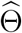 is fully dense. The MCC identifies these as random. Nevertheless, *rags2ridges* outperformed all other PMEMs, except *GLassoElnetFast*, in most simulation approaches in the differential edge recovery metric. This means that neither the matrix norms nor the binary classification metrics values provide any direct indication of the PMEMs’ performance in differential network analysis. It also became evident that estimating a dense matrix alone is not sufficient if the weights of the individual edges are not accurately recovered. Methods such as *rope* and *clime* generally performed similar to the sparse estimators in terms of differential edge recovery. In contrast, *rags2ridges* and, most notably, *GLassoElnetFast* were the only ones that consistently stood out with notably better edge recovery. However, in a real world scenario, a post-hoc edge selection also introduces a further source of error if, for example, too many edges are deleted.

The simulated analyses presented here demonstrate that the performance of the PMEMs is substantially impacted by most of the analysed parameters. Notably, covariance structure, matrix density and sample size had a substantial impact on estimator performance. This showed that previous less extensive evaluations ran the risk of testing and evaluating the PMEMs too selectively. This resulted in misleading, non-comparable and non-reproducible findings. Therefore, it is not surprising that the previous evaluation studies in the field have not been reproduced, given the influence of numerous factors, including covariance structure, density, covariance values, sample size-to-dimension ratio, sampling distribution, normalization, and the choice of regularization parameters. A comprehensive evaluation, that systematically varies all relevant parameters and incorporates diverse evaluation metrics, is essential for a robust comparison of precision matrix estimation methods. Overall, many weaknesses of the various PMEMs were identified, such as excessive regularization, scaling problems and low consistency. Notably, some PMEMs are only poorly documented, further limiting their usefulness. In the differential network analysis, *GLassoElnetFast* was clearly identified as the most accurate and effective method to identify the simulated differences between the artificial conditions. Additionally, *GLassoElnetFast* benefited most from optimal conditions, such as a high signal-to-noise ratio and a high sample-to-dimension ratio.

As a separate observation, the Poisson method proposed by Yahav and Shmueli [75] is recommended for future simulations, as it produces results comparable to the other Poisson-based approaches while being significantly faster. The method is described in the Supplementary Information (Section 11), and runtime comparisons are provided in Figure S36.

## 6 Potential implications

Precision matrix estimation is commonly applied to gene expression datasets to infer underlying network structures [69, 76–79]. We show that different estimators vary substantially both in their overall performance and in their sensitivity to data properties. If similar variability occurs in empirical analyses, the choice of estimator may strongly influence the resulting networks, with the risk of missing biologically relevant interactions or recovering spurious ones. This reduces confidence in the stability and interpretability of real-world analyses and highlights the importance of careful method selection and benchmarking.

Within this context, the comparative analyses provide several methodological insights. Dense estimators apply global shrinkage without creating exact zeros in the precision matrix. Consequently, they generate results that are challenging to interpret and typically necessitate post-hoc edge selection to derive a meaningful network structure. Very sparse estimators, which enforce hard sparsity through zero-valued entries, can be useful for recovering single networks directly. However, their utility in differential network analysis is limited, as strong regularization risks eliminating too many true edges. Elastic net–based approaches combine the advantages of sparse and dense estimators by applying a more flexible form of regularization. This makes the inferred networks directly interpretable and suitable for differential analysis, since excessive sparsity is avoided and the risk of eliminating important edges is reduced. These results suggest that further methodological development of elastic net–based precision matrix estimation could be especially valuable. Beyond differential gene co-expression analysis, the results of this study also provide insights into how the estimation of individual precision matrices depends on factors such as covariance structure, density, and sample size. These findings may therefore be of relevance in other domains where precision matrix estimation is applied. These include single-cell transcriptomics, proteomics, and metabolomics. Beyond biology, similar challenges arise in fields like neuroscience and finance, where reliable precision matrix estimation is equally essential for uncovering underlying dependency structures.

Taken together, and within the scope of the analyses and evaluation criteria we considered, these results provide a clear practical recommendation. When sparse solutions are required, *glassofast* proves to be a reliable option. Across settings, and especially in differential network analysis, *GLassoElnetFast* represents the most robust choice overall. *Rags2ridges* may also be considered when post-hoc thresholding is acceptable.

## 7 Methods

### 7.1 Covariance Matrix Generation

We define the density of a matrix **A** ∈ ℝ^*p*×*p*^ by the proportion of non-zero entries in **A**

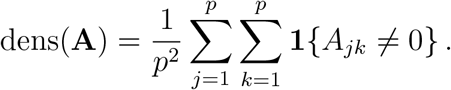

#### 7.1.1 Block Covariance Structures

The *single block* covariance method was developed based on the simulation approaches from Kuismin and Sillanpää (2017) and Fop et al. (2017) [69, 80]. With the standard settings, this approach generates a symmetric 100 × 100 matrix with one 50 × 50 sized non-zero cluster. All non-zero offdiagonal values (covariances) within this cluster are set to 1, while all diagonal values (variances) of the entire matrix are set to 2 (Figure S37A). The specified settings result in a positive definite covariance matrix with a density of 0.255. The *multiple block* covariance structure is based on the fourth simulation setting from Fop et al. (2017) [80]. The default settings result in the generation of a symmetric 100 × 100 matrix comprising ten clusters, each of size *p/*10. Similar to the single block covariance matrix, all non-zero off-diagonal values within these clusters are set to 1, while all diagonal values of the entire matrix are set to 2 (Figure S37B). The aforementioned settings result in a positive definite covariance matrix with a density of 0.1. The *band network* structure approach is based on the covariance and precision matrix generation from Liu and Wang (2012) as well as Pang et al. (2014) [81, 82]. Both utilize the band network structure to directly generate a dense precision matrix. In contrast, the band network structure is used in this work to generate a sparse covariance matrix with a density of 0.0494. Using the standard settings, the matrix is generated with diagonal values of 2. The first off-diagonal values are set to 1, and the second off-diagonal values are set to be 0.5 times the values of the first off-diagonal (Figure S37C). This guarantees a positive definite matrix.

#### 7.1.2 Scale Free Network Structures

The *scale free 1* method aims to generate a scale free network, in which the distribution of links to nodes follows a power law. For this purpose, this a network is generated using the `generate_BA()` function from the R package **PAFit** (version: 1.2.10, [83]). The parameter `num_seed` is set to 4, while the `multiple_node` parameter is set to 2. This function generates networks from the generalized Barabási-Albert model [30]. In this model, the preferential attachment function is power-law and node fitnesses are all equal to 1 (Figure S38). The generated network’s adjacency matrix is used as covariance matrix where non-zero off-diagonal values are set to 1 and diagonal values are set to 2 (Figure S39). Since the covariance matrix is generated in this way is not positive definite, the function `nearPD()` from the R Package **Matrix** (version: 1.7-0, [84]) was used to find the nearest positive definite matrix. While this approach preserves the underlying links in the network and guarantees a positive definite matrix, all edges are changed in their weighting. As a result, the matrix also loses its sparse property.

The *scale free 2* method was developed by Zhao and Shojaie (2022) to simulate changes in biological networks to test their proposed qualitative hypothesis testing framework [85]. In this approach *E*_1_ is generated from a power-law distribution with the power parameter 5 using the `sample_fitness pl()` function from the R package **igraph** (version: 2.0.3, [86]). The number of edges is chosen to be | *E*_1_ |= *p*(*p* − 1)*/*100, which corresponds to an edge density of 0.02. In the case of |*V* |≡ *p* = 200, *E*_2_ is simulated selecting 20 random nodes among the 100 most connected nodes in *G*_1_, and removing all the edges that are connected to them. Afterwards edges are added randomly to graph *G*_2_ so that |*E*_2_ | = |*E*_1_ |.

To simulate Θ_1_, for *j* ≠ *k*:

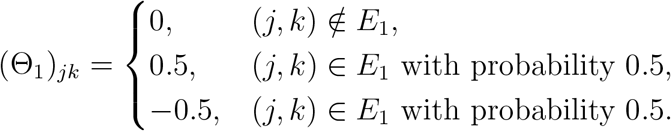

To simulate Θ_2_, for *j* ≠ *k*:

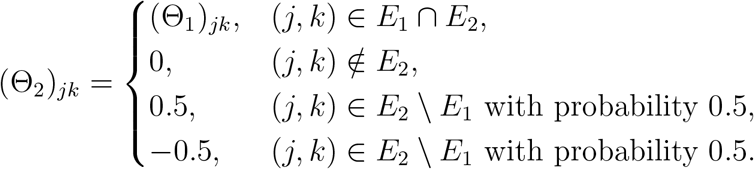

For *m* ∈ {1, 2}, the diagonal entries are defined as

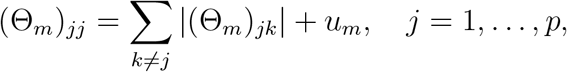

where *u*_*m*_ is chosen such that the smallest eigenvalue satisfies *ϕ*_min_(Θ_*m*_) = 0.1. Afterwards, Θ_*m*_ is normalized such that the corresponding covariance matrix 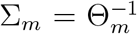 satisfies (Σ_*m*_)_*jj*_ = 1, *j* = 1, …, *p*. Examples for the generated network and covariance matrices are shown in the Supplementary Figures 40 and 41.

#### 7.1.3 Iterative Conditional Fitting Algorithm Based Structures

The following methods: *single block icf, multiple block icf, random graph icf*, and *random hubs icf* are based on Fob et al. (2018) [80] and the R package **mixggm** (version: 1.0, [87]). The included `fitGGM()` function is able to estimate a graph structure constrained GGM using the Iterative Conditional Fitting (ICF) algorithm by Chaudhuri et al. (2007) [88], ultimately yielding a positive definite covariance matrix. This approach was incorporated like the other covariance generation methods, by providing two specified adjacency matrices and a simulated mixture of *K* = 3 multivariate Gaussian distributions with mixing proportions of *τ* =(0.2, 0.5, 0.3). The means of the Gaussian distributions were randomly selected within specific ranges: (-1, 1), (-2, 2), and (-3, 3). In the *single block icf* and *multiple block icf* methods the first adjacency matrices are generated using the previously described *single block* or *multiple block* method, respectively. In the *random graph icf* method random graphs are generated according to the G(n,p) Erdős–Rényi model [89] using the `sample_gnp()` function from the R package **igraph** (version: 2.0.3, [86]) and used as the first adjacency matrix. For the *random hubs icf* method the first adjacency matrix is based on a hub structure generated by the `HubNetwork()` function from the R package **hglasso** (version: 1.3, [90]). For all these methods, the second adjacency matrix is generated using the covariance alteration method knockout. The final covariance matrices for both conditions were generated using the respective adjacency matrices and the same simulated data set (Figures S42, S43, S44, and S45).

#### 7.1.4 Covariance Alteration

In differential network analysis, the performance of precision matrix estimation methods depends on their ability to detect differences between networks across conditions. Accordingly, when simulating data from covariance matrices, these differences are introduced directly at the level of the condition-specific covariance structures. Therefore, a second covariance matrix (Σ_2_) with predefined edge-level differences relative to the first condition, based on Σ_1_, is required. To introduce controlled differences between conditions, two covariance alteration strategies were considered: *knockout* and *mutate*.

The *knockout* strategy is motivated by biological scenarios in which mutations in regulatory genes lead to the loss of downstream interactions, and is used here to model the removal of network connections between conditions. Specifically, it reflects scenarios in which mutations in a set of genes lead to structural alterations in corresponding regulatory hub proteins. These alterations affect the proteins binding properties, ultimately resulting in a loss of regulatory influence and co-expression with downstream genes. Technically, *knockout* is implemented by deleting all connections of selected nodes in Σ_2_, modifying the correlation structure without modifying the marginal distributions (Figure S46A). In line with the biological intuition, this means that the expression levels and variances of the affected genes are preserved across conditions. Analogously, the *mutate* strategy is motivated by biological scenarios involving both loss- and gain-of-function effects and is used to model simultaneous removal and introduction of network interactions between conditions. Thus, it simulates scenarios in which mutations in regulatory hub genes lead to structural changes in the corresponding proteins, resulting in both the removal of existing interactions and the formation of new ones with downstream targets. Technically, this is implemented by deleting selected edges and introducing novel connections in Σ_2_, thereby altering the correlation structure between conditions. To ensure positive definiteness of Σ_2_, the variances of the affected genes are also doubled (Figure S46B).

### 7.2 Sampling

A total of five methods were used to generate data sets for the respective simulation approaches. With a defined number of samples *n* and a given covariance matrix (Σ_1_ or Σ_2_) of the dimension *p* × *p*, these methods generate random samples from a multivariate Gaussian or Poisson distribution, thus creating a data set (**X**) with the dimension *n* × *p* for each condition. In this step the highdimensional characteristics and noise of real data sets are simulated by always selecting *p* ≥ *n*. The methods mvrnorm and rmvnorm were used to simulate random samples from a multivariate normal distribution. Additionally, data sets were generated in which the random samples are drawn from a Poisson distribution using the methods PoisNor, PoisBinOrdNonNor, and Poisson.

In the majority of the analyses conducted in this study, the mvrnorm function was utilized to generate random samples from a multivariate normal distribution. This function is based on Ripley (1987) [91] and is available in the R package **MASS** (version: 7.3-61, [92]). An example for *S* can be seen in Supplementary Figure 47. Similarly, the function `rmvnorm()` from the R package **SimDesign** (version: 2.17.1, [93, 94]) was used to generate random samples based on a normal distribution. For both of these methods, the mean value of every variable in generated data sets was set by default to zero. Alternatively, another mean value or range can be set.

In addition to drawing samples from a multivariate normal (Gaussian) distribution, a Poisson distribution was used. Three methods were utilized for this, which in the following are referred to as PoisNor, PoisBinOrdNonNor, and Poisson. PoisNor is based on the function `genPoisNor()`included in the R package **PoisNor** (version: 1.3.3, [95]). PoisBinOrdNonNor is based on the function `genPBONN()` from the R package **PoisBinOrdNonNor** (version: 1.5.3, [96]). The third sampling method, simply referred to as Poisson, is based on Yahav and Shmueli (2008) [75]. These Poisson distribution generation methods rely on a provided correlation matrix. For this, the previously generated covariance matrix was transformed into a correlation matrix using the `cov2cor()` function from the base R package **stats** [97]. Furthermore, all Poisson distribution-based sample generation methods used a rate parameter (*λ* = 0.9) for all Poisson variables. The sample generation method, previously introduced as Poisson was developed by Yahav and Shmueli (2008) [75]. A detailed description of the algorithm and implementation is provided in the Supplementary Information (Section 11).

#### 7.2.1 Normalization

Additionally after sampling, two optional normalization strategies can be applied to the generated data sets. In the first approach, variables are standardized to have both zero mean and unit variance, thereby removing differences in scale across features. In the second approach, variables are mean-centered only, retaining their original variances.

### 7.3 Precision Matrix Estimation Methods

A list of all methods included in the analysis is provided in Table 2. Methods were considered for inclusion if they (i) directly estimate the precision matrix, (ii) were described in a publication or cited in related work, (iii) provided an implementation in R as a package or through publicly available source code, and (iv) were released after *glasso* and *clime*, which have become widely used reference methods in the field. Faster implementations of earlier approaches were also included. Methods were excluded if they lacked sufficient documentation to be executed or if no working implementation was available.

#### 7.3.1 Cross-validation

To select the optimal regularization parameter for estimating the precision matrix with the PMEMs *glasso, quic, gdtrace, rope, glassofast*, and *squic*, a K-fold cross-validation procedure was employed. The process involves splitting the data set into K disjoint folds of approximately equal size. For each fold, the data in that fold is treated as a validation set, while the remaining K-1 folds constitute the training set. The simulated data set (with *n* samples and *p* variables) is partitioned into K folds, ensuring that each observation appears in the validation set exactly once. Using the training data, the sample covariance matrix (*S*_*train*_) is computed, and a precision matrix (Θ_*train*_) is estimated. The estimated precision matrix is validated using the corresponding validation data. Specifically, the sample covariance matrix for the testing set (*S*_*test*_) is computed, and the model’s performance is evaluated using the negative log-likelihood in Equation 14.

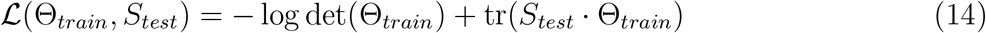

Where:

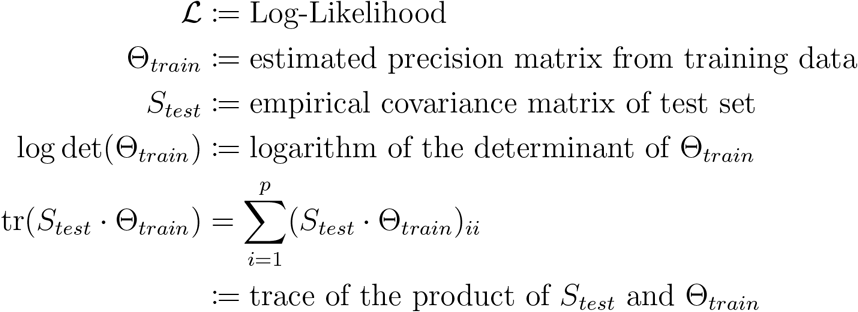

For each value of the regularization parameter (*ρ, α*), the negative log-likelihood scores are averaged across the K folds to produce a cross-validation score. This procedure is repeated for all *ρ* or *α* values inside a given list, and the parameter that minimizes the average cross-validation score is selected as the optimal regularization parameter (*ρ*_*opt*_, *α*_*opt*_).

### 7.4 Evaluation

To evaluate the performance of the precision matrix estimation methods, a set of established metrics was used, which are briefly summarized below and described in detail in the following subsections. These include matrix norms, specifically the matrix 1-norm, Frobenius norm, and spectral norm, as applied in previous studies [69, 81, 98, 99, 105–107]. Kullback-Leibler losses (KLL and reverse KLL) were also considered, consistent with their use in [69, 106, 107]. Binary classification metrics, namely F1 Score, Accuracy, and Matthews Correlation Coefficient (MCC), were used to summarize overall classification performance in identifying non-zero entries of the precision matrix, corresponding to the presence or absence of edges in the inferred network [68, 98, 106]. Additionally, a differential edge recovery metric was used to capture graph reconstruction performance, similar to approaches in [106]. In this study, the binary classification metrics (F1 Score, Accuracy, and MCC) are used to concisely summarize PMEM performance. However, the underlying evaluation pipeline also supports more detailed assessments using receiver operating characteristic (ROC) curves and Precision-Recall curves, enabling the analysis of sensitivity, specificity, precision, and recall. These metrics are likewise reported in related works [81, 98, 99, 102]. The ROC and precision–recall curves were not used in this work because they would need to be generated for every precision matrix estimation method in every simulation condition, resulting in a large number of curves that are difficult to summarize and compare. Instead, the aforementioned binary classification metrics were used to provide an interpretable summary of overall performance. ROC and precision–recall curves are nevertheless useful for detailed method-specific analyses. Two examples are shown in Figures S48 and S49.

#### 7.4.1 Matrix Norms

The following matrix norms are used as numerical benchmarks to evaluate the discrepancy between the true precision matrices Θ_1,2_ and the estimated precision matrices 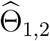. The difference matrix is defined as

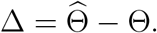

The estimation error is quantified using several matrix norms of Δ, including the matrix 1-norm, the Frobenius norm, and the spectral norm. Lower values of these norms indicate more accurate estimation performance. The matrix 1-norm (maximum absolute column sum norm) is defined for *A* ∈ ℝ^*p*×*p*^ by

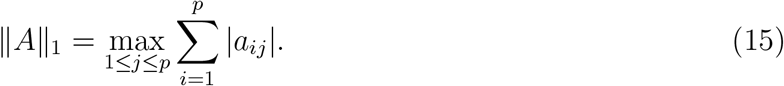

The Frobenius norm, also referred to as Schur norm, or Hilbert–Schmidt norm [109], is defined for *A* ∈ ℝ^*p*×*p*^ by

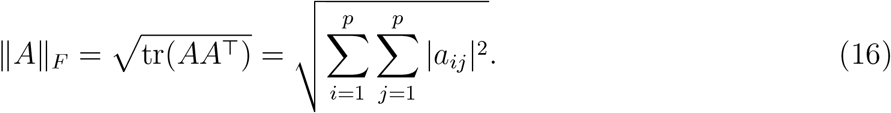

The spectral norm is defined for *A* ∈ ℝ^*p*×*p*^ as

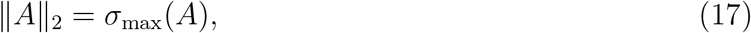

where *σ*_max_(*A*) denotes the largest singular value of *A*, and therefore represents the maximum amount by which the matrix can stretch a vector [109]. In addition to matrix norms, the estimation accuracy was evaluated using the Kullback–Leibler Loss (KLL) and the Reverse Kullback–Leibler Loss (RKLL). The Kullback–Leibler Loss is defined as

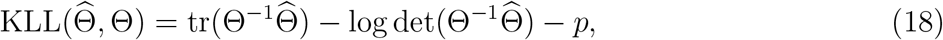

where *p* is the dimension of the matrix. The Reverse Kullback–Leibler Loss is given by

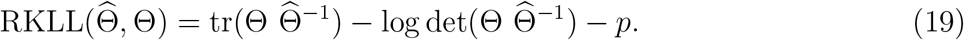

Both divergences are minimized when 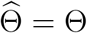, and lower values indicate more accurate estimation.

#### 7.4.2 Binary Classification Metrics

While matrix norms and divergence-based losses quantify overall estimation accuracy, they do not directly capture the ability to recover the underlying graph structure. Therefore, performance was also evaluated using the binary classification metrics F1 score, accuracy, and Matthews Correlation Coefficient (MCC), as defined in [110]. These metrics assess how well the estimated precision matrices reflect the presence or absence of edges in the true graph.

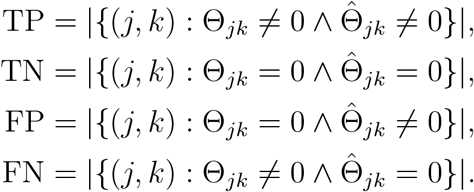

The F1 score takes values between 0, indicating the poorest performance, and 1, indicating perfect precision and recall, and is defined as:

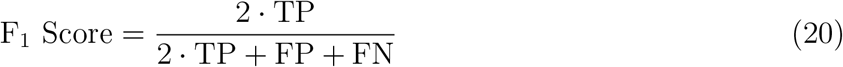

Accuracy measures the proportion of correctly classified instances among all predictions. It ranges from 0, indicating no correct classifications, to 1, indicating that all instances were classified correctly., and is defined as:

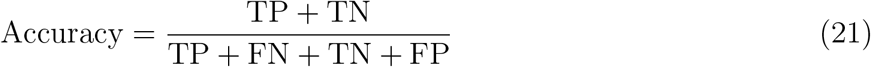

The Matthews correlation coefficient, originally proposed by B. W. Matthews (1975) [111], is a balanced measure for evaluating binary classification performance. It considers all four entries of the confusion matrix and returns a value between −1 and +1, where −1 indicates total disagreement between prediction and truth, 0 indicates random prediction, and +1 indicates perfect prediction. It is defined as:

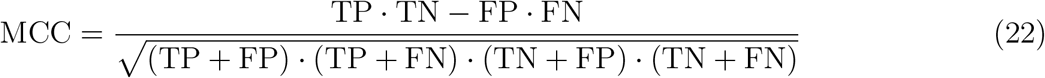

To improve interpretability, a normalized version is used, rescaling the MCC to the interval [0, 1], where 0 corresponds to the worst performance and 1 to the best:

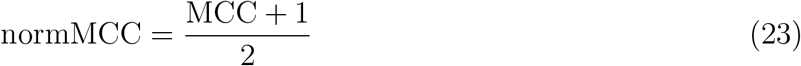

In cases where MCC is undefined due to division by zero, it is common to assign a value of 0 to MCC and 0.5 to normMCC, as justified by Chicco and Jurman (2020) [112].

#### 7.4.3 Differential Edge Recovery

To overcome the limitations of binary classification metrics, which are highly sensitive to the density of either the estimated or the true precision matrix, and of matrix norms, which do not directly assess edge recovery, we introduced a differential edge recovery (DER) metric. This metric evaluates how accurately the PMEMs capture the true structural differences between the partial correlation networks of the two simulated conditions. In addition, it provides a thresholding mechanism for PMEMs that do not inherently yield sparse matrices.

The idea of the DER score is as follows. From the ground-truth differential partial correlation matrix Ψ_Δ_, only the important differential edges are considered. Less relevant edges with very small absolute weights are excluded by applying a weight threshold *τ >* 0, which defines the filtered edge set

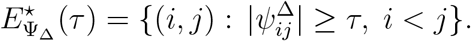

Let 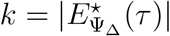 be the number of important ground-truth edges. From the estimated differential partial correlation matrix 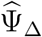, the *k* edges with the largest absolute weights are selected to form the set 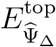. The DER score is then defined as

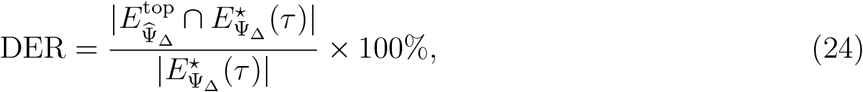

where | · | denotes set cardinality. The DER score ranges from 0% (no correct recovery) to 100% (perfect recovery).

## 8 Availability of source code and requirements

All simulations and analyses were performed on a server equipped with an AMD EPYC 7452 32-core/64-thread processor and 500 GB of RAM.

- Project name: PMEM-Evaluator
- Dockerhub: https://hub.docker.com/r/matthias587/pmem-evaluator
- Operating system: platform independent (docker)
- Programming language: R (4.4.1)
- Source and configuration files: https://github.com/scibiome/PMEM_Evaluator.git

## 9 Declarations

### 9.1 List of abbreviations and notation

The abbreviations used in this manuscript are summarized in Table 3, and the mathematical notation is given in Table 4.

**Table 2:** Overview over the used precision matrix estimation methods. Internal PT: Internal Parameter-Tuning.

| Abbreviation | Name | Publication | Year | Version | Internal PT |
| --- | --- | --- | --- | --- | --- |
| glasso | Graphical Lasso | [56] | 2008 | 1.11 | - |
| CLIME | Constrained $\ell_1$-Minimization for Inverse Matrix Estimation | [98] | 2011 | 0.5.0 | + |
| TIGER | Tuning-Insensitive Graph Estimation and Regression | [81] | 2012 | (flare 1.7.0.1) | + |
| SCIO | Sparse Column-wise Inverse Operator | [99, 100] | 2012 | 0.9.0 | + |
| GLASSOFAST | Graphical Lasso Fast | [101] | 2012 | 1.0.1 | - |
| QUIC | Quadratic Approximate Inverse Covariance | [60, 102] | 2013 | 1.1.1 | - |
| BIGQUIC | Big Quadratic Approximate Inverse Covariance | [103] | 2013 | 1.1-13 | + |
| fastclime | Fast Constrained $\ell_1$-Minimization for Inverse Matrix Estimation | [82] | 2014 | 1.4.1.1 | - |
| ROPE | Ridge-type Operator for the Precision matrix Estimation | [104] | 2017 | - | - |
| SQUIC | Sparse Quadratic Approximate Inverse Covariance | [22, 68] | 2019 | 1 | - |
| EQUAL | Efficient ADMM algorithm via the QUAdratic Loss | [105] | 2020 | 1.2 | + |
| GLassoElnetFast | Graphical Elastic Net | [106] | 2021 | 1.1.0 | + |
| GDTrace | Generalized D-Trace estimator | [107] | 2022 | - | - |
| rags2ridges | rags2ridges | [108] | 2022 | 2.2.7 | + |

**Table 3:**
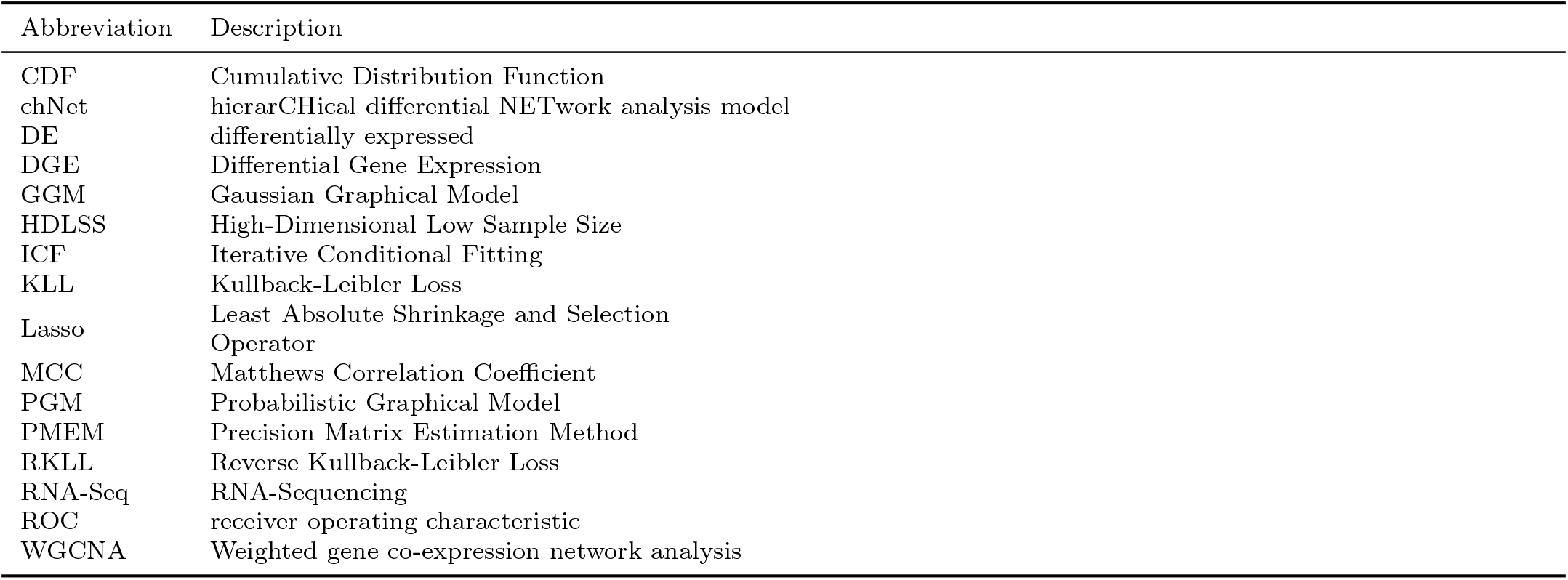
Overview over the abbreviations.

**Table 4:**
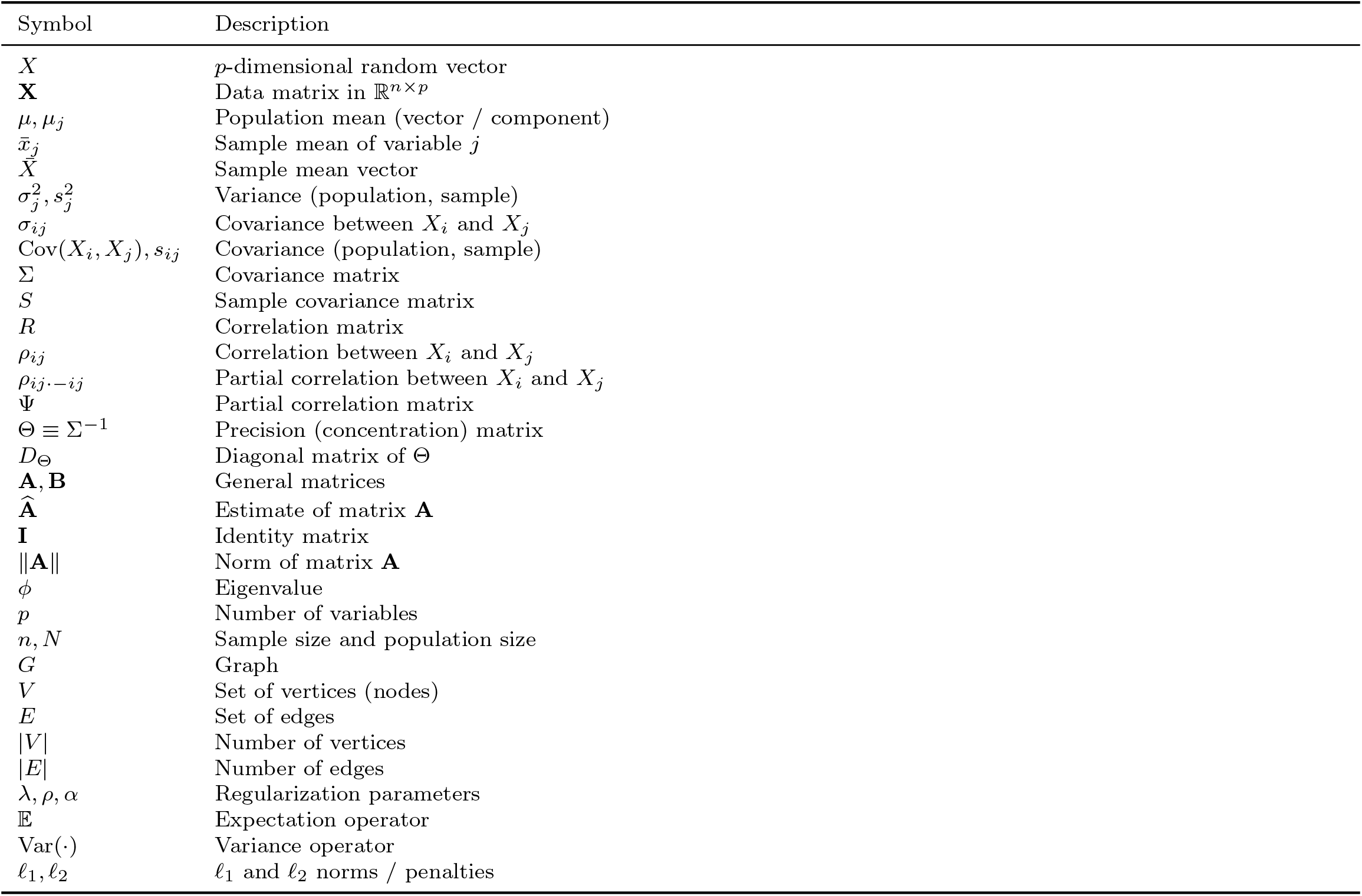
Overview of notation

### 9.2 Disclosure of use of AI-assisted tools including generative AI

AI-assisted tools (ChatGPT-4/5 and DeepL) were used solely for language support (e.g., grammar and clarity) and limited assistance in code drafting; all scientific content and analyses were developed and verified by the authors.

### 9.3 Ethical Approval

Not applicable.

#### 9.4 Consent for publication

Not applicable.

### 9.7 Author’s Contributions

**Matthias Overmann:** Conceptualization, Data curation, Formal analysis, Investigation, Methodology, Software, Visualization, Writing - original draft

**Gordon Grabert:** Conceptualization, Supervision, Validation, Writing - review & editing

**Tim Kacprowski:** Funding acquisition, Project administration, Resources, Supervision, Validation, Writing - review & editing

