## Supplementary Data for "Benchmarking precision matrix estimation methods for differential co-expression network analysis"

### Contents

1 Supplementary Figures for Replicates 5

2 Supplementary Figures for Covariance 1 Methods 6

3 Supplementary Figures for Covariance Density 9

4 Supplementary Figures for Covariance 2 Methods 13

5 Supplementary Figures for Covariance Increase 17

6 Supplementary Figures for Dimensions 20

7 Supplementary Figures for Means 23

8 Supplementary Figures for Normalization 26

9 Supplementary Figures for Sample Size 29

10 Supplementary Figures for Runtimes 35

11 Supplementary Information for Sampling 36

12 Supplementary Information for Methods 42

### 1 Supplementary Figures for Replicates

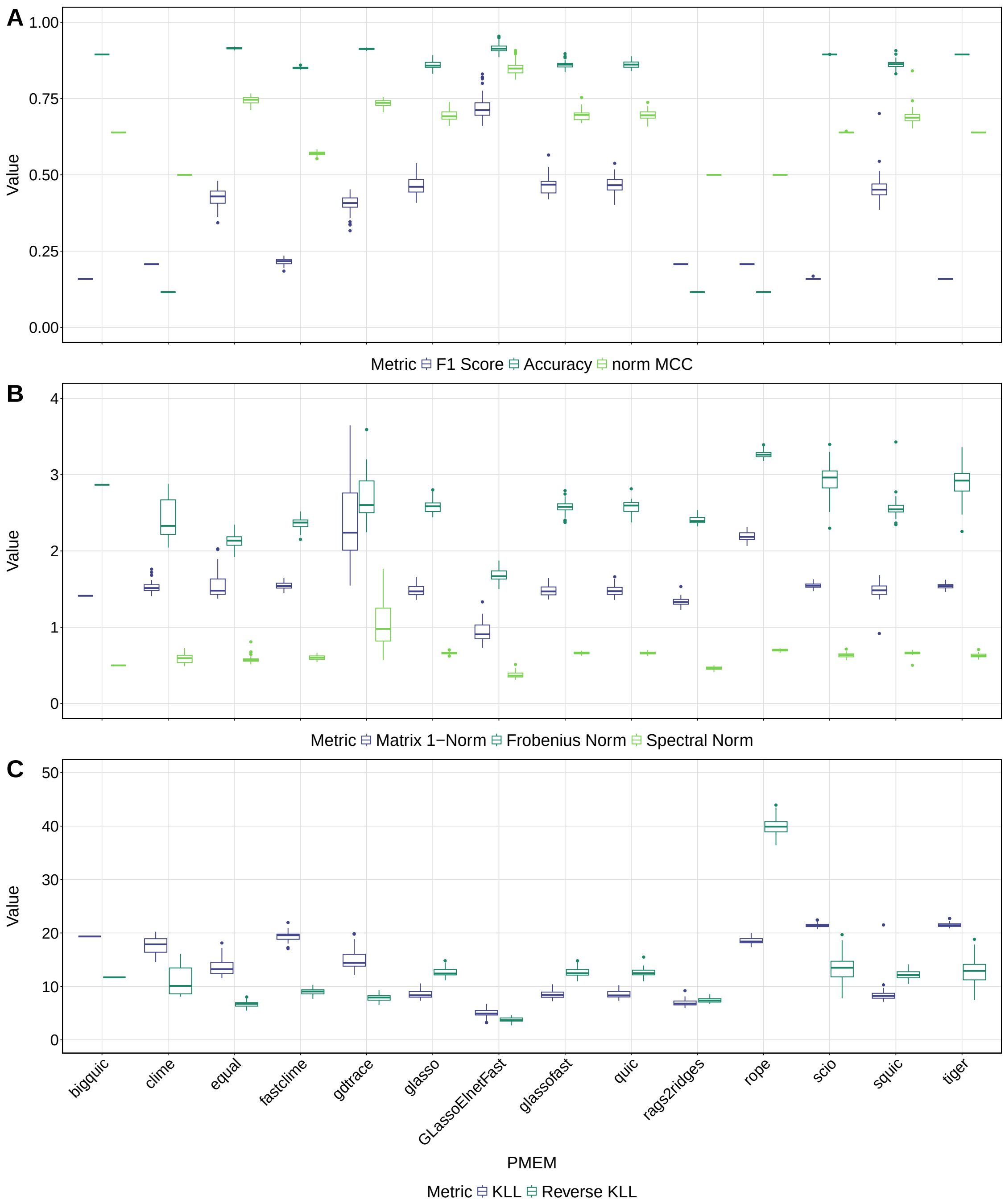

Supplementary Figure 1: Boxplot illustrating the reproducibility of different PMEMs in repeated simulations, assessed across the binary classification metrics (A): F1 score, accuracy, and normalized MCC, the matrix norms (B): matrix 1-norm, Frobenius norm, and spectral norm as well as the Kullback-Leibler Loss and the reverse Kullback-Leibler Loss (C). The performance of each method is represented by the distribution of the metric values over 50 replicates. $\Sigma_1$ was generated in each replicate with the single block method with $p = 100$. The sample data set was created using the mvrnorm sampling method with a sample size of 50. One outlier from *squic* (KLL $\approx 310$) was excluded for clarity.

### 2 Supplementary Figures for Covariance 1 Methods

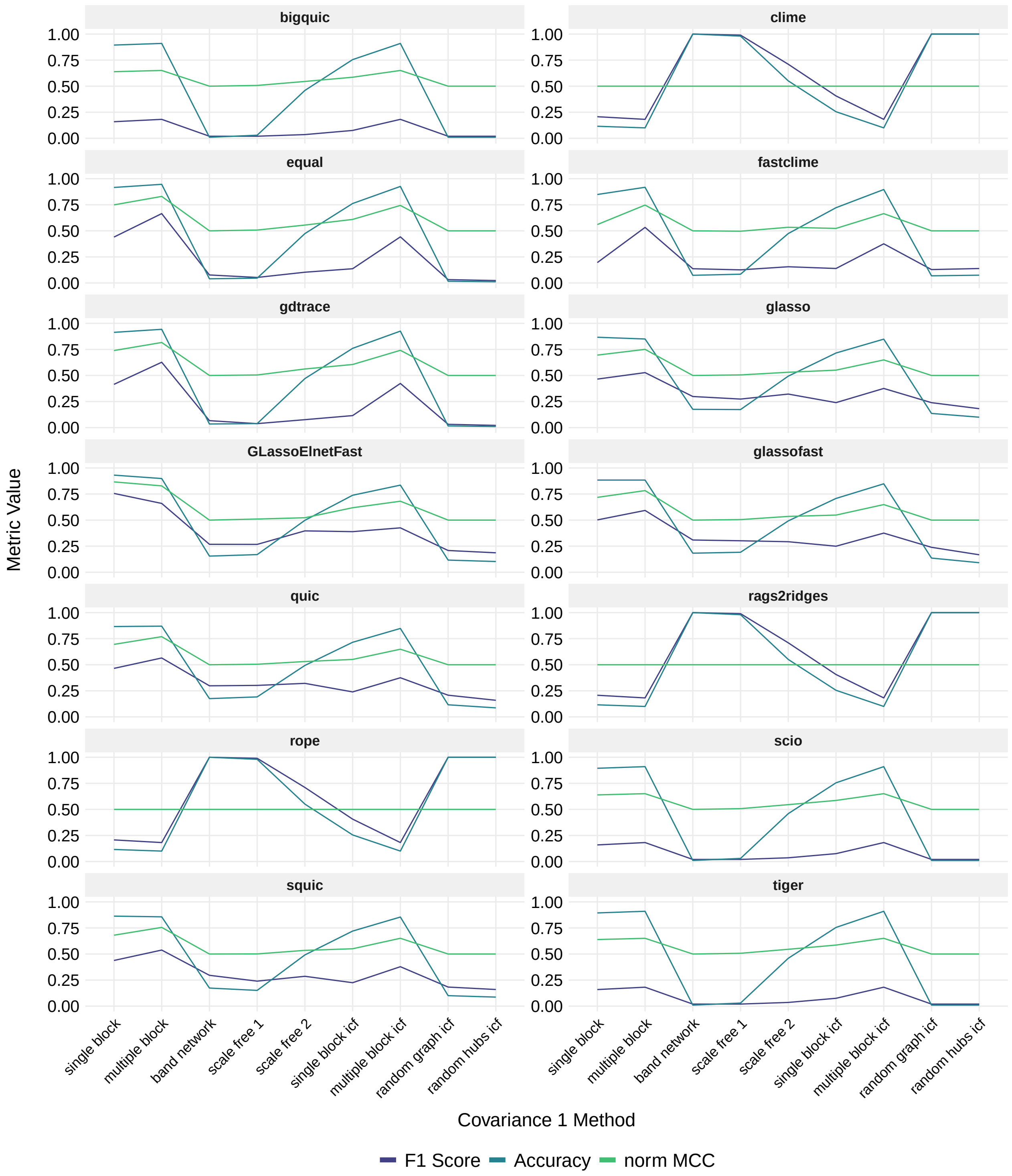

Supplementary Figure 2: Line plot illustrating the influence of different covariance 1 methods on the performances of precision matrix estimation methods, assessed across the binary classification metrics F1 score, accuracy, and normalized MCC for $\widehat{\Theta}_1$. $\Sigma_1$ was generated using the respective covariance 1 method with $p = 100$. The sample data set was created using the mvrnorm sampling method with $n = 50$.

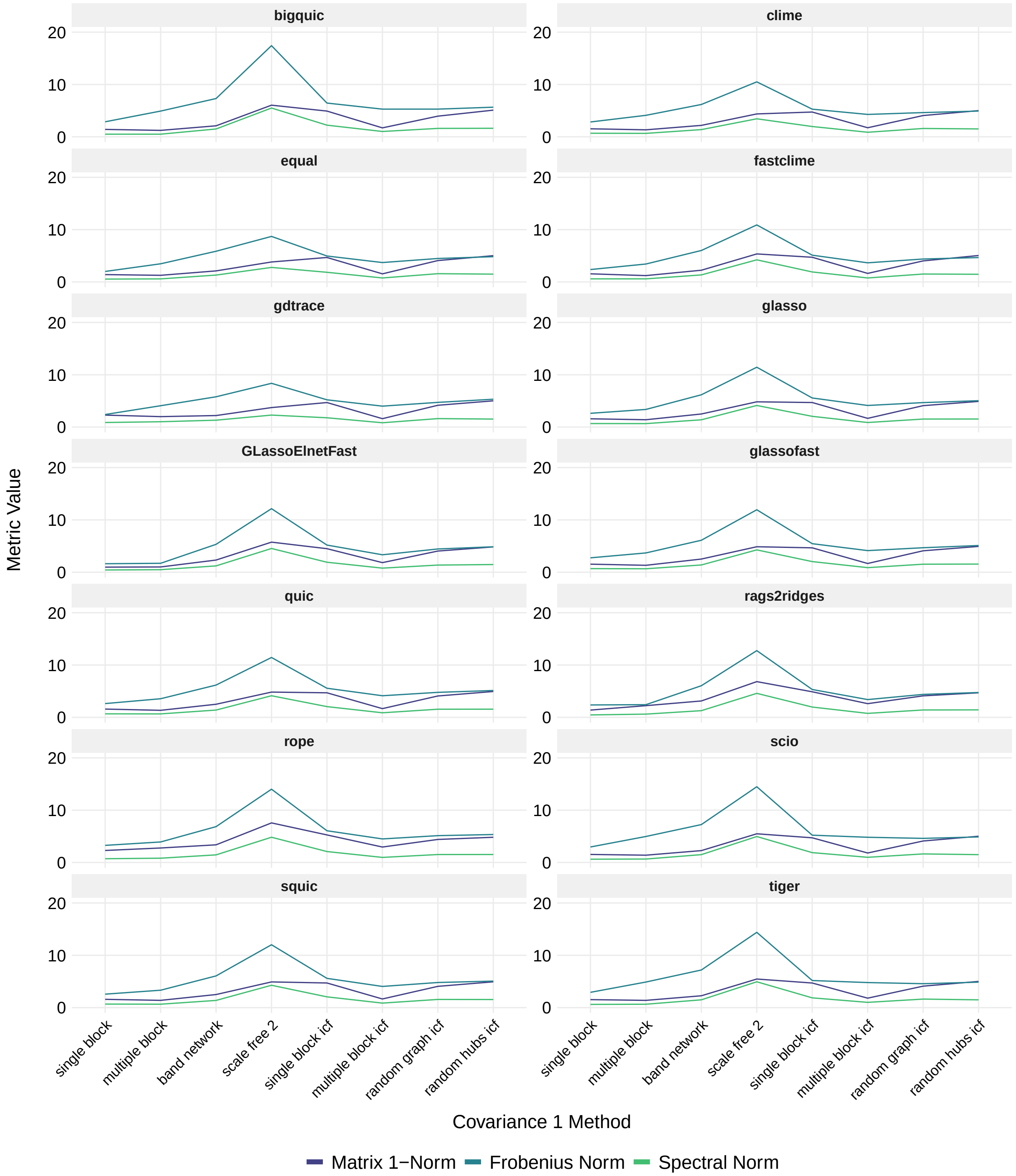

Supplementary Figure 3: Line plot illustrating the influence of different covariance 1 methods on the performances of precision matrix estimation methods, assessed across the matrix norms matrix 1-norm, Frobenius norm, and spectral norm for $\widehat{\Theta}_1$. $\Sigma_1$ was generated using the respective covariance 1 method with $p = 100$. The sample data set was created using the mvrnorm sampling method with $n = 50$. The results of the covariance 1 method scale free 1 were excluded in the figure. The matrix 1-norm was 46985054, the Frobenius norm was 33520059, and the spectral norm was 13684507 for all PMEMs in this approach.

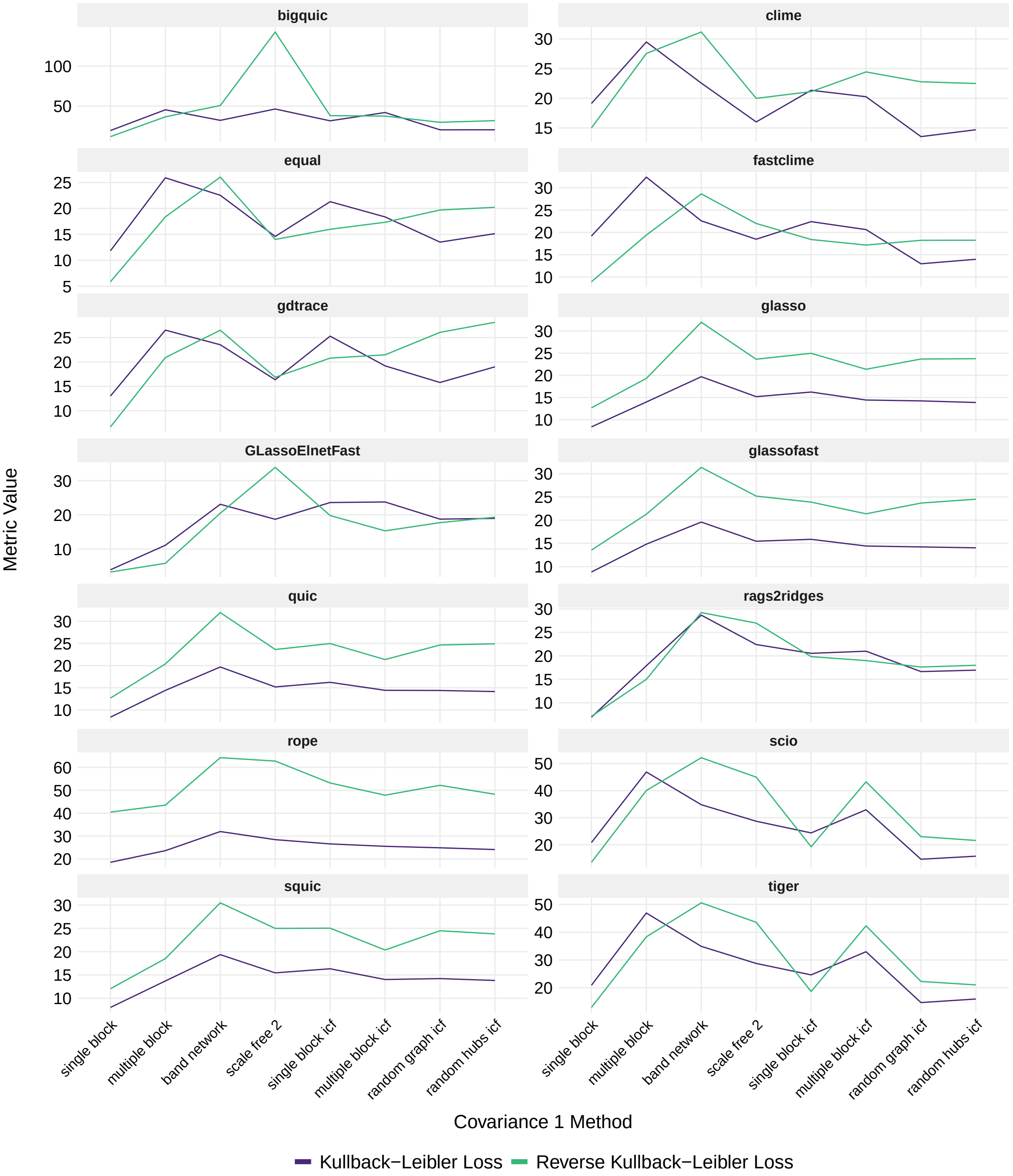

Supplementary Figure 4: Line plot illustrating the influence of different covariance 1 methods on the performances of precision matrix estimation methods, assessed across the Kullback-Leibler Loss and the reverse Kullback-Leibler Loss for $\widehat{\Theta}_1$. $\Sigma_1$ was generated using the respective covariance 1 method with $p = 100$. The sample data set was created using the mvrnorm sampling method with $n = 50$.

### 3 Supplementary Figures for Covariance Density

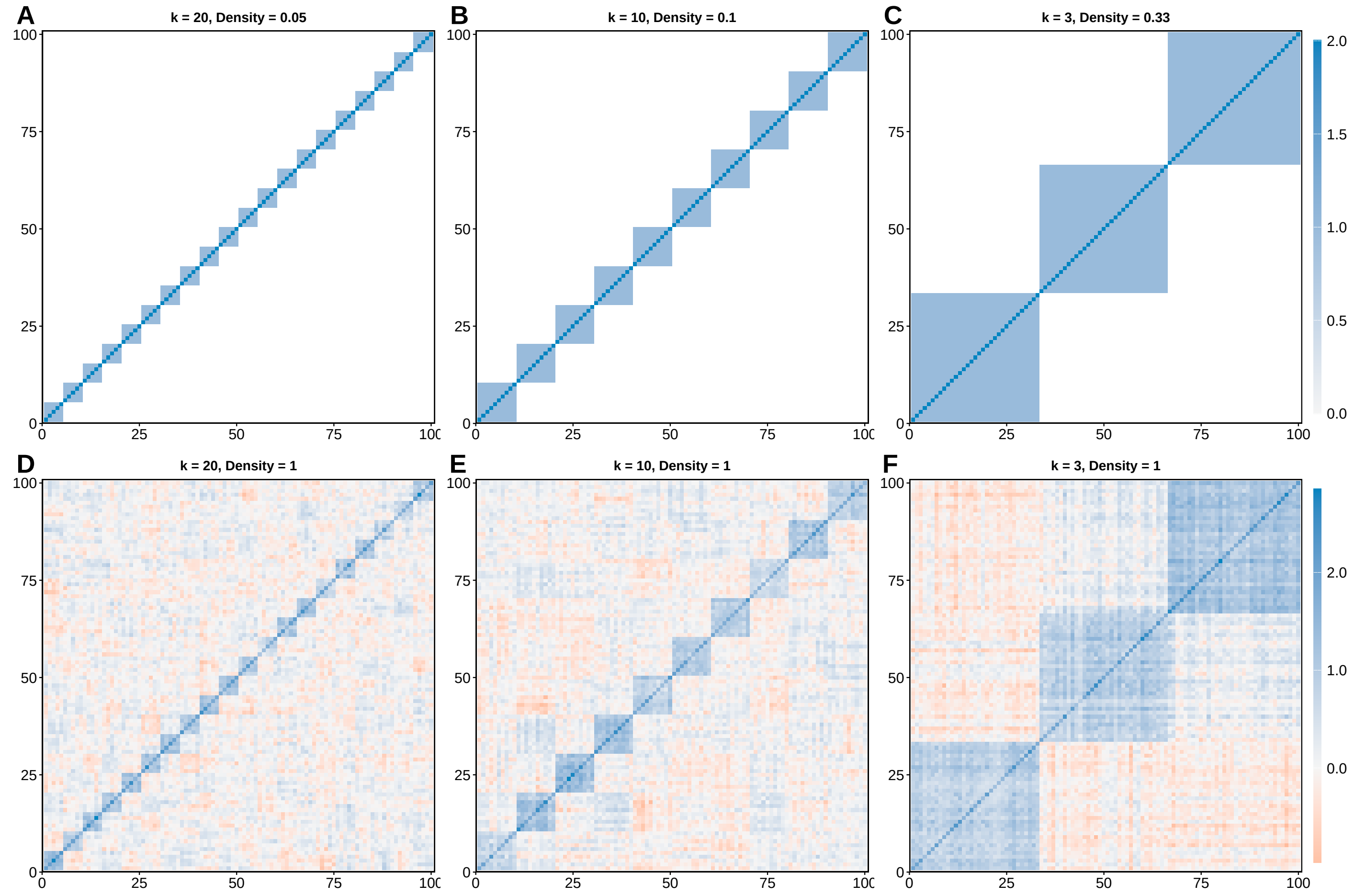

Supplementary Figure 5: A-C: $\Sigma_1$ generated using covariance 1 method multiple block with k blocks of non-zero off-diagonal values. D-F: $S_1$ generated by the mvrnorm method based on the respective $\Sigma_1$ in A-C.

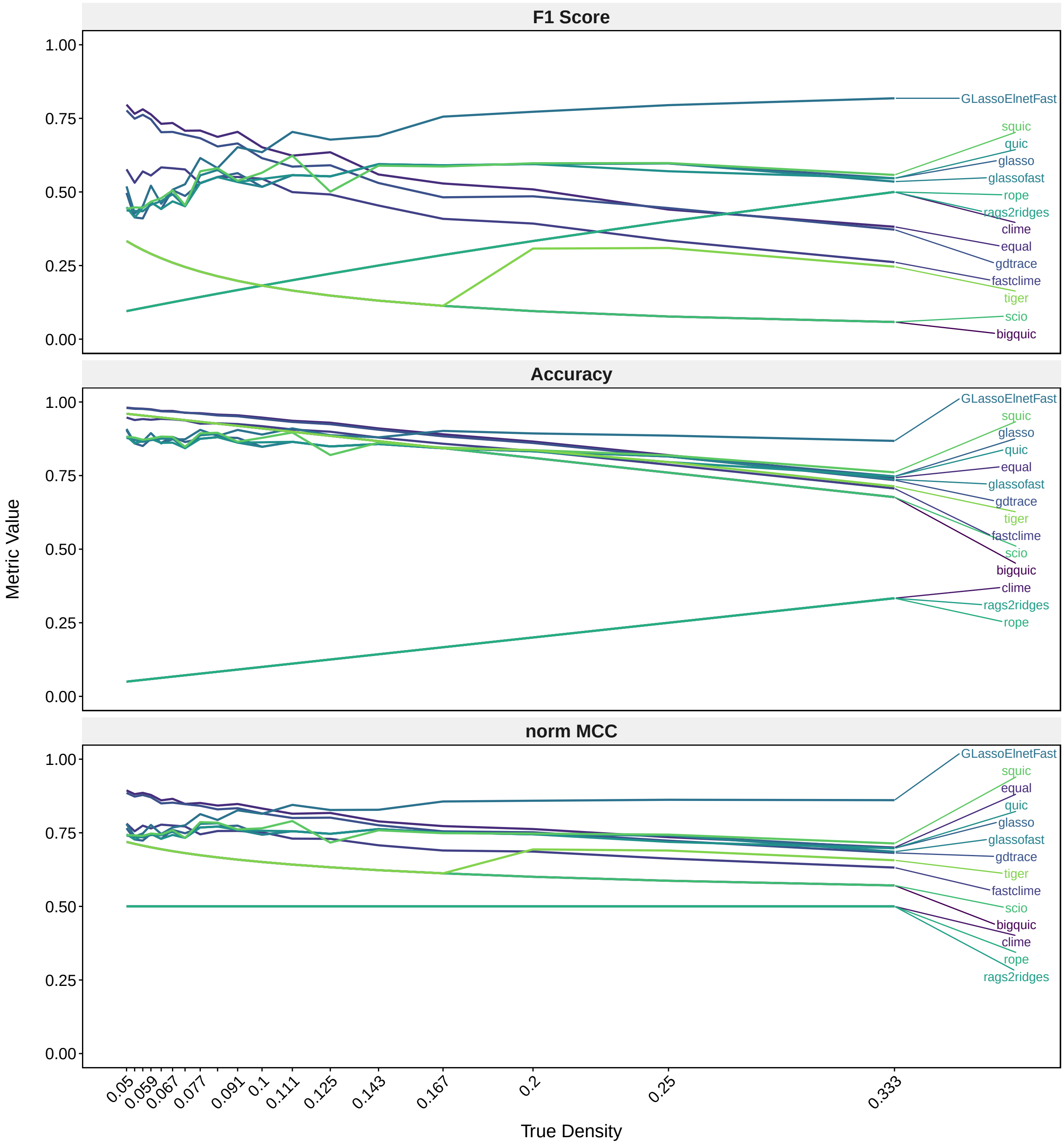

Supplementary Figure 6: Line plot illustrating the influence of different $\Theta_1$ densities on the performances of precision matrix estimation methods, assessed across the binary classification metrics F1 score, accuracy, and normalized MCC. Densities from 0.05 to 0.33 were analysed. $\Sigma_1$ was generated using the covariance 1 method multiple block with $p = 100$. The different densities for $\Theta_1$ were achieved by varying the number of non-zero blocks within $\Sigma_1$. The sample data set was created using the sampling method mvrnorm with $n = 50$.

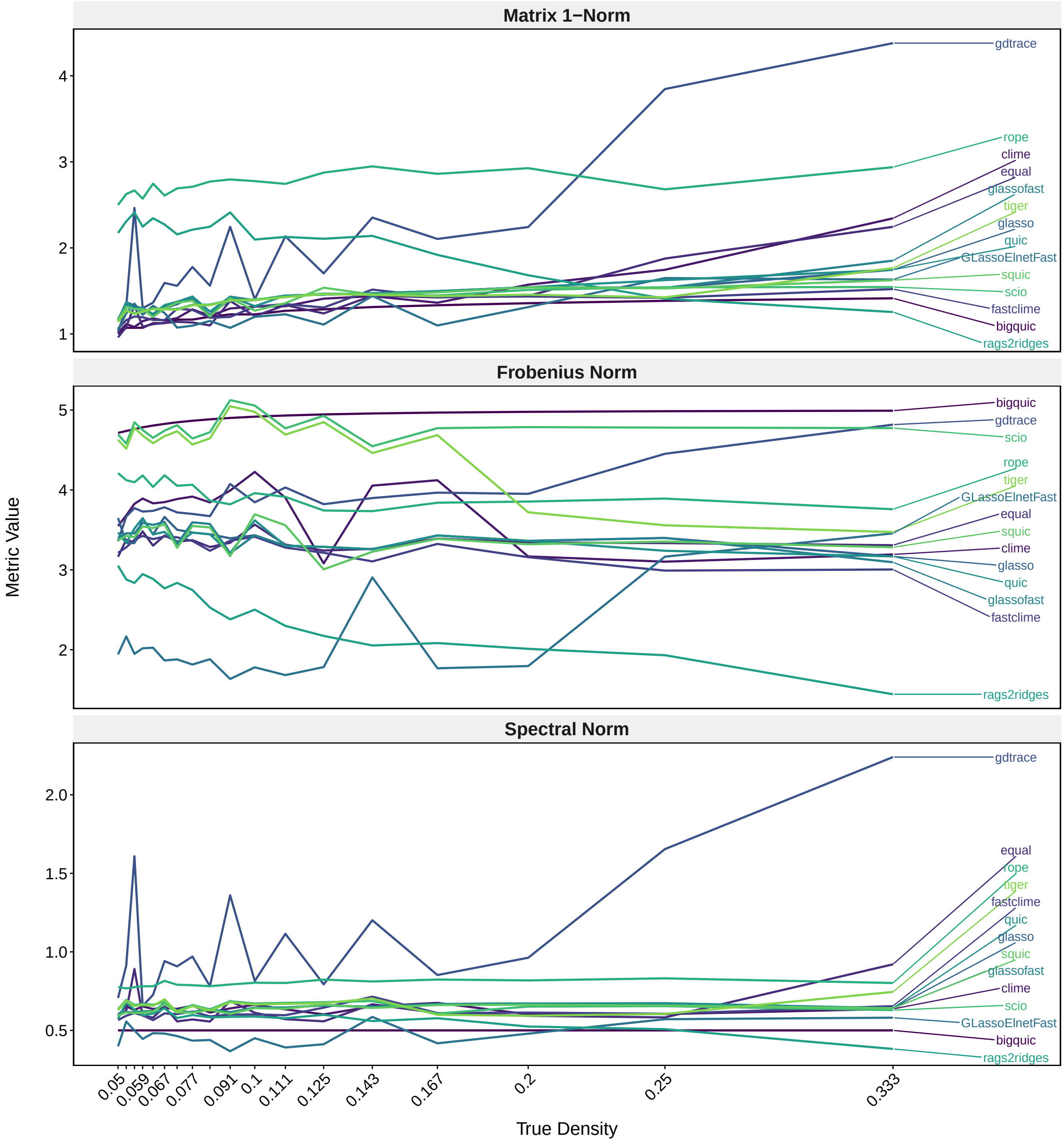

Supplementary Figure 7: Line plot illustrating the influence of different $\Theta_1$ densities on the performances of precision matrix estimation methods, assessed across the matrix norms matrix 1-norm, Frobenius norm, and spectral norm. Densities from 0.05 to 0.33 were analysed. $\Sigma_1$ was generated using the covariance 1 method multiple block, using $p = 100$. The different densities for $\Theta_1$ were achieved by varying the number of non-zero blocks within $\Sigma_1$. The sample data set was created using the sampling method mvrnorm with $n = 50$.

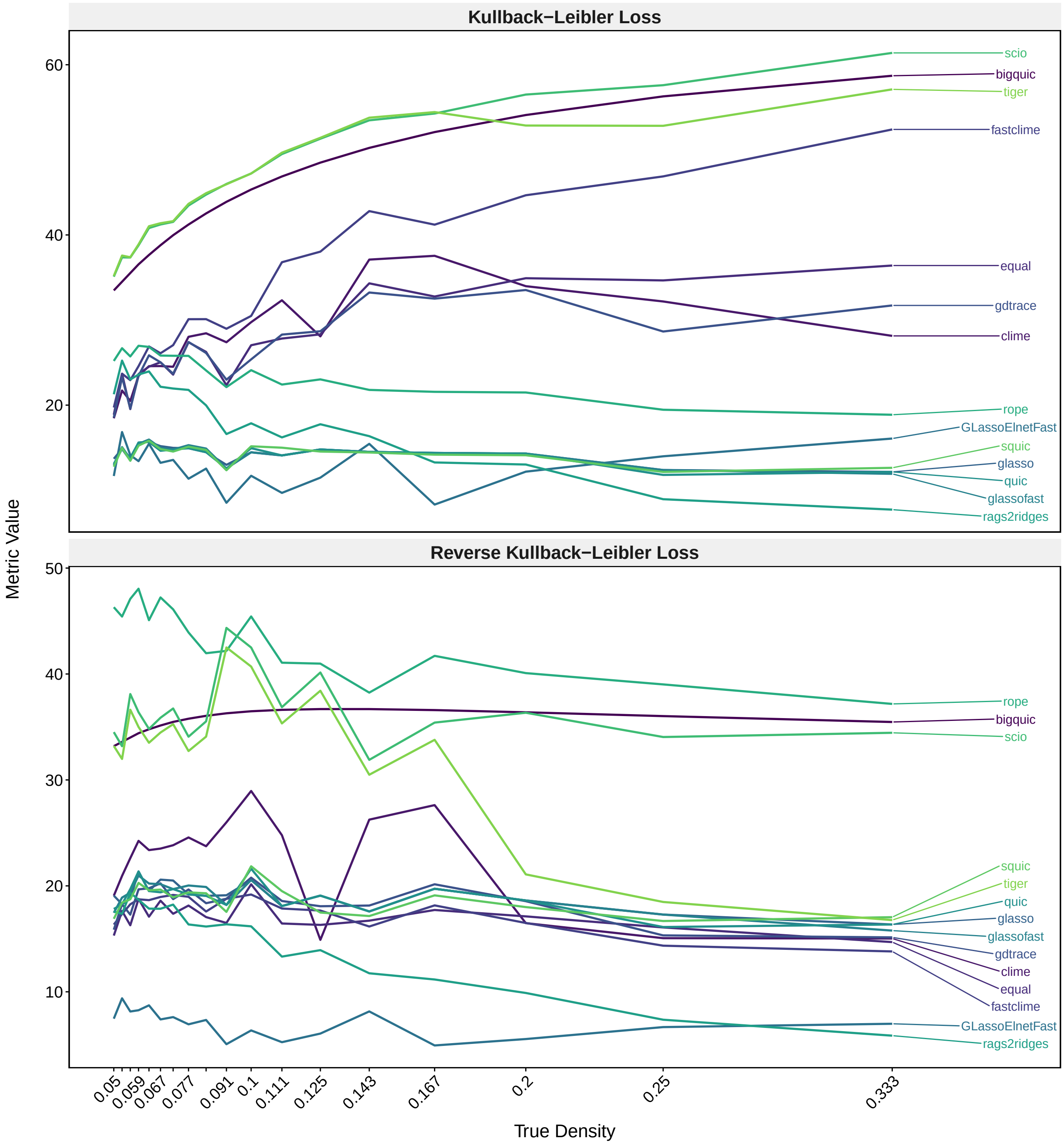

Supplementary Figure 8: Line plot illustrating the influence of different $\Theta_1$ densities on the performances of precision matrix estimation methods, assessed across the Kullback-Leibler Loss and the reverse Kullback-Leibler Loss. Densities from 0.05 to 0.33 were analysed. $\Sigma_1$ was generated using the covariance 1 method multiple block, using $p = 100$. The different densities for $\Theta_1$ were achieved by varying the number of non-zero blocks within $\Sigma_1$. The sample data set was created using the sampling method mvrnorm with $n = 50$.

### 4 Supplementary Figures for Covariance 2 Methods

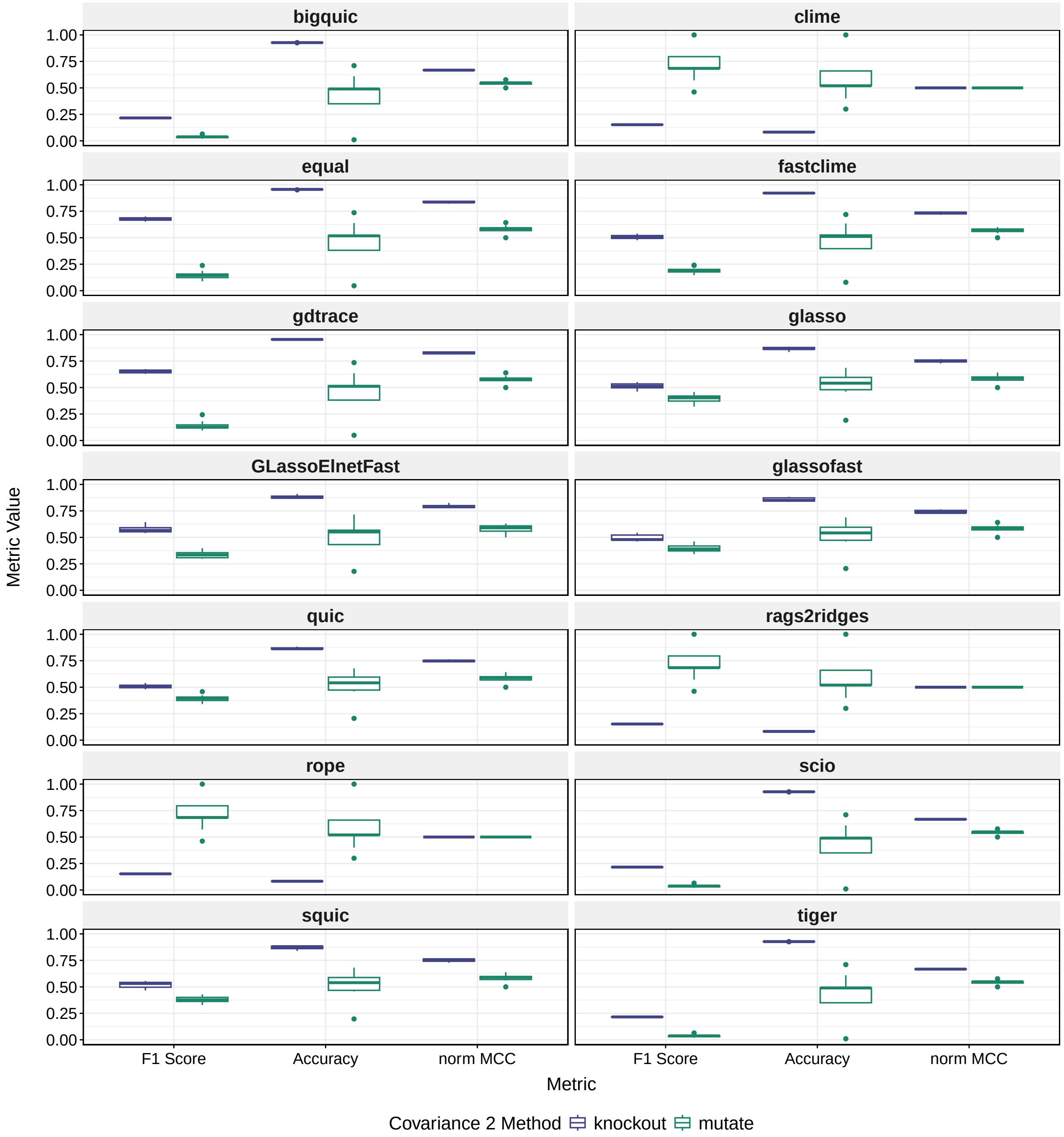

Supplementary Figure 9: Boxplot illustrating the influence of different covariance 2 methods on the performances of precision matrix estimation methods, assessed across the binary classification metrics F1 score, accuracy, and normalized MCC for $\widehat{\Theta}_2$. The performance of each method is represented by the distribution of the metric values over 10 replicates for each covariance 2 method. $\Sigma_1$ was generated using multiple block with $p = 100$. The sample data set was created using the mvrnorm sampling method with $n = 50$.

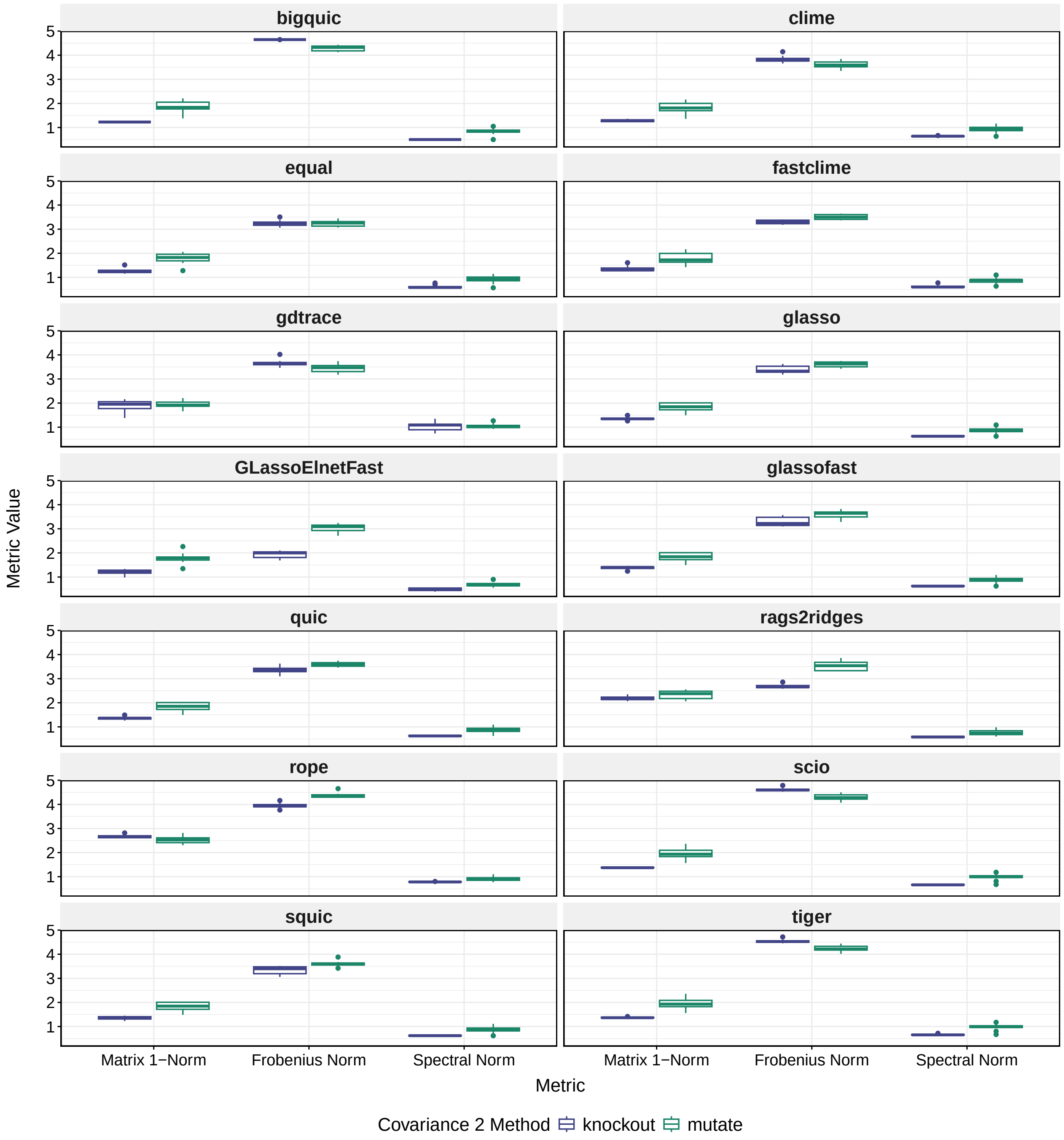

Supplementary Figure 10: Boxplot illustrating the influence of different covariance 2 methods on the performances of precision matrix estimation methods, assessed across the matrix norms matrix 1-norm, Frobenius norm, and spectral norm for $\widehat{\Theta}_2$. The performance of each method is represented by the distribution of the metric values over 10 replicates for each covariance 2 method. $\Sigma_1$ was generated using multiple block with $p = 100$. The sample data set was created using the mvrnorm sampling method with $n = 50$.

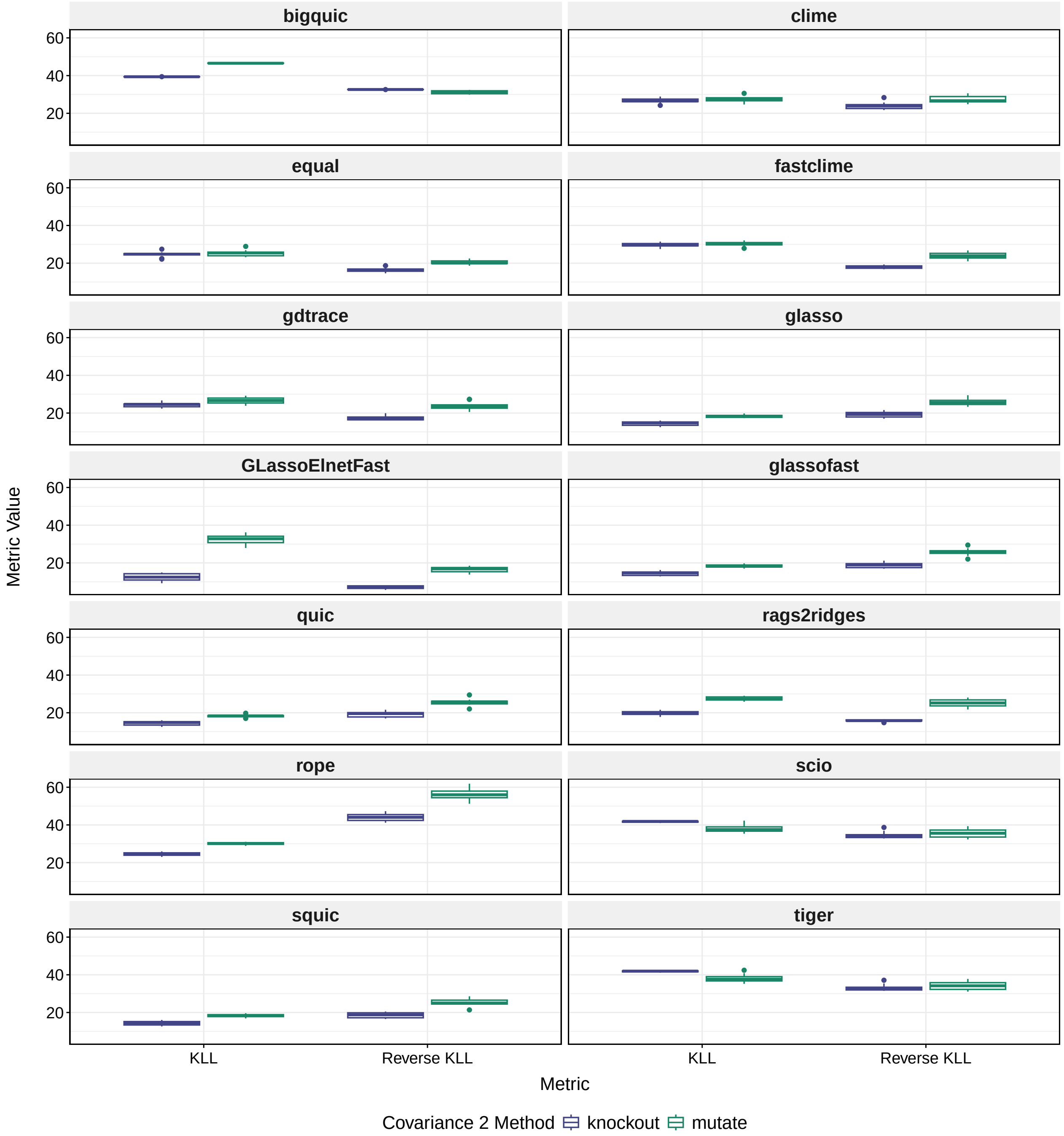

Supplementary Figure 11: Boxplot illustrating the influence of different covariance 2 methods on the performances of precision matrix estimation methods, assessed across the Kullback-Leibler Loss and the reverse Kullback-Leibler Loss for $\widehat{\Theta}_2$. The performance of each method is represented by the distribution of the metric values over 10 replicates for each covariance 2 method. $\Sigma_1$ was generated using multiple block with $p = 100$. The sample data set was created using the mvrnorm sampling method with $n = 50$.

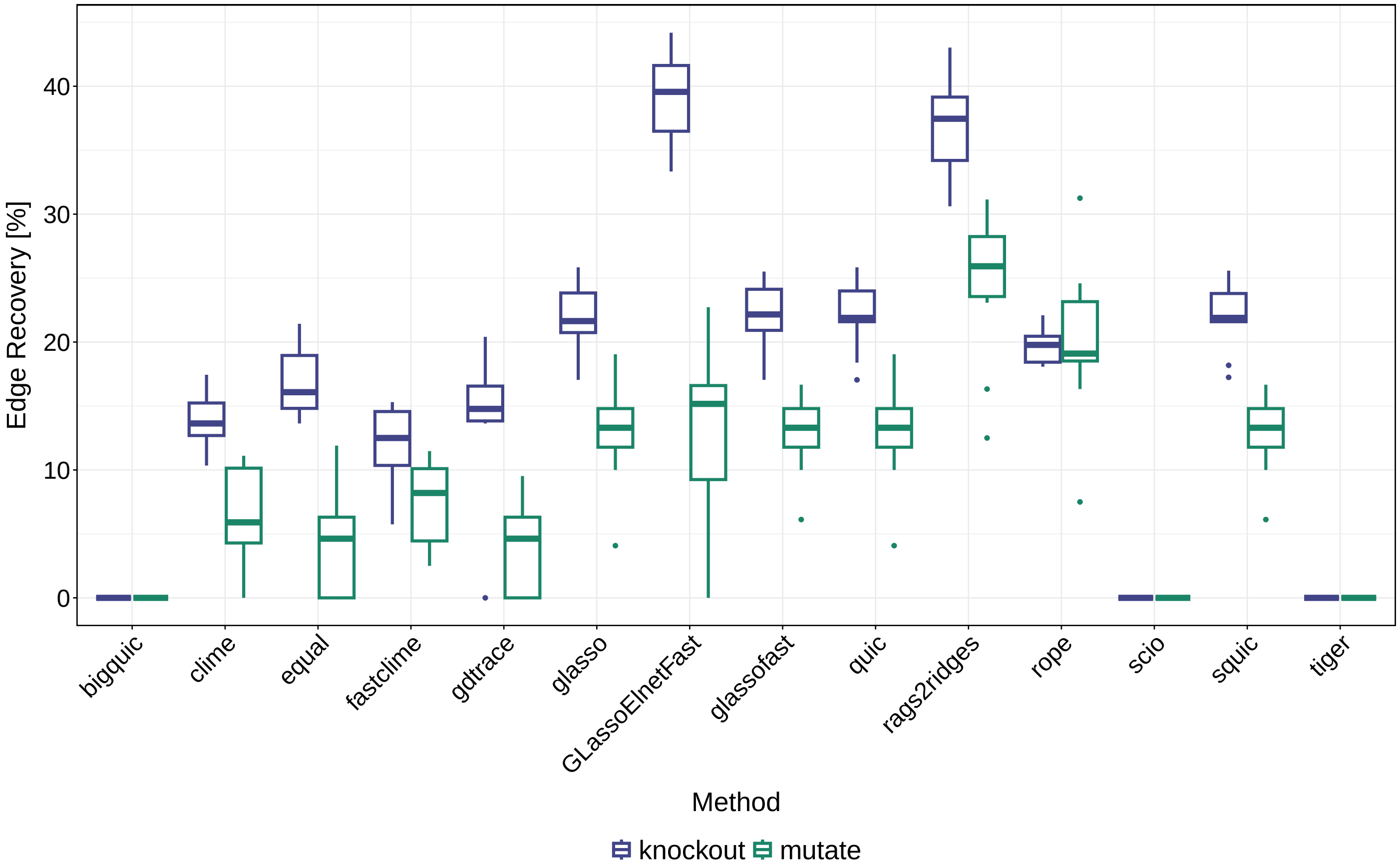

Supplementary Figure 12: Boxplot illustrating the influence of different covariance 2 methods on the performances of precision matrix estimation methods, assessed across by differential edge recovery rates. The performance of each method is represented by the distribution of the metric values over 10 replicates for each covariance 2 method. $\Sigma_1$ was generated using multiple block with $p = 100$. The sample data set was created using the mvrnorm sampling method with $n = 50$.

### 5 Supplementary Figures for Covariance Increase

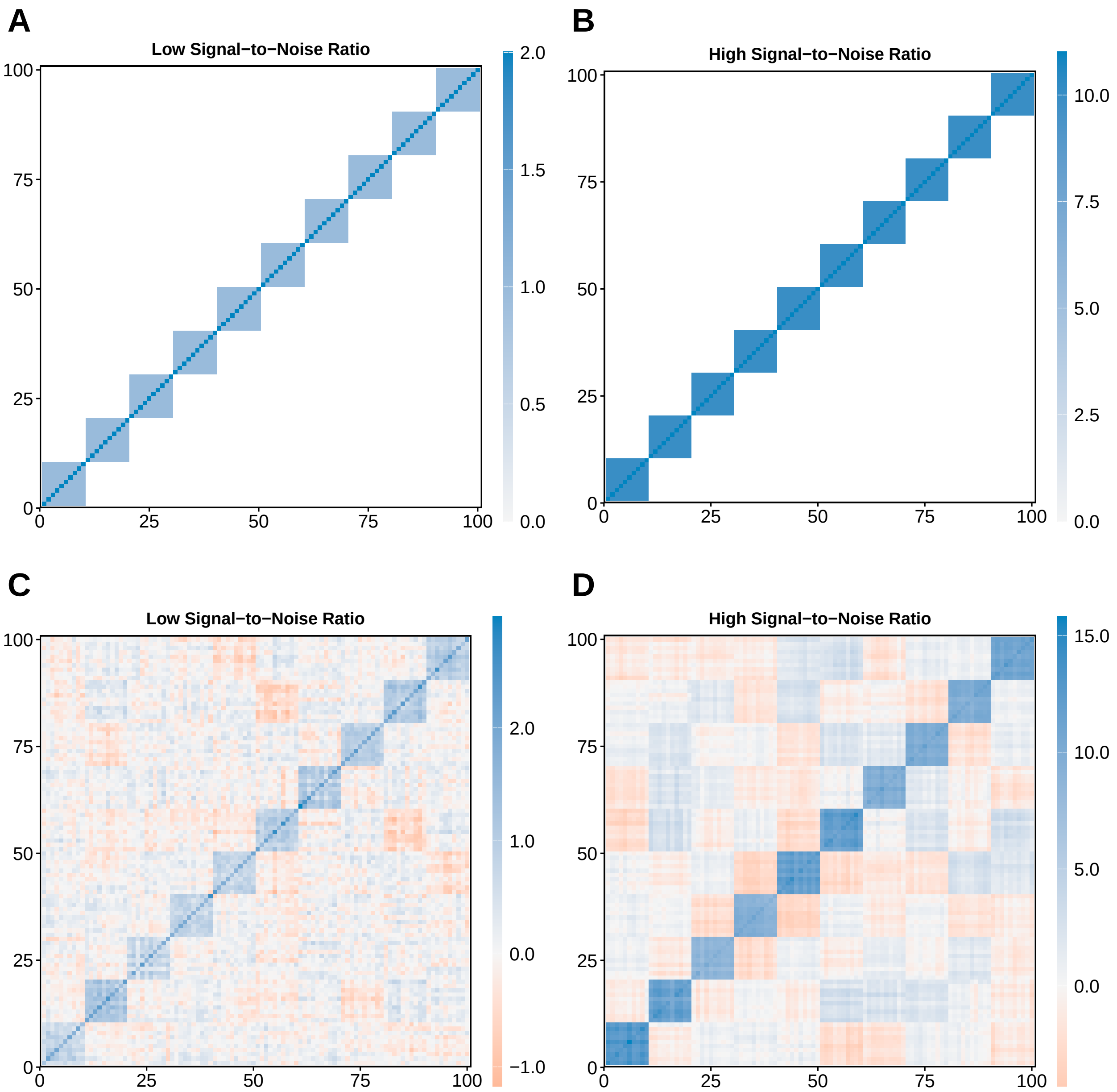

Supplementary Figure 13: A & B: $\Sigma_1$ generated with multiple block method using covariances of 1 or 10 and variances 2 or 11, respectively. C & D: $S_1$ generated with mvrnorm based on the $\Sigma_1$ in A & B.

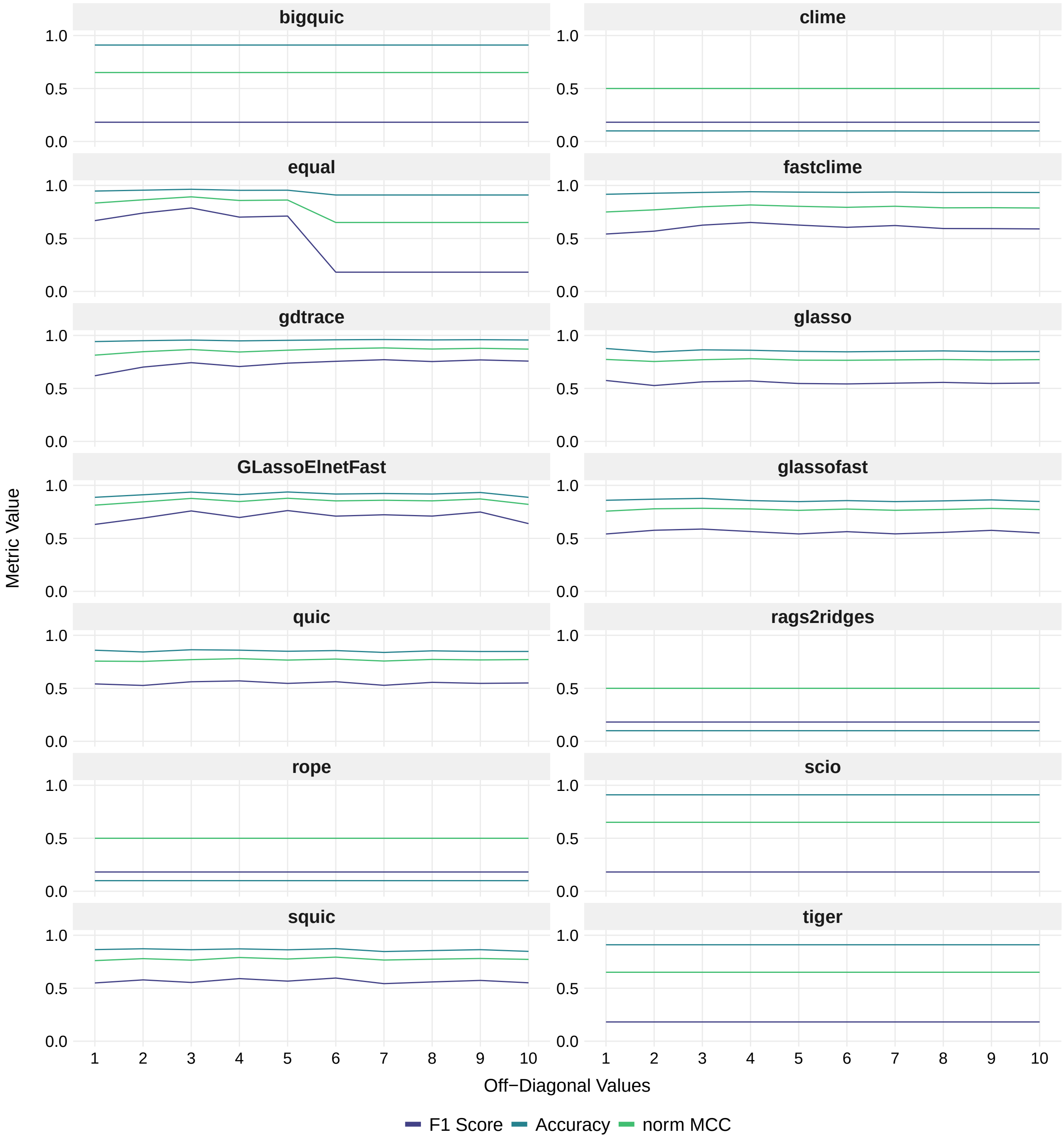

Supplementary Figure 14: Line plot illustrating the influence of different off-diagonal values (covariances) on the performances of precision matrix estimation methods, assessed across the binary classification metrics F1 score, accuracy, and normalized MCC. The covariances in $\Sigma_1$ and thus also $\Sigma_2$ were varied, ranging from 1 to 10. The diagonal values (variances) were set to 2 to 11, respectively. $\Sigma_1$ was generated using the covariance 1 method multiple block with $p = 100$. $\Sigma_2$ was generated based on $\Sigma_1$ using the covariance 2 method knockout. The simulated data sets were created with the sampling method mvrnorm with $n = 50$.

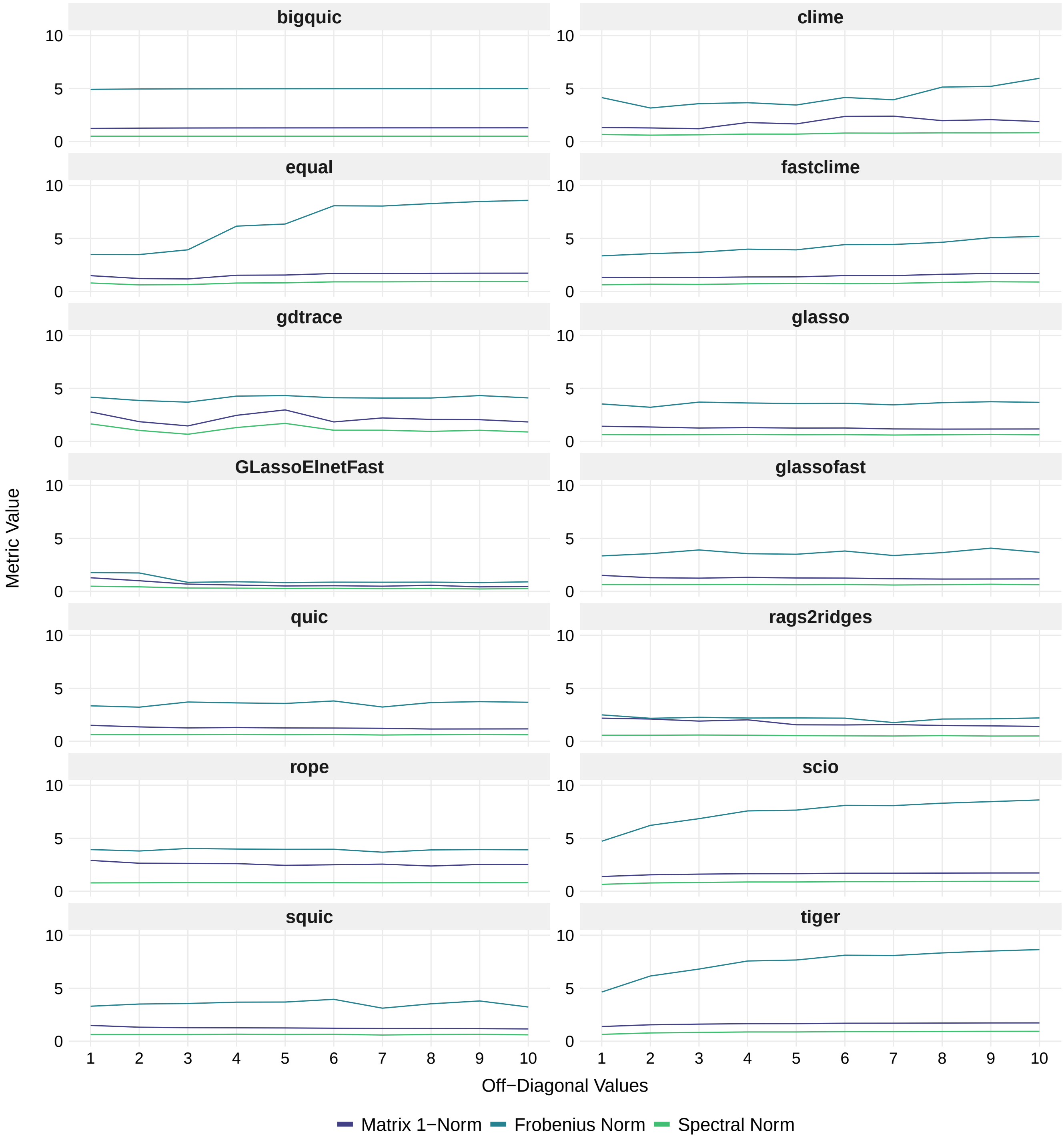

Supplementary Figure 15: Line plot illustrating the influence of different off-diagonal values (covariances) on the performances of precision matrix estimation methods, assessed across the matrix norms matrix 1-norm, Frobenius norm, and spectral norm. The covariances in $\Sigma_1$ and thus also $\Sigma_2$ were varied, ranging from 1 to 10. The diagonal values (variances) were set to 2 to 11, respectively. $\Sigma_1$ was generated using the covariance 1 method multiple block with $p = 100$. $\Sigma_2$ was generated based on $\Sigma_1$ using the covariance 2 method knockout. The simulated data sets were created with the sampling method mvrnorm with $n = 50$.

### 6 Supplementary Figures for Dimensions

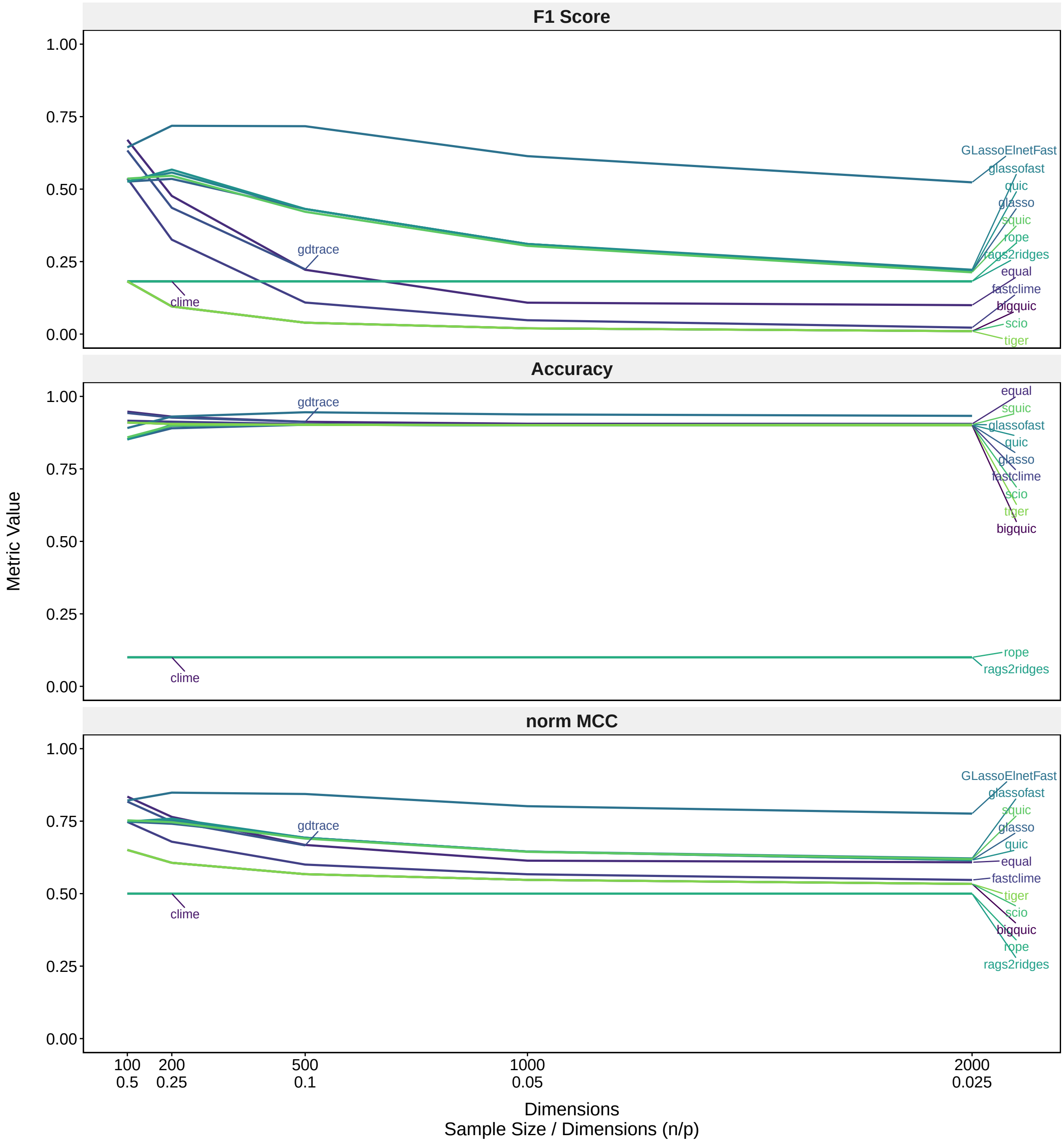

Supplementary Figure 16: Line plot illustrating the impact of increasing the dimensions on the performances of precision matrix estimation methods, assessed across the binary classification metrics F1 score, accuracy, and normalized MCC for $\widehat{\Theta}_1$. Dimensions from 100 to 2000 were analysed. $\Sigma_1$ was generated using the covariance 1 method multiple block, using $p = 100, 200, 500, 1000, 2000$. The sample data sets were created using the sampling method mvrnorm with $n = 50$. *Clime* and *gdtrace* were only analysed in the approaches with $p = 100$ and $p = 100, 200$, respectively.

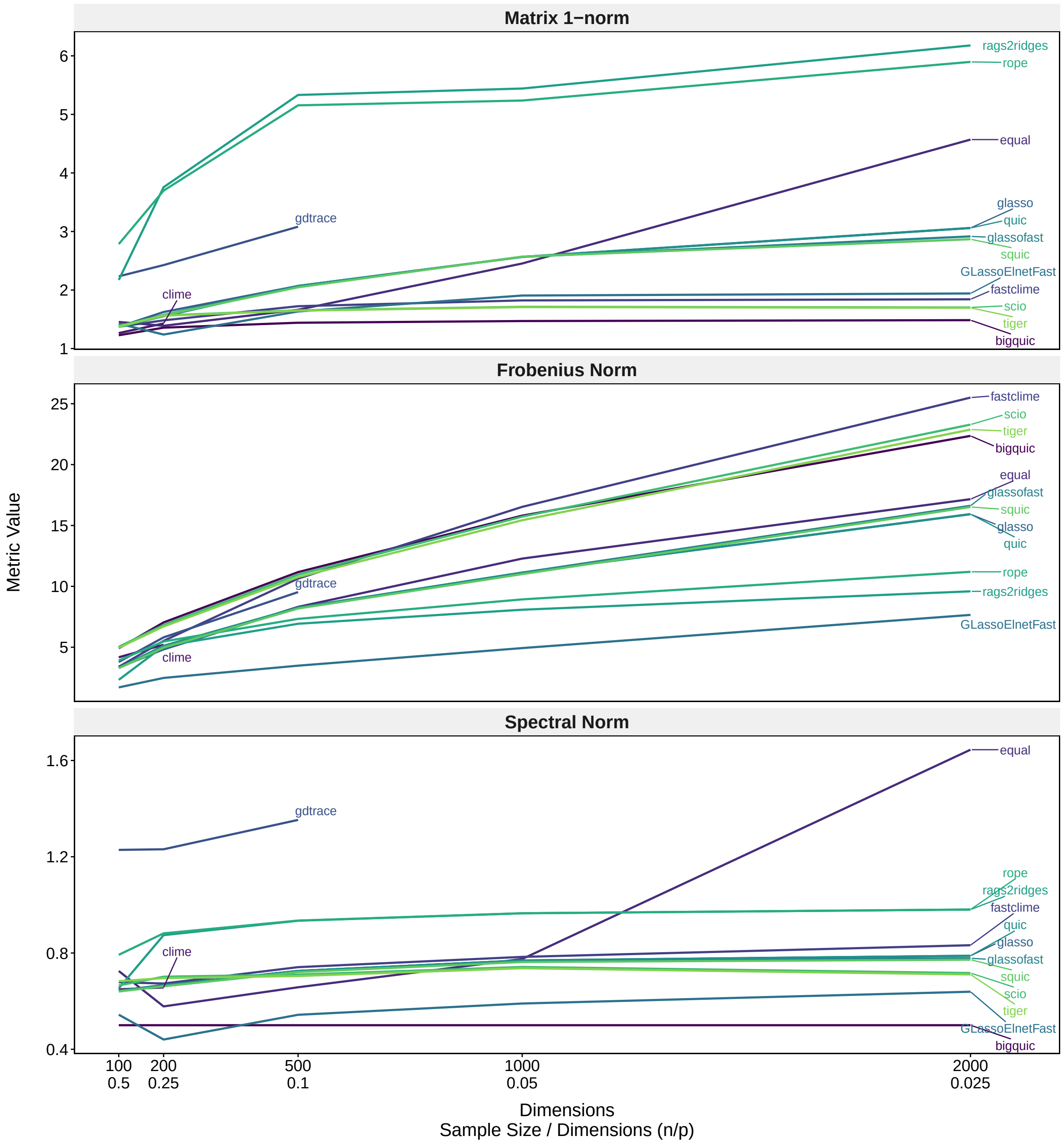

Supplementary Figure 17: Line plot illustrating the impact of increasing the dimensions on the performances of precision matrix estimation methods, assessed across the matrix norms matrix 1-norm, Frobenius norm, and spectral norm for $\widehat{\Theta}_1$. Dimensions from 100 to 2000 were analysed. $\Sigma_1$ was generated using the covariance 1 method multiple block, using $p = 100, 200, 500, 1000, 2000$. The sample data sets were created using the sampling method mvrnorm with $n = 50$. *Clime* and *gdtrace* were only analysed in the approaches with $p = 100$ and $p = 100, 200$, respectively.

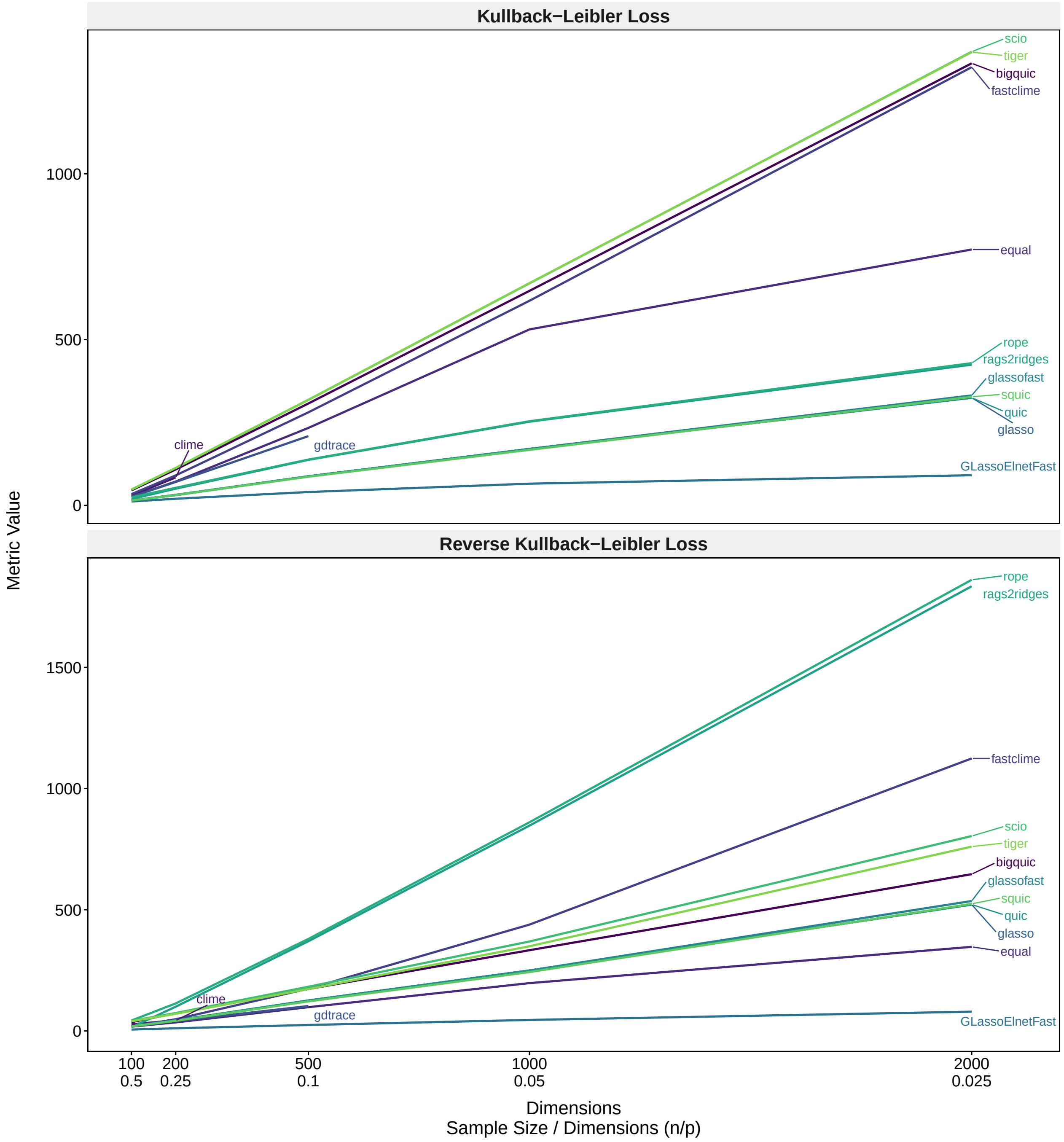

Supplementary Figure 18: Line plot illustrating the impact of increasing the dimensions on the performances of precision matrix estimation methods, assessed across the Kullback-Leibler Loss and the reverse Kullback-Leibler Loss for $\widehat{\Theta}_1$. Dimensions from 100 to 2000 were analysed. $\Sigma_1$ was generated using the covariance 1 method multiple block, using $p = 100, 200, 500, 1000, 2000$. The sample data sets were created using the sampling method mvrnorm with $n = 50$. *Clime* and *gdtrace* were only analysed in the approaches with $p = 100$ and $p = 100, 200$, respectively.

### 7 Supplementary Figures for Means

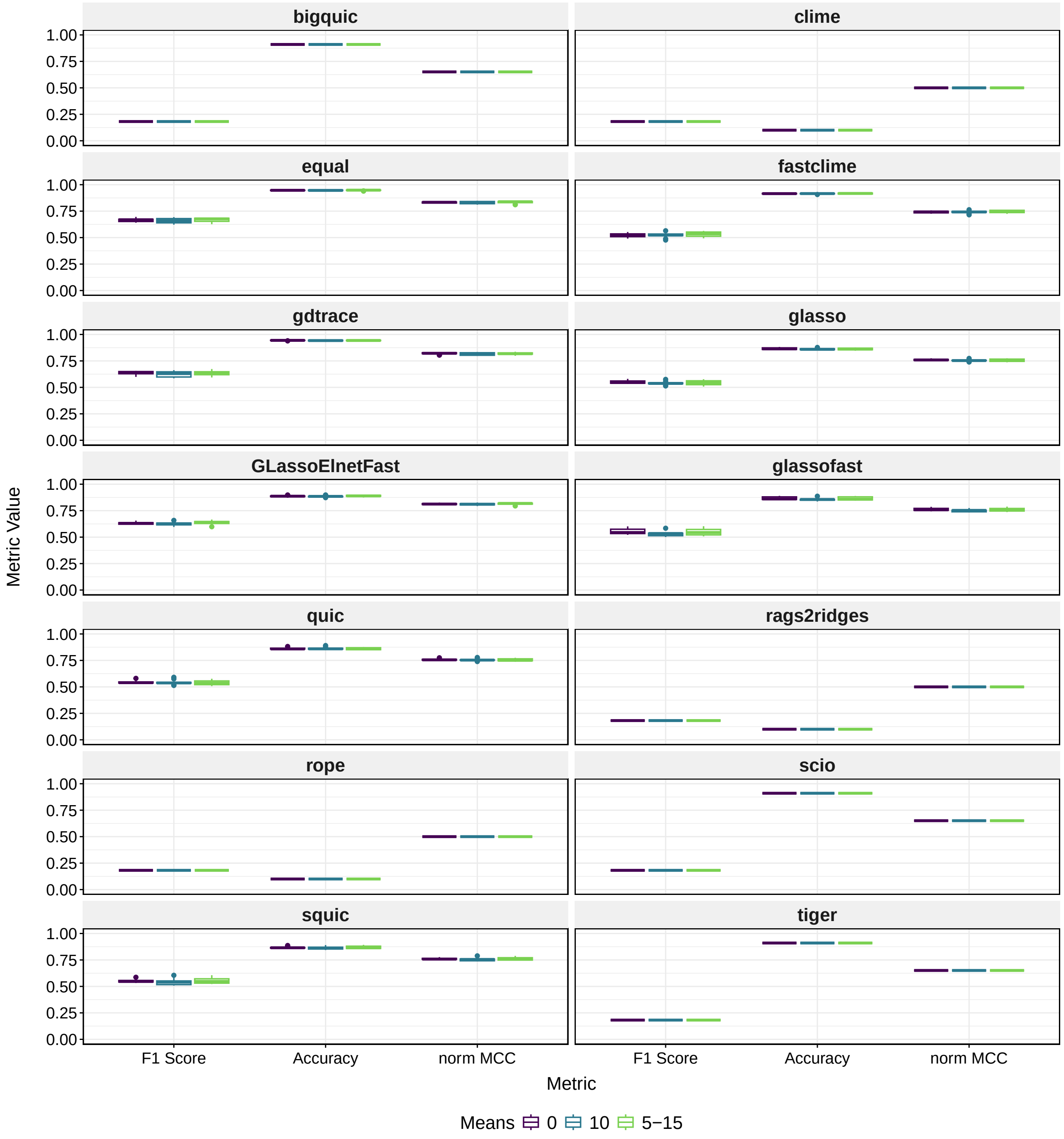

Supplementary Figure 19: Boxplot illustrating the effect of shifting means on the performance of different PMEMs in repeated simulations, assessed across the binary classification metrics F1 score, accuracy, and normalized MCC. The performance of each method is represented by the distribution of the metric values over 10 replicates for each mean. $\Sigma_1$ was generated using the covariance 1 method multiple block, using $p = 100$. The sample datasets were generated using the mvrnorm sampling method with $n = 50$, and the variable means were set to 0, to 10, or randomly drawn from the range 5 to 15.

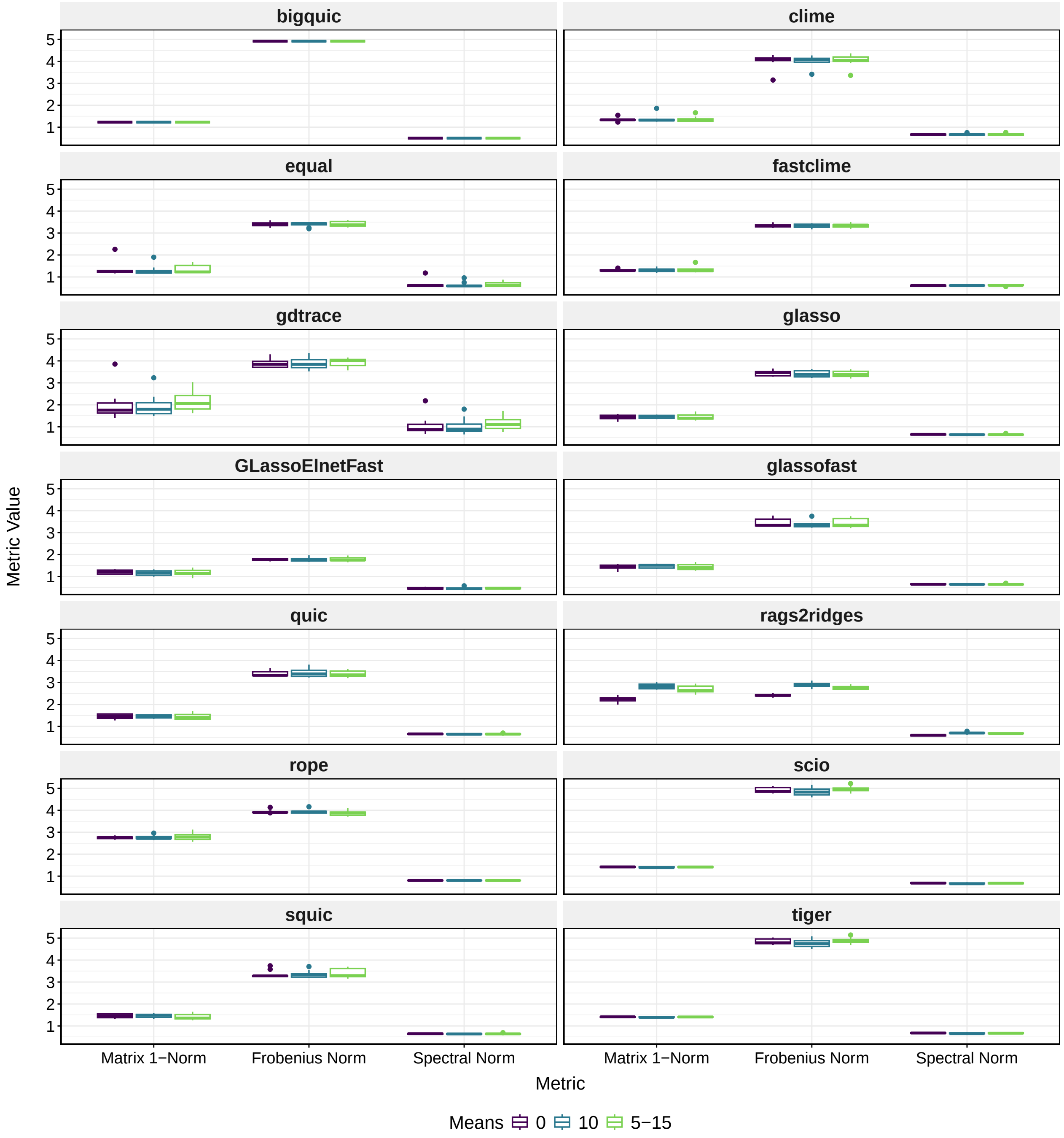

Supplementary Figure 20: Boxplot illustrating the effect of shifting means on the performance of different PMEMs in repeated simulations, assessed across the matrix norms matrix 1-norm, Frobenius norm, and spectral norm. The performance of each method is represented by the distribution of the metric values over 10 replicates for each mean. $\Sigma_1$ was generated using the covariance 1 method multiple block, using $p = 100$. The sample datasets were generated using the mvrnorm sampling method with $n = 50$, and the variable means were set to 0, to 10, or randomly drawn from the range 5 to 15.

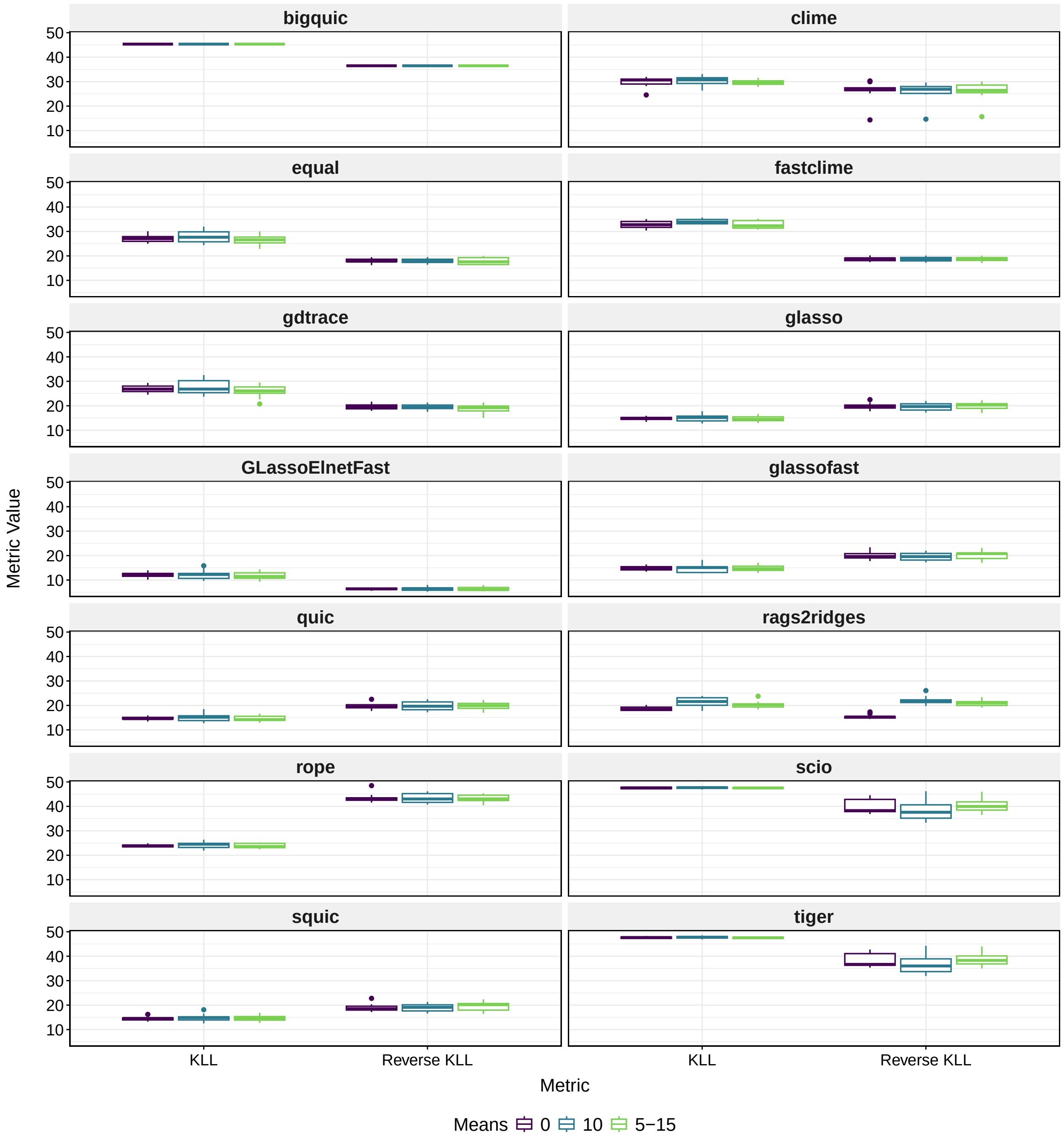

Supplementary Figure 21: Boxplot illustrating the effect of shifting means on the performance of different PMEMs in repeated simulations, assessed across the Kullback-Leibler Loss and the reverse Kullback-Leibler Loss. The performance of each method is represented by the distribution of the metric values over 10 replicates for each mean. $\Sigma_1$ was generated using the covariance 1 method multiple block, using $p = 100$. The sample datasets were generated using the mvrnorm sampling method with $n = 50$, and the variable means were set to 0, to 10, or randomly drawn from the range 5 to 15.

### 8 Supplementary Figures for Normalization

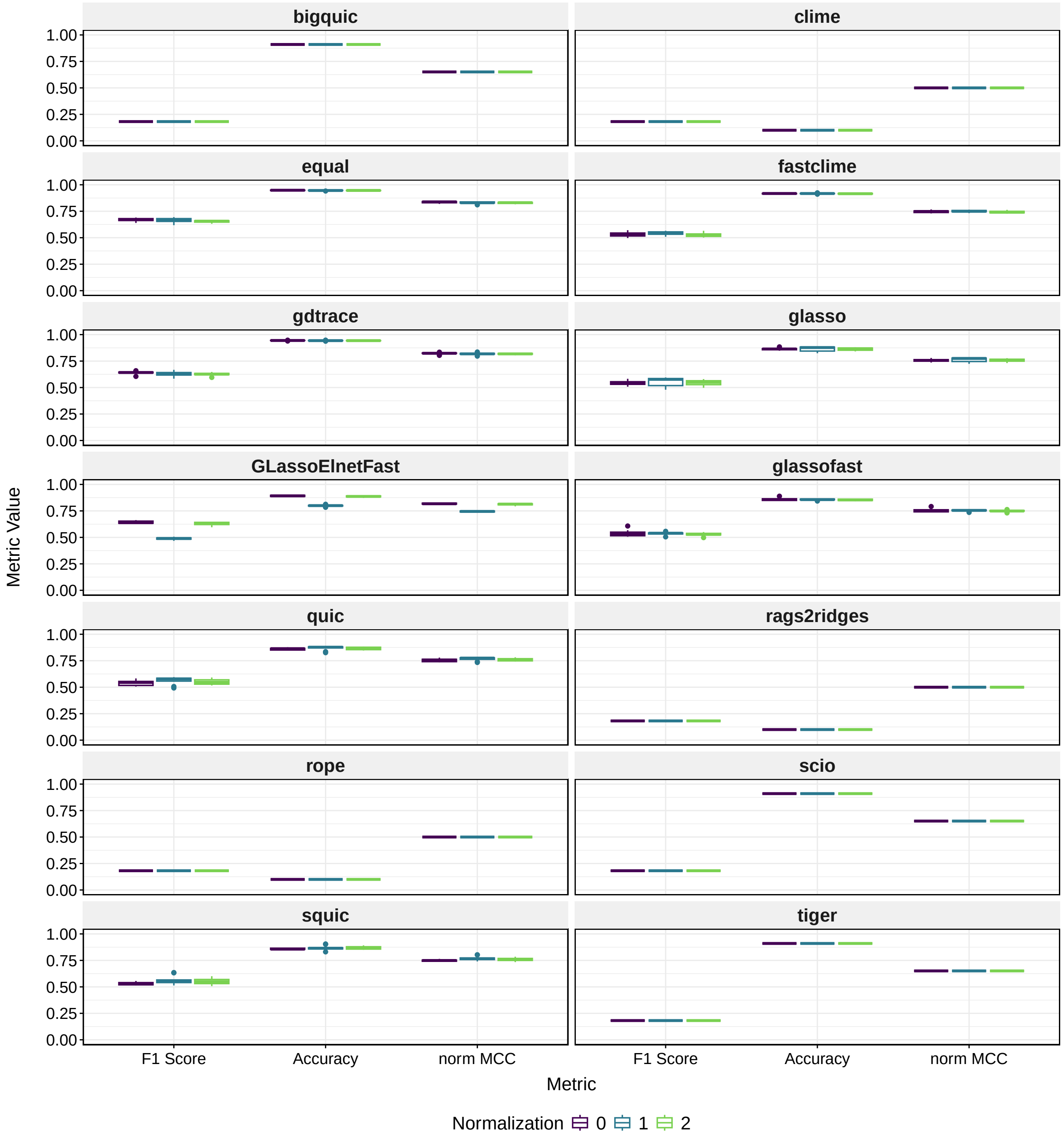

Supplementary Figure 22: Boxplot illustrating the effect of different normalization methods on the performance of different PMEMs in repeated simulations, assessed across the binary classification metrics F1 score, accuracy, and normalized MCC. The performance of each method is represented by the distribution of the metric values over 10 replicates for each mean. $\Sigma_1$ was generated using the covariance 1 method multiple block, using $p = 100$. The sample datasets were generated using the mvrnorm sampling method with $n = 50$, and the variable means were randomly drawn from the range 5 to 15. After sampling the following normalization strategies are applied, 0: no normalization, 1: normalizing means to 0 and variances to 1, 2: normalizing means to 0.

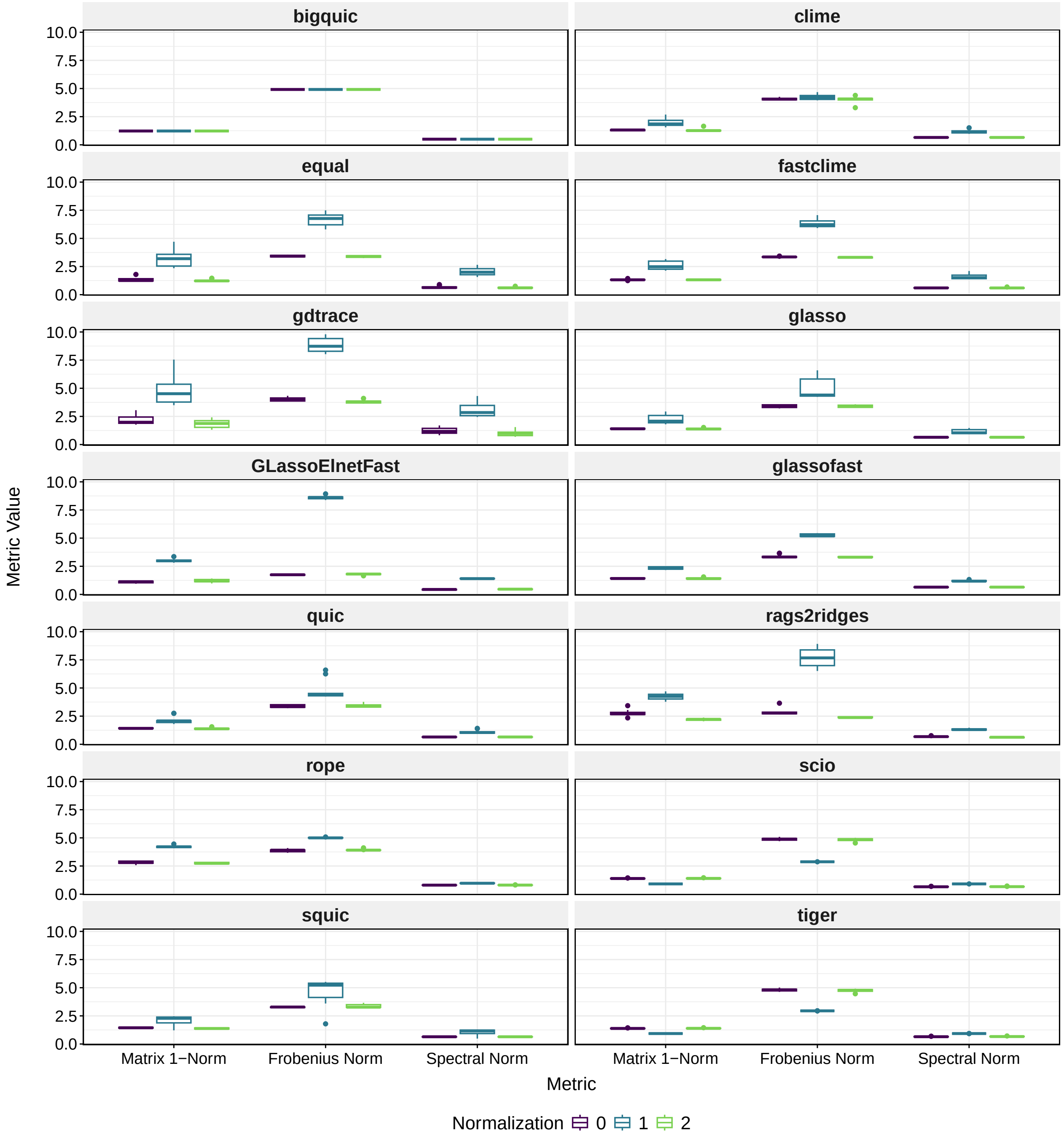

Supplementary Figure 23: Boxplot illustrating the effect of different normalization methods on the performance of different PMEMs in repeated simulations, assessed across the matrix norms matrix 1-norm, Frobenius norm, and spectral norm. The performance of each method is represented by the distribution of the metric values over 10 replicates for each mean. $\Sigma_1$ was generated using the covariance 1 method multiple block, using $p = 100$. The sample datasets were generated using the mvrnorm sampling method with $n = 50$, and the variable means were randomly drawn from the range 5 to 15. After sampling the following normalization strategies are applied, 0: no normalization, 1: normalizing means to 0 and variances to 1, 2: normalizing means to 0.

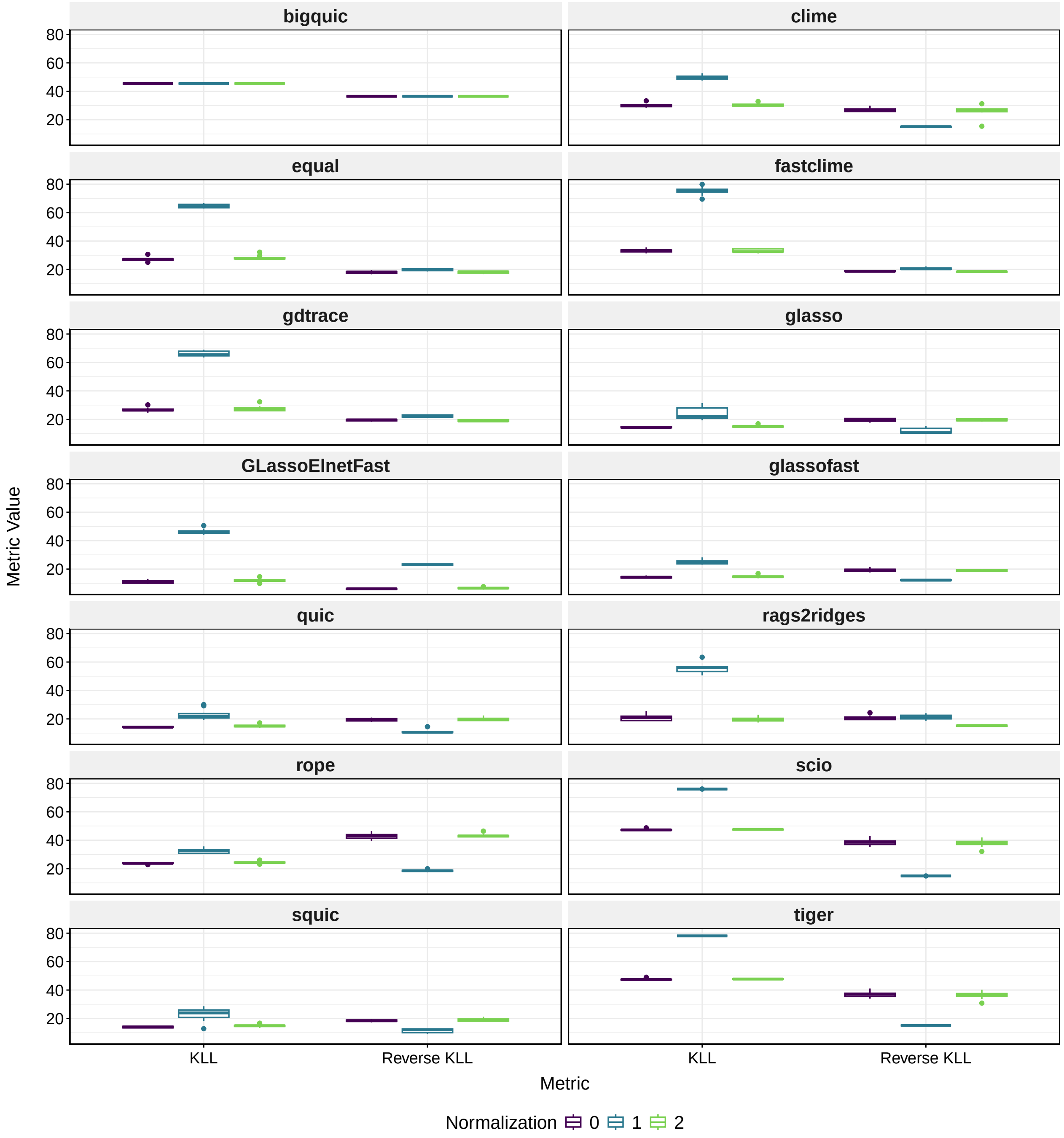

Supplementary Figure 24: Boxplot illustrating the effect of different normalization methods on the performance of different PMEMs in repeated simulations, assessed across the Kullback-Leibler Loss and the reverse Kullback-Leibler Loss. The performance of each method is represented by the distribution of the metric values over 10 replicates for each mean. $\Sigma_1$ was generated using the covariance 1 method multiple block, using $p = 100$. The sample datasets were generated using the mvrnorm sampling method with $n = 50$, and the variable means were randomly drawn from the range 5 to 15. After sampling the following normalization strategies are applied, 0: no normalization, 1: normalizing means to 0 and variances to 1, 2: normalizing means to 0.

### 9 Supplementary Figures for Sample Size

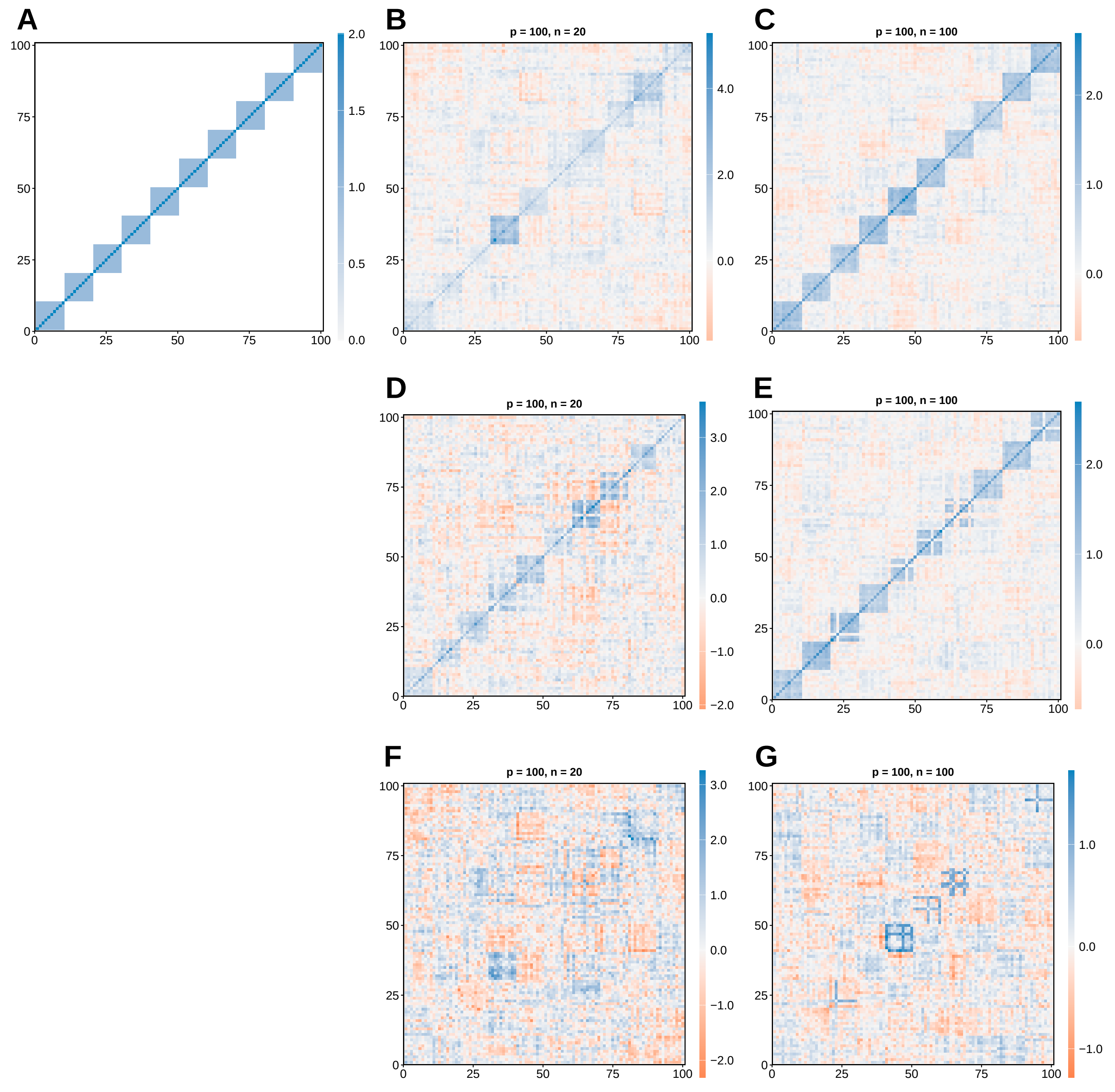

Supplementary Figure 25: A: $\Sigma_1$ generated with the multiple block method, using $p = 100$. B & C: $S_1$ generated with the mvrnorm method based on $\Sigma_1$, using $n = 20, 100$, respectively. D & E: $S_2$ generated with the mvrnorm method, using $n = 20, 100$, respectively. F & G: The respective differential sample covariance matrices $S_\Delta$.

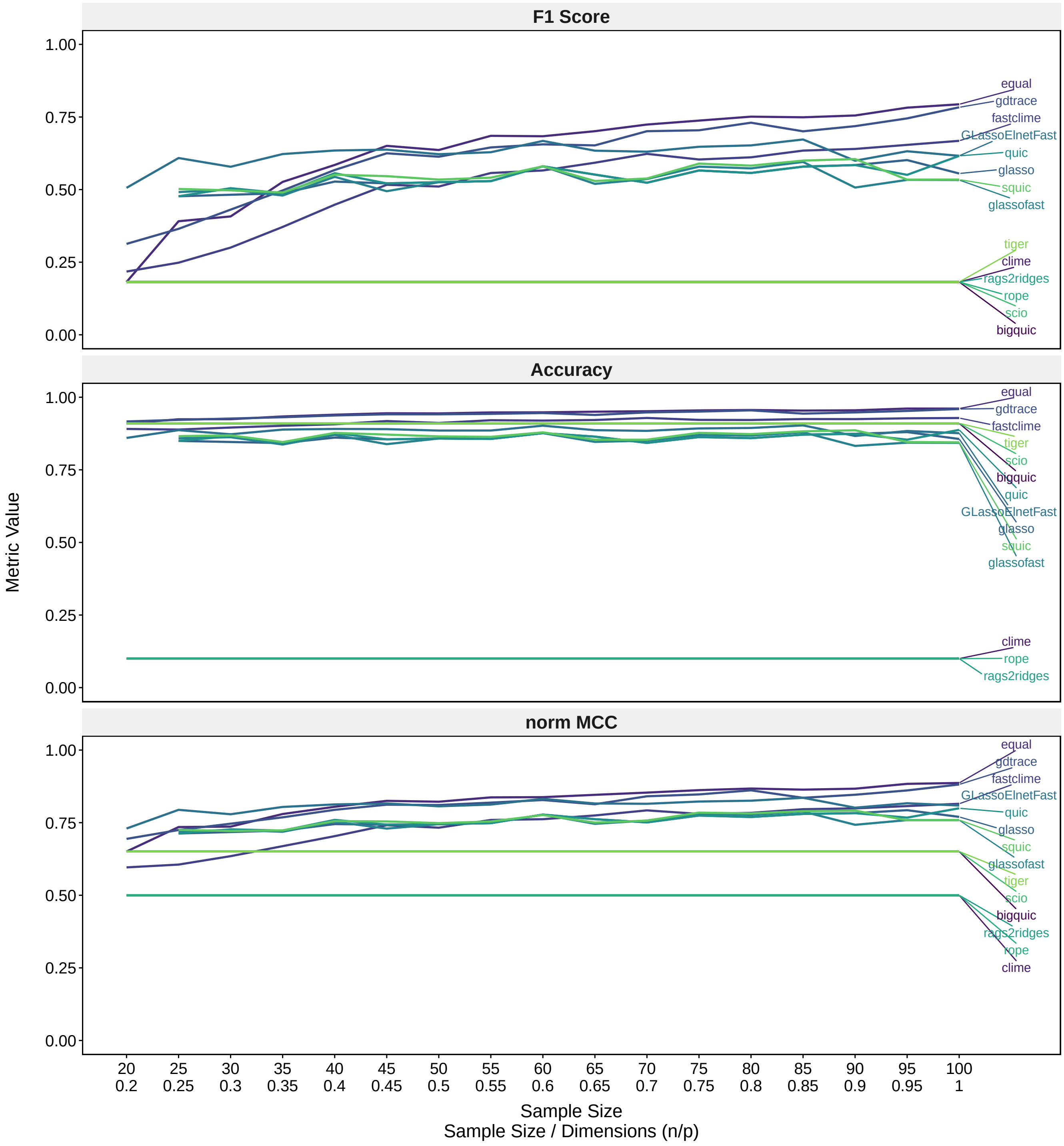

Supplementary Figure 26: Line plot illustrating the impact of increasing sample sizes on the performances of precision matrix estimation methods, assessed across the binary classification metrics F1 score, accuracy, and normalized MCC. Sample sizes from 20 to 100 were analysed. $\Sigma_1$ was generated using the covariance 1 method multiple block, with $p = 100$. The simulated data sets were created using the sampling method mvrnorm with $n = 20, 25, \ldots, 95, 100$.

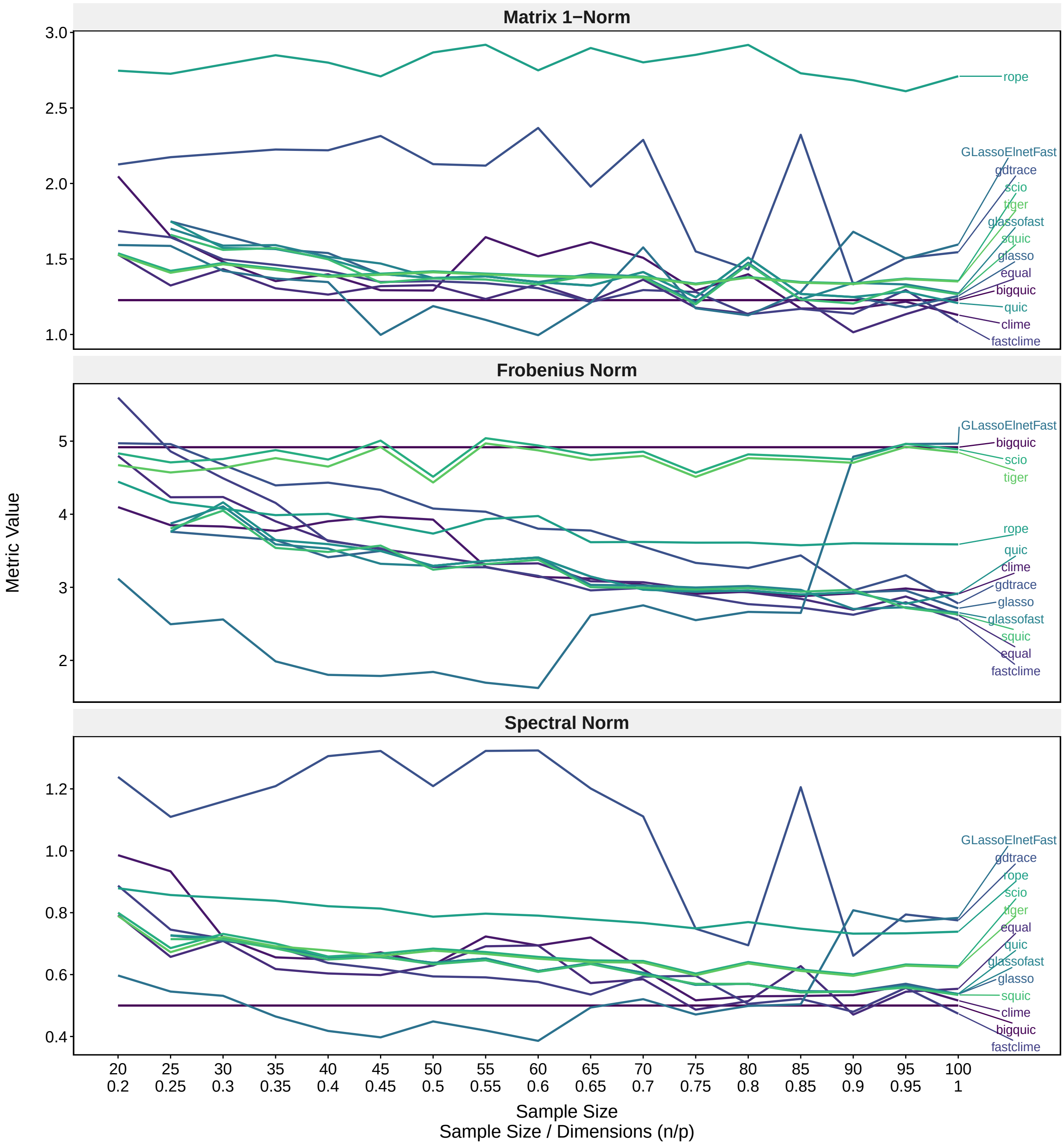

Supplementary Figure 27: Line plot illustrating the impact of increasing sample sizes on the performances of precision matrix estimation methods, assessed across the matrix norms matrix 1-norm, Frobenius norm, and spectral norm. Sample sizes from 20 to 100 were analysed. $\Sigma_1$ was generated using the covariance 1 method multiple block, with $p = 100$. The simulated data sets were created using the sampling method mvrnorm with $n = 20, 25, \ldots, 95, 100$. *Rags2ridges* is excluded from this graph as its exceptionally high norm values obscured the differences among the remaining methods. Figure S28 includes *rags2ridges*.

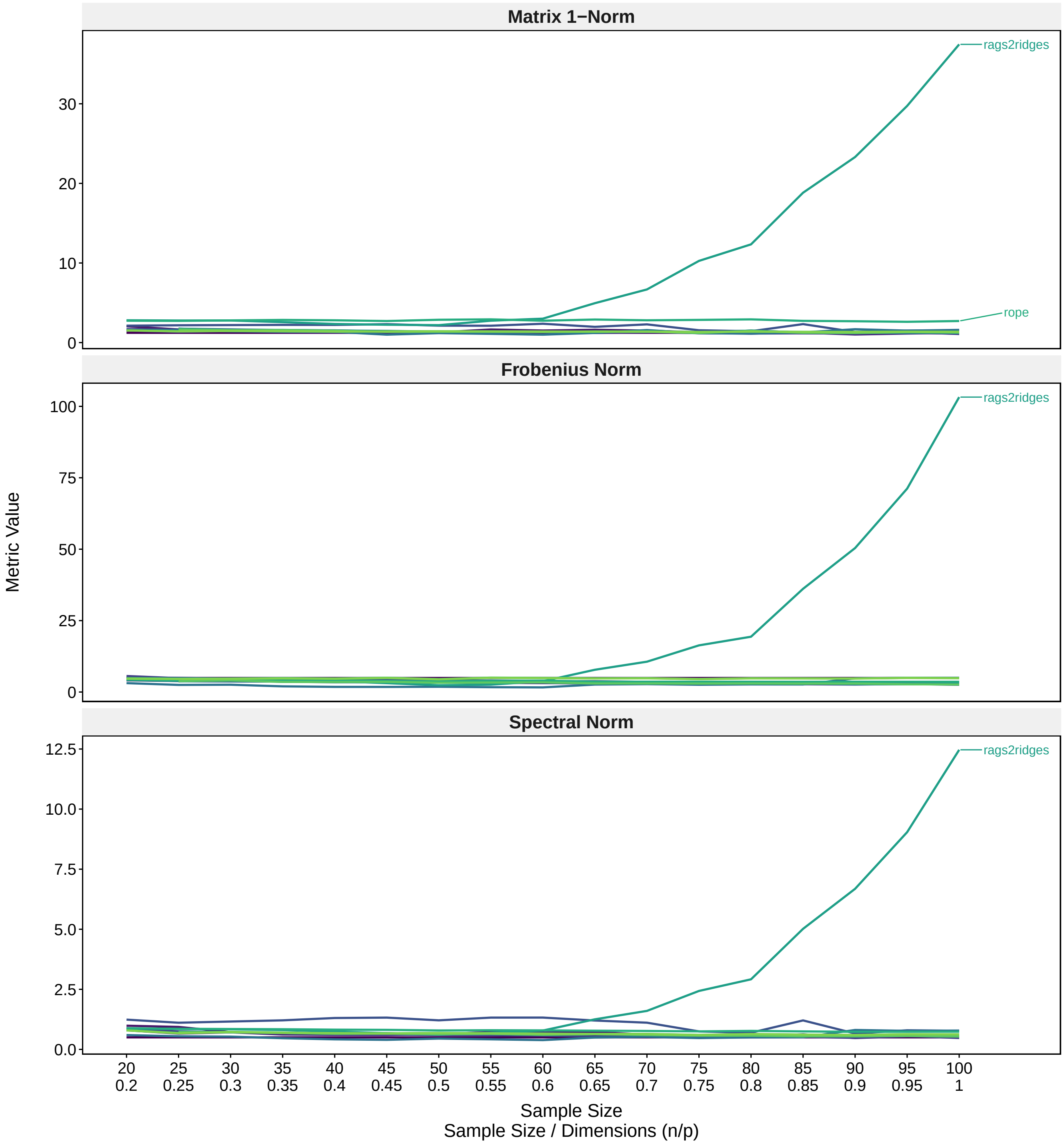

Supplementary Figure 28: Line plot illustrating the impact of increasing sample sizes on the performances of precision matrix estimation methods, assessed across the matrix norms matrix 1-norm, Frobenius norm, and spectral norm. Sample sizes from 20 to 100 were analysed. $\Sigma_1$ was generated using the covariance 1 method multiple block, with $p = 100$. The simulated data sets were created using the sampling method mvrnorm with $n = 20, 25, \ldots, 95, 100$.

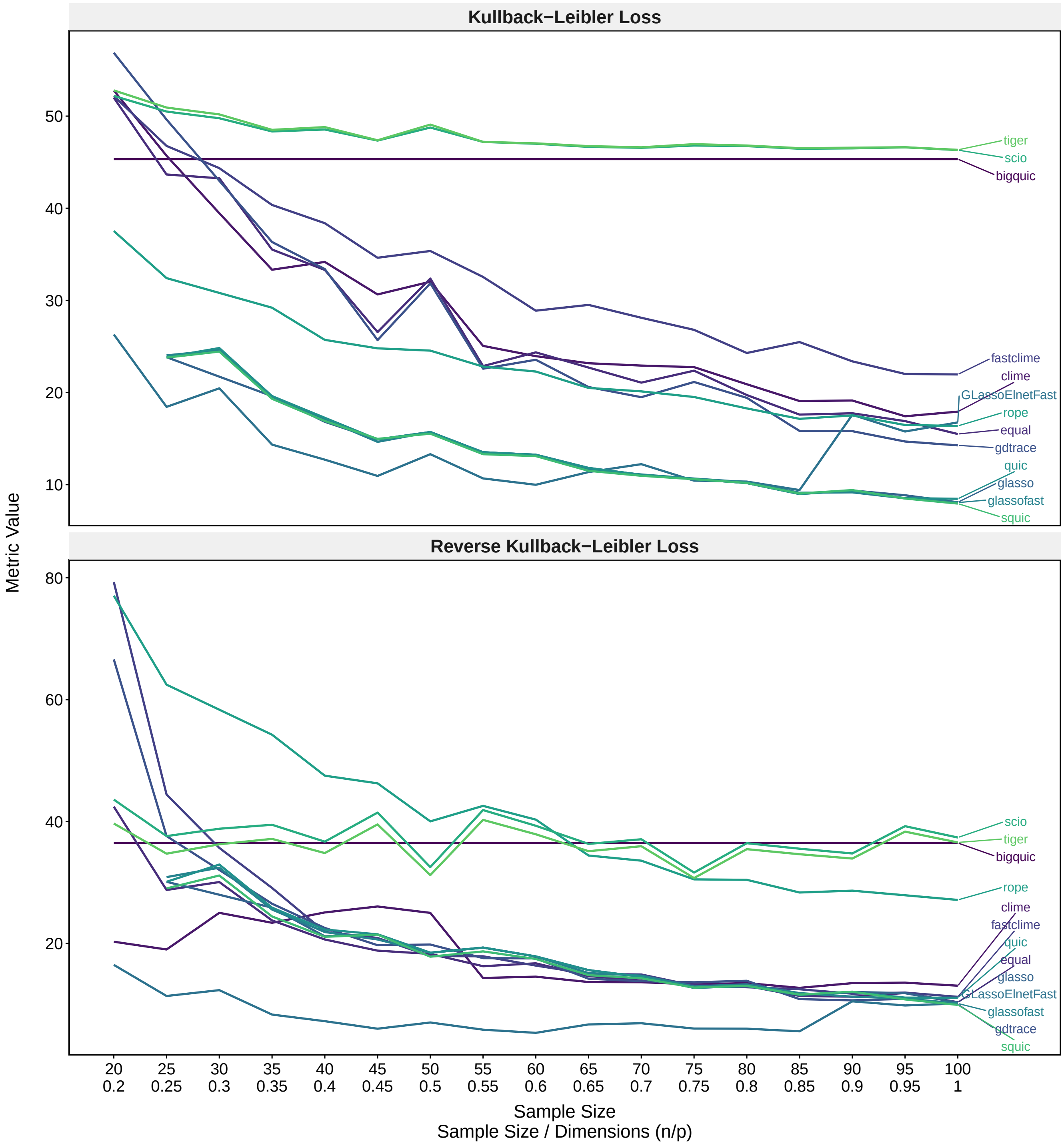

Supplementary Figure 29: Line plot illustrating the impact of increasing sample sizes on the performances of precision matrix estimation methods, assessed across across the Kullback-Leibler Loss and the reverse Kullback-Leibler Loss. Sample sizes from 20 to 100 were analysed. $\Sigma_1$ was generated using the covariance 1 method multiple block, with $p = 100$. The simulated data sets were created using the sampling method mvrnorm with $n = 20, 25, \ldots, 95, 100$. *Rags2ridges* is excluded from this graph as its exceptionally high loss values obscured the differences among the remaining methods. Figure S30 includes *rags2ridges*.

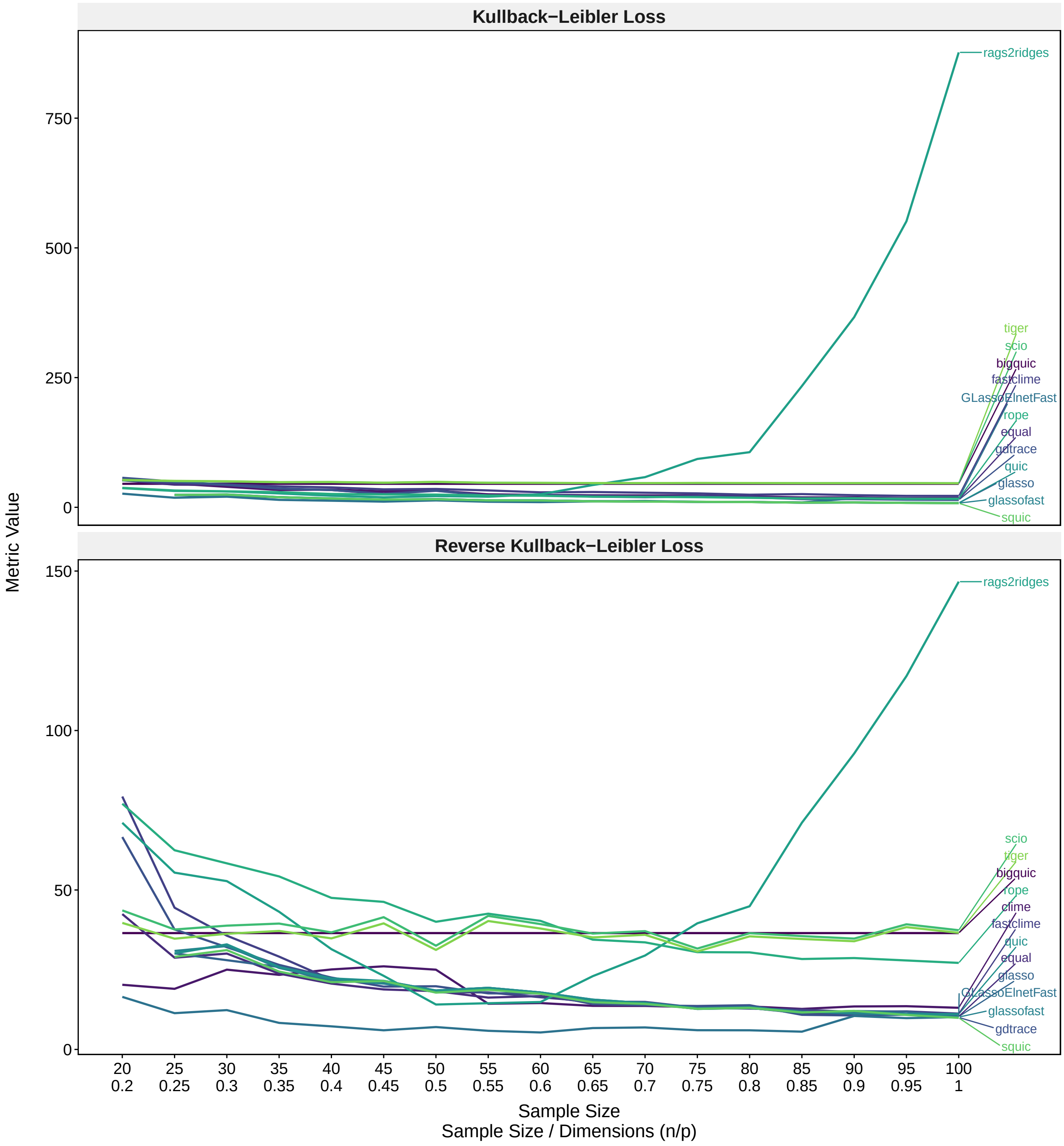

Supplementary Figure 30: Line plot illustrating the impact of increasing sample sizes on the performances of precision matrix estimation methods, assessed across across the Kullback-Leibler Loss and the reverse Kullback-Leibler Loss. Sample sizes from 20 to 100 were analysed. $\Sigma_1$ was generated using the covariance 1 method multiple block, with $p = 100$. The simulated data sets were created using the sampling method mvrnorm with $n = 20, 25, \ldots, 95, 100$.

### 10 Supplementary Figures for Runtimes

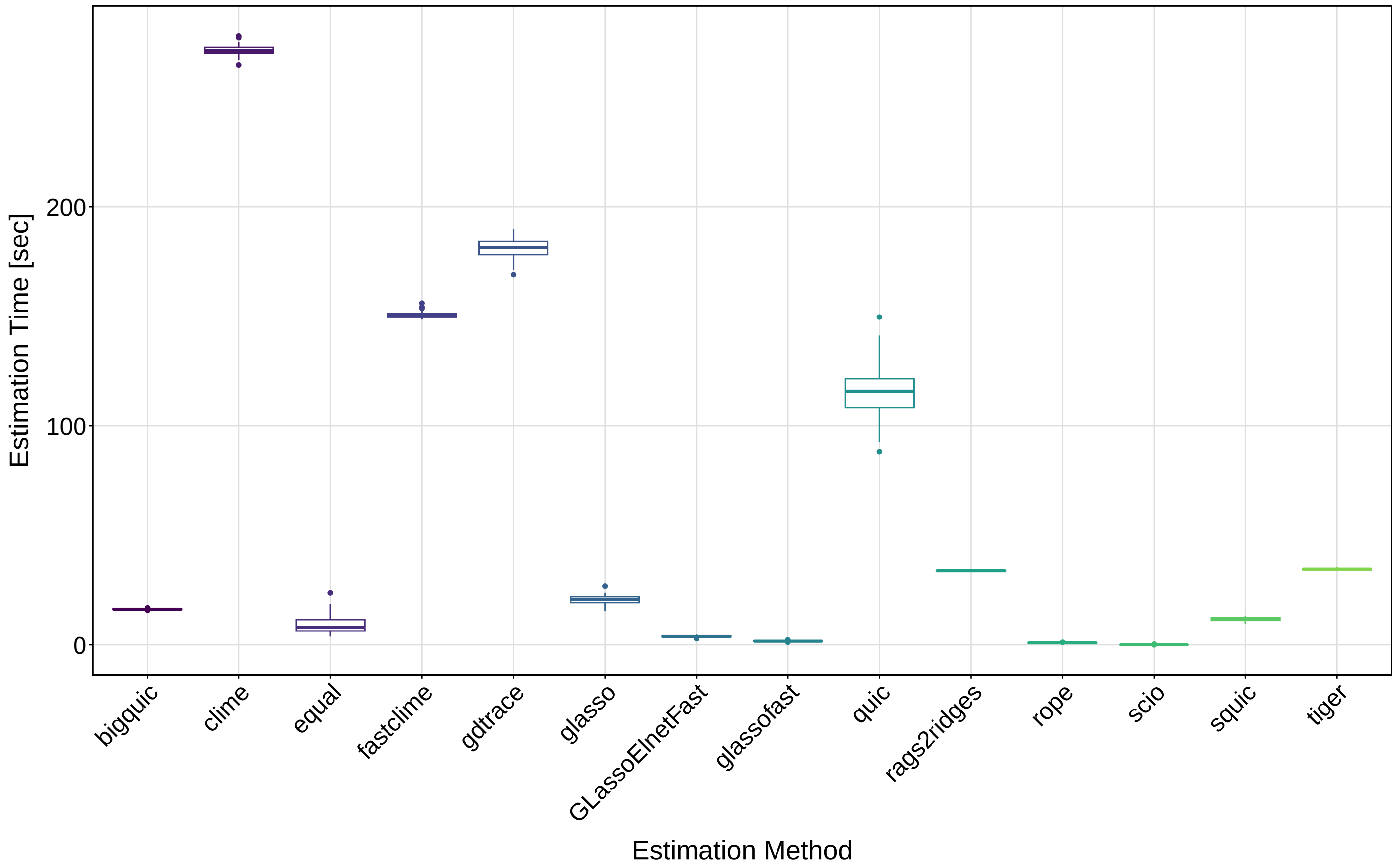

Supplementary Figure 31: Boxplot illustrating the variation in estimation times of different PMEMs over 50 replicates. $\Sigma_1$ was generated with the single block method, using $p = 100$. The sample data set was created using the mvrnorm sampling method with $n = 50$.

### 11 Supplementary Information for Sampling

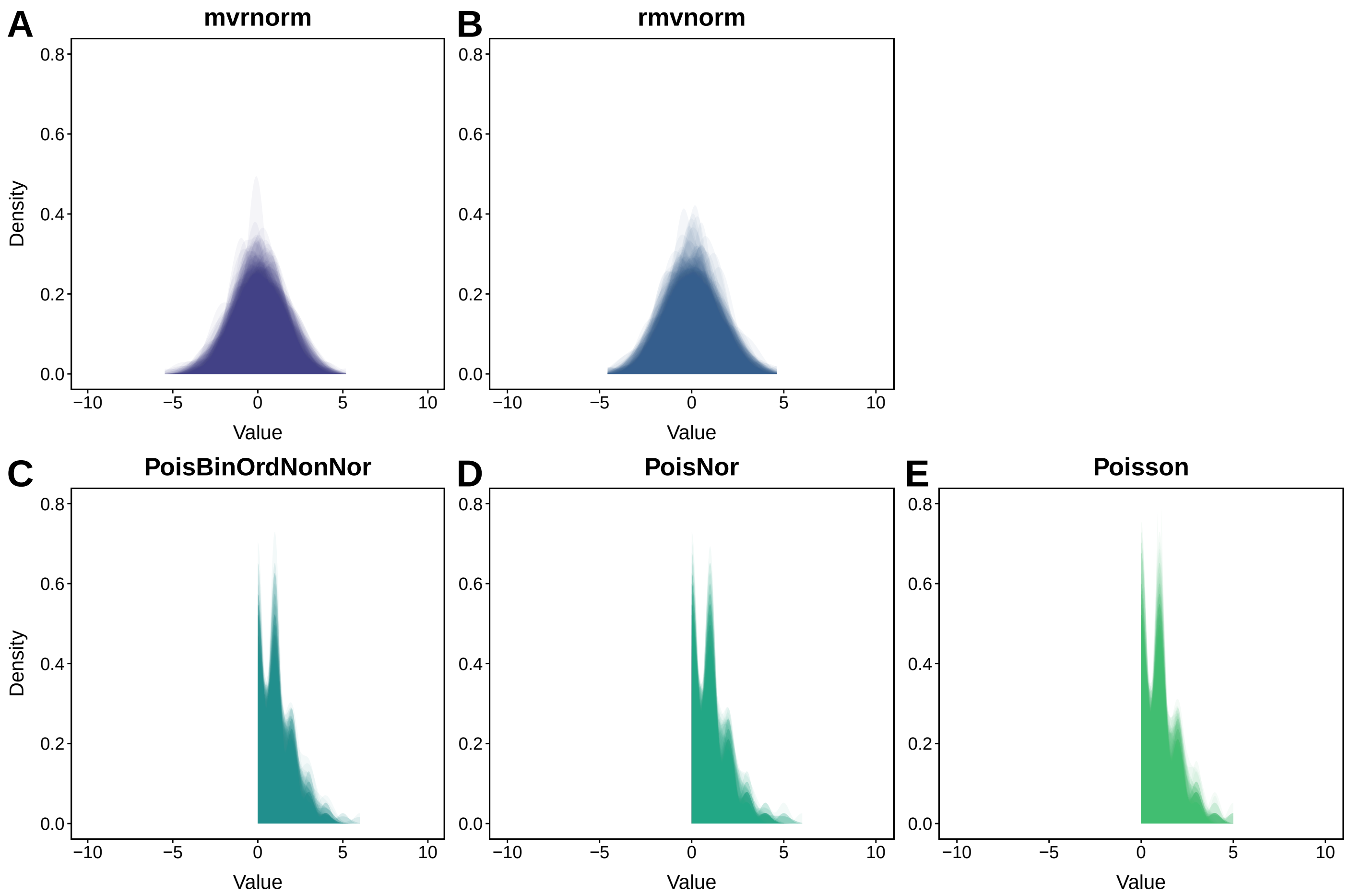

Supplementary Figure 32: Superimposed distributions of 100 variables across 50 samples from the simulated data sets for condition 1, generated by the sampling methods mvrnorm, rmvnorm, PoisNor, PoisBinOrdNonNor, and Poisson.

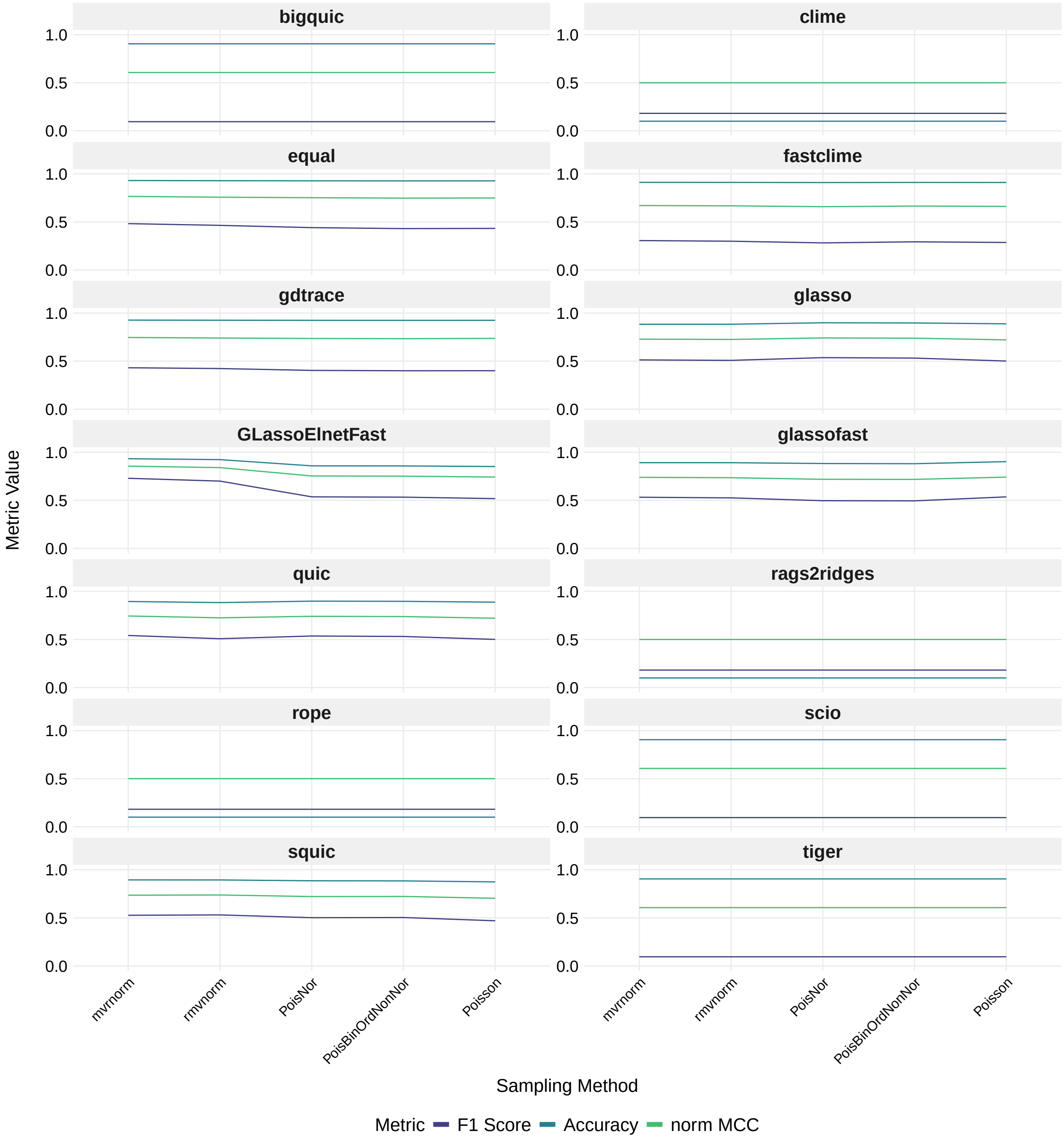

Supplementary Figure 33: Line plot illustrating the influence of different sampling methods on the performances of precision matrix estimation methods, assessed across the binary classification metrics F1 score, accuracy, and normalized MCC. $\Sigma_1$ was generated using the covariance 1 method multiple block with $p = 200$. The sample data set was created using the respective sampling method with $n = 50$.

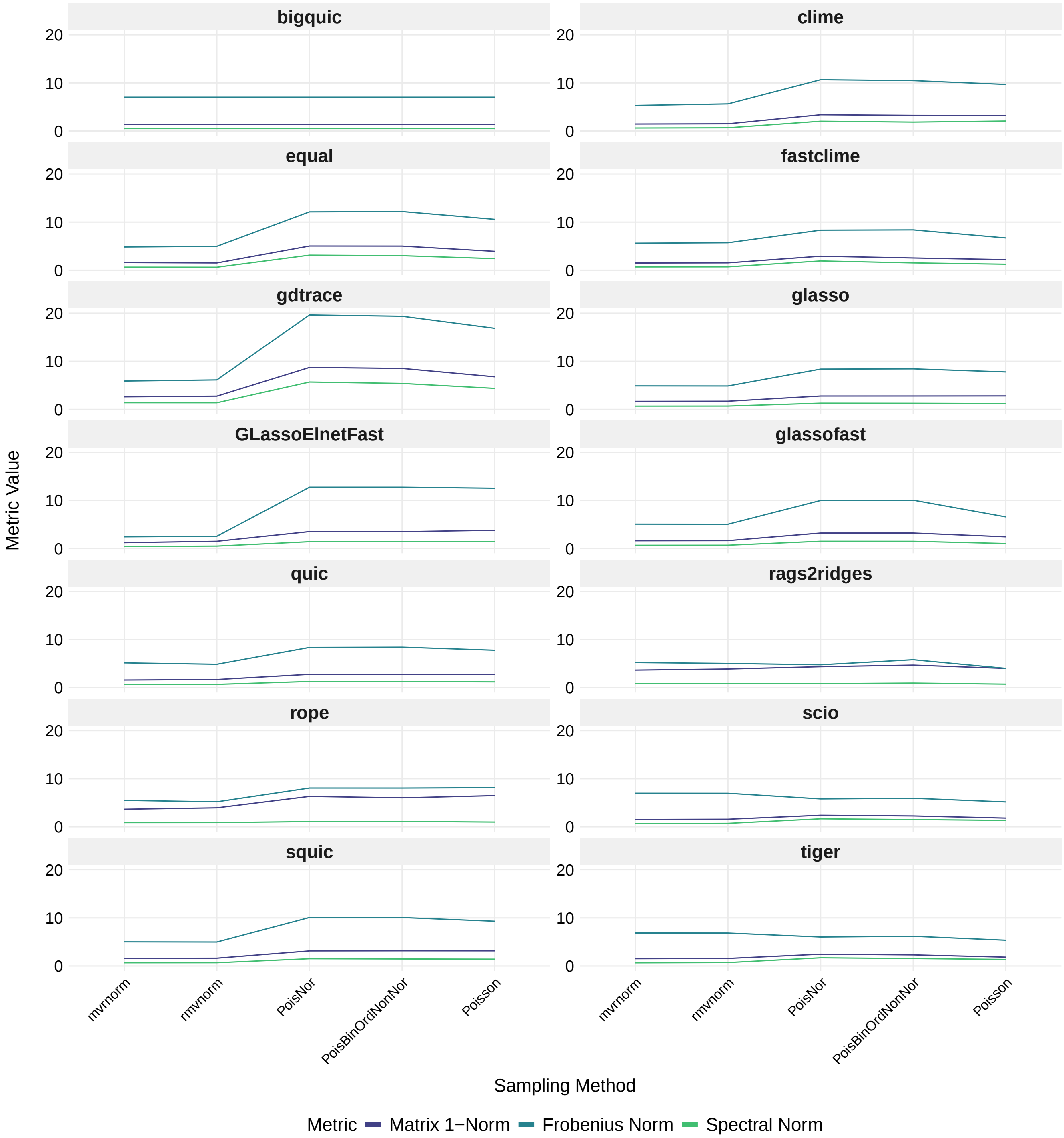

Supplementary Figure 34: Line plot illustrating the influence of different sampling methods on the performances of precision matrix estimation methods, assessed across the matrix norms matrix 1-norm, Frobenius norm, and spectral norm. $\Sigma_1$ was generated using the covariance 1 method multiple block with $p = 200$. The sample data set was created using the respective sampling method with $n = 50$.

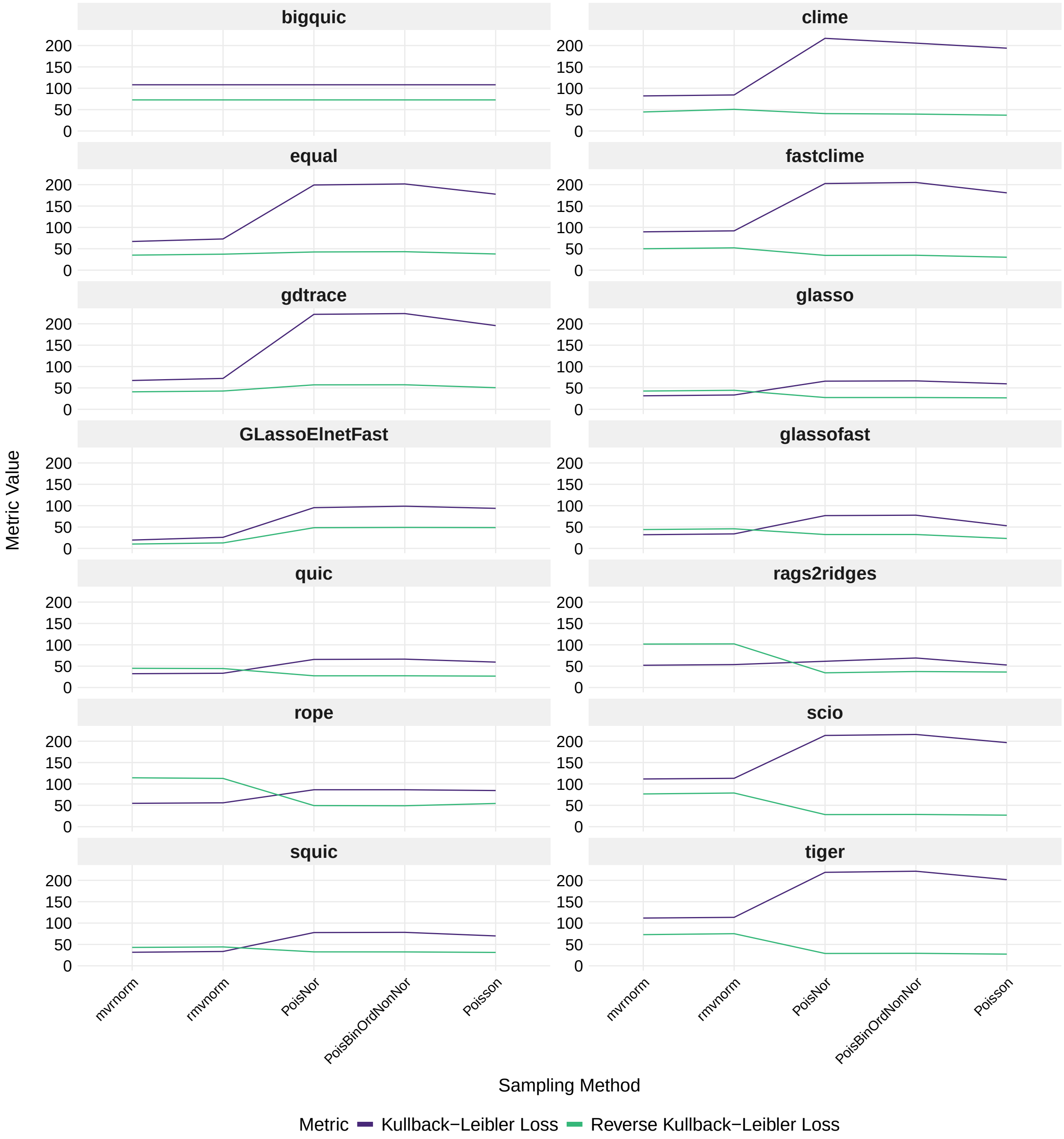

Supplementary Figure 35: Line plot illustrating the influence of different sampling methods on the performances of precision matrix estimation methods, assessed across the Kullback-Leibler Loss and the reverse Kullback-Leibler Loss. $\Sigma_1$ was generated using the covariance 1 method multiple block with $p = 200$. The sample data set was created using the respective sampling method with $n = 50$.

Yahav and Shmueli (2008) [1] proposed an approach for the generation of a $p$-dimensional Poisson vector with covariance matrix $\Sigma_{Pois}$ and rate $\vec{\Lambda}$. Specifically, the method involves generating a multivariate normal vector and then transforming it into Poisson-distributed values using the cumulative distribution functions (CDFs) of both the normal and Poisson distributions. The CDF of the Gaussian and Poisson distribution are shown in Equation 1 and 2, respectively.

$$\Phi(x) = \int_{-\infty}^{x} \frac{1}{\sqrt{2\sigma^2}} e^{-\frac{u^2}{2\sigma^2}} du \tag{1}$$

$$\Xi(x) = \sum_{i=0}^{x} \frac{e^{-\lambda}\lambda^i}{i!} \tag{2}$$

Equation (1) transforms values from a normal distribution into cumulative probabilities, while Equation (2) transforms those cumulative probabilities into Poisson-distributed values, which allows the generation of Poisson random variables from normally distributed data. The algorithm proceeds as follows:

(a) Generate a $p$-dimensional normal vector $\vec{X}^N$ with mean $\vec{\mu} = 0$, variance $\vec{\sigma} = 1$, and a correlation matrix $R^N$.

(b) For each value $X_i^N$ in the multivariate normal vector $\vec{X}^N$, compute the Normal CDF, resulting in values that follow a uniform distribution between 0 and 1:

$$\Phi(X_i^N)$$

(c) Transform the uniform values $\Phi(X_i^N)$ obtained in the previous step into Poisson-distributed values by applying the inverse of the Poisson CDF with the desired rate parameter $\lambda_i$:

$$X_i^{Pois} = \Xi^{-1}(\Phi(X_i^N))$$

The vector $\vec{X}^{\text{Pois}}$ is then a $p$-dimensional Poisson vector with correlation matrix $R^{\text{Pois}}$ and rates $\vec{\Lambda}$.

[1] Yahav, I., & Shmueli, G. (2007). An elegant method for generating multivariate Poisson random variable. `doi:10.48550/ARXIV.0710.5670`

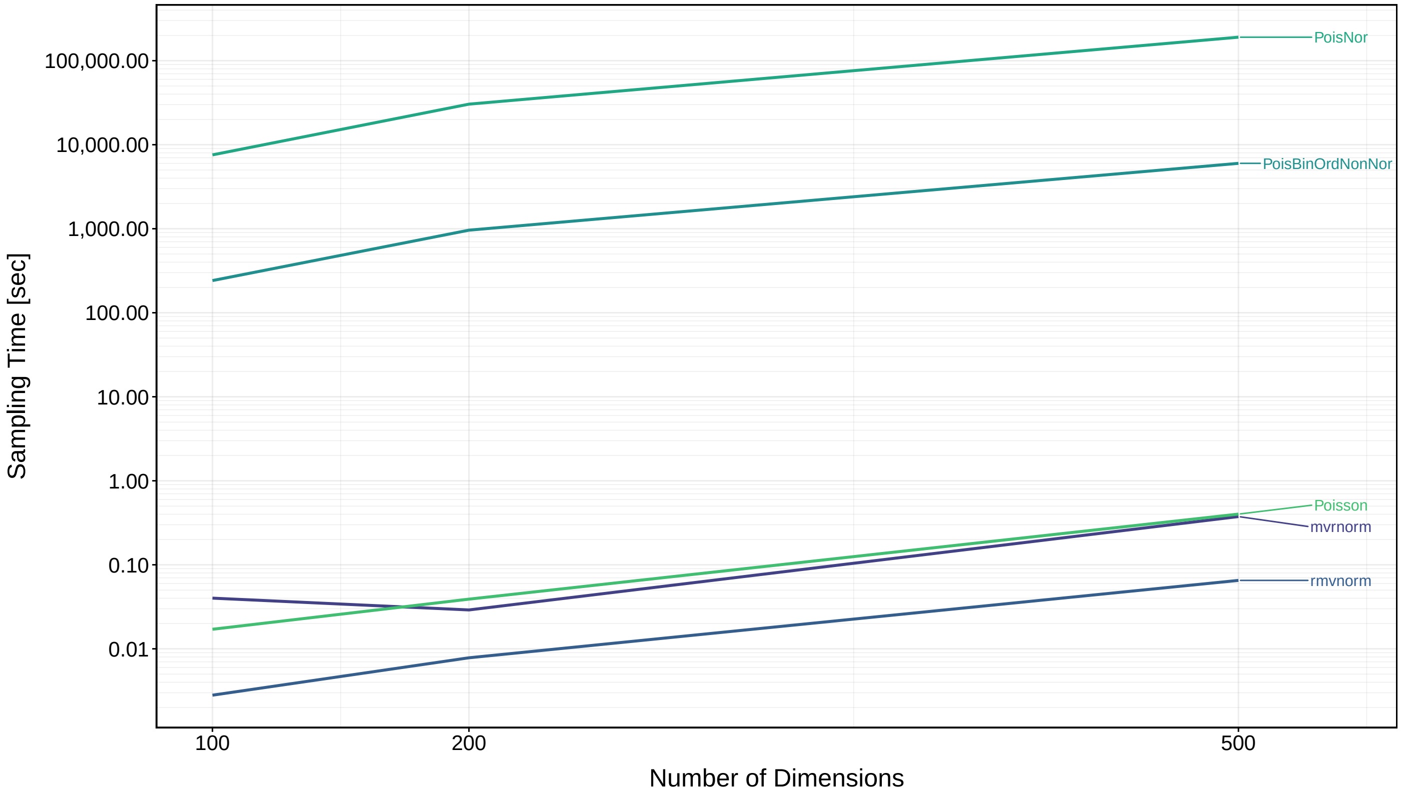

Supplementary Figure 36: Runtimes for the sampling methods mvrnorm and rmvnorm, PoisNor, PoisBinOrdNonNor, and Poisson in relation to the amount of variables (dimensions). $\Sigma_1$ was created by the multiple block method, with a size of $100 \times 100$, $200 \times 200$ or $500 \times 500$. $\Sigma_2$ was generated using the knockout method. The respective sampling methods generated the corresponding simulated data sets for both condition using a sample size of 50. The runtime includes the generation of both data sets as well as the calculation of the sample covariance matrices $S_1$ and $S_2$.

### 12 Supplementary Information for Methods

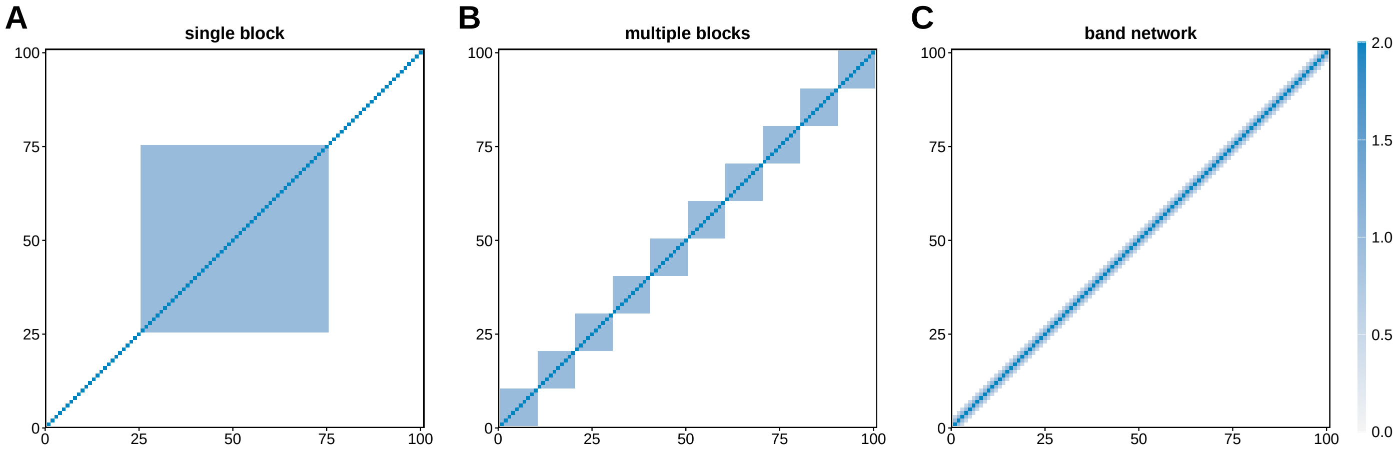

Supplementary Figure 37: Example covariance matrices generated with: single block, multiple block and band network. A: single block structure (size = 3, position = center), B: multiple blocks (cliques = 10), C: band network. All diagonal values are set to 2 and off-diagonal values in single block and multiple blocks are set to 1.

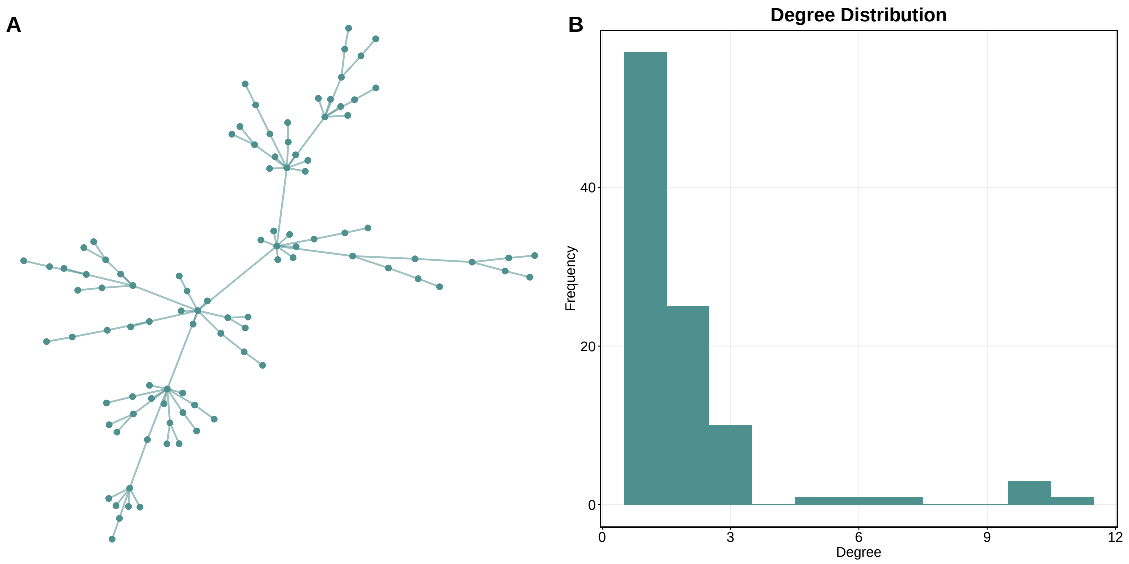

Supplementary Figure 38: Example network and degree distribution of the initial network constructed by scale free 1.

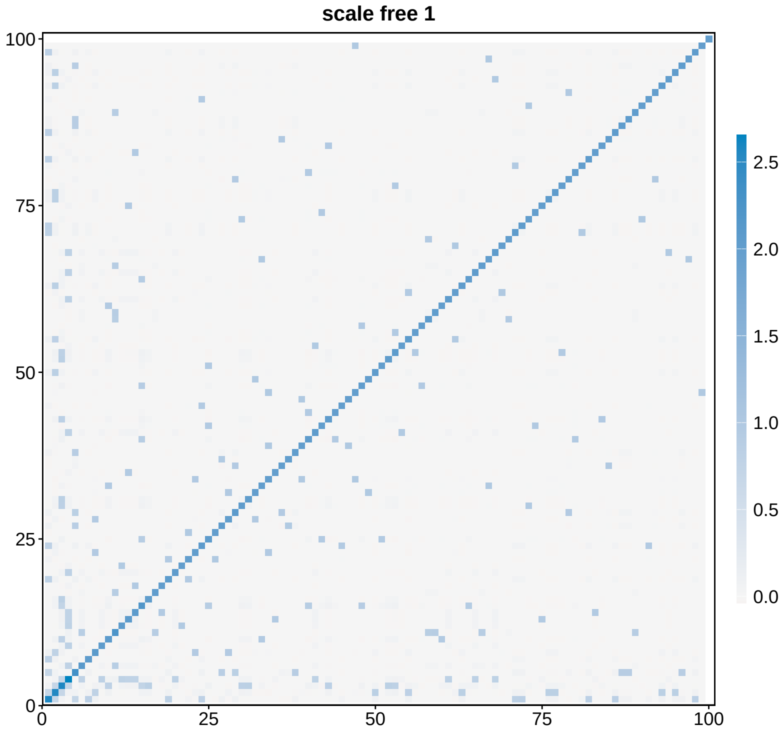

Supplementary Figure 39: Example covariance matrix generated with the scale free 1 method.

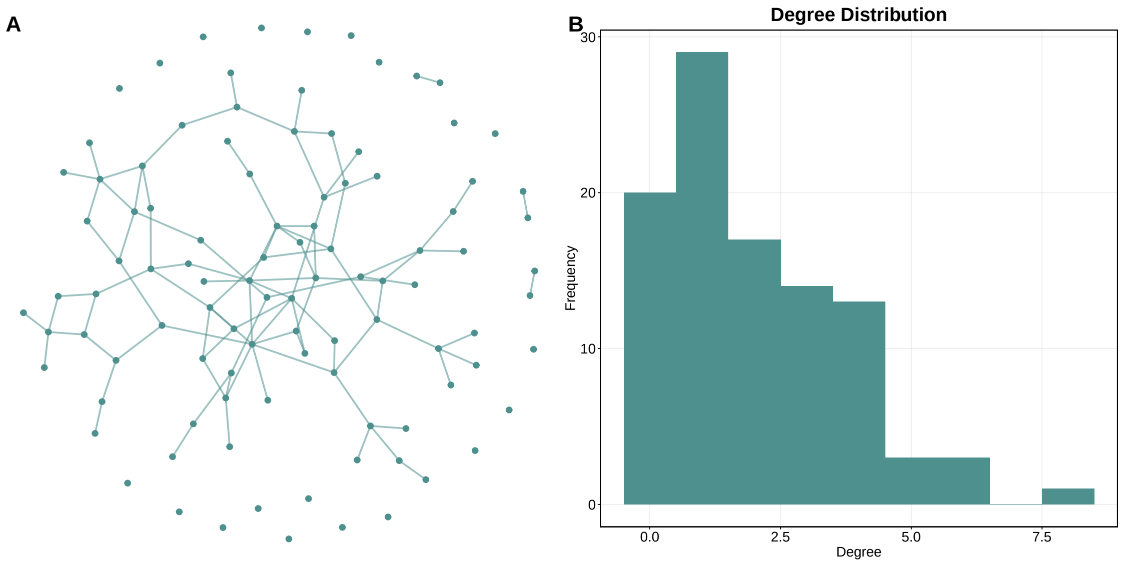

Supplementary Figure 40: Example network and degree distribution of the initial network constructed by scale free 2.

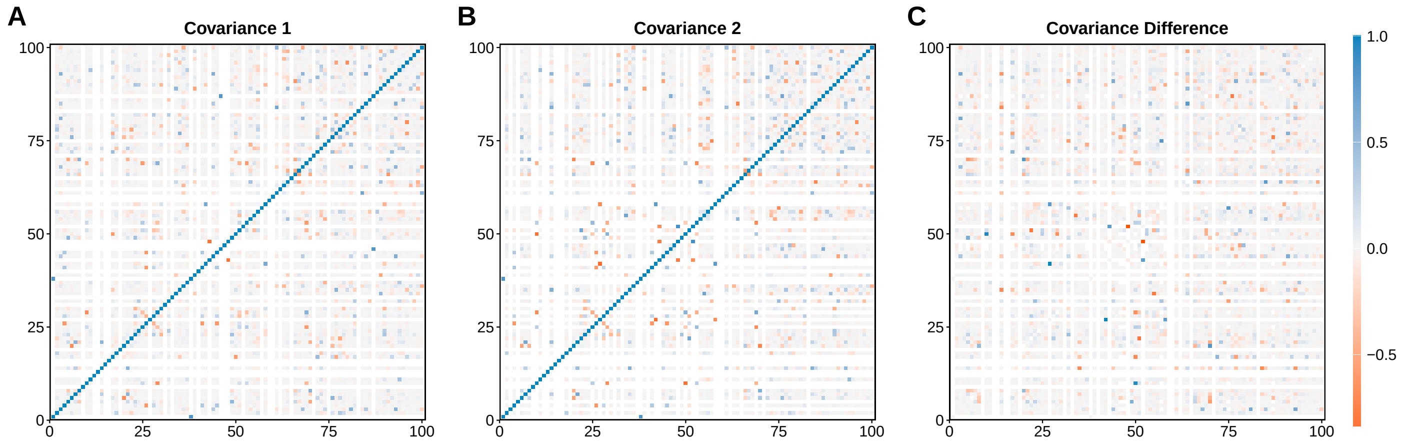

Supplementary Figure 41: Example covariance matrices ($\Sigma_1, \Sigma_2,$ and $\Sigma_\Delta$) generated with the scale free 2 method.

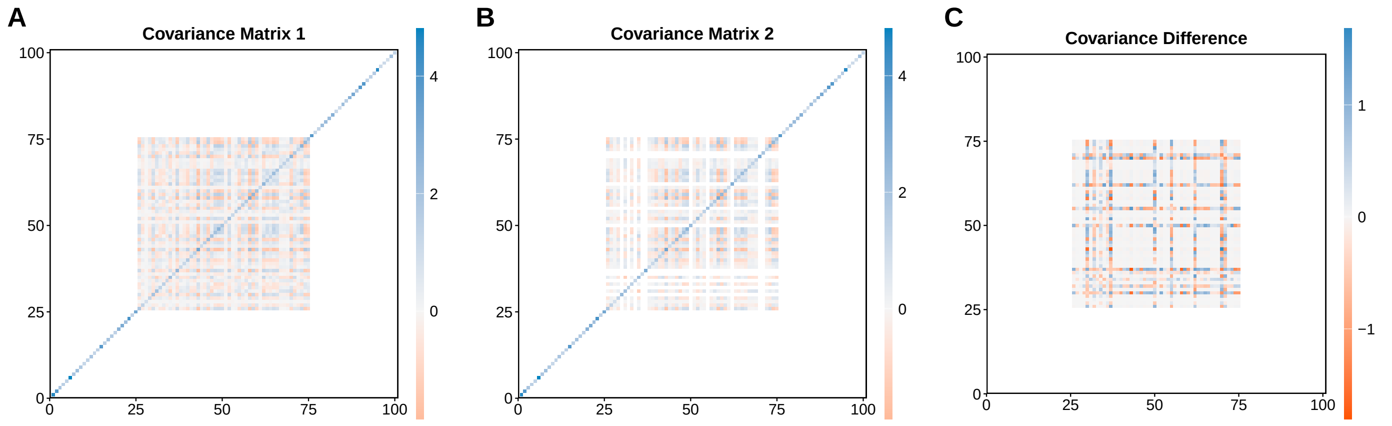

Supplementary Figure 42: Example covariance matrices generated with covariance 1 method single block icf. A: Covariance matrix for condition 1 $\Sigma_1$, B: Covariance matrix for condition 2 $\Sigma_2$, C: Differential covariance matrix $\Sigma_\Delta$.

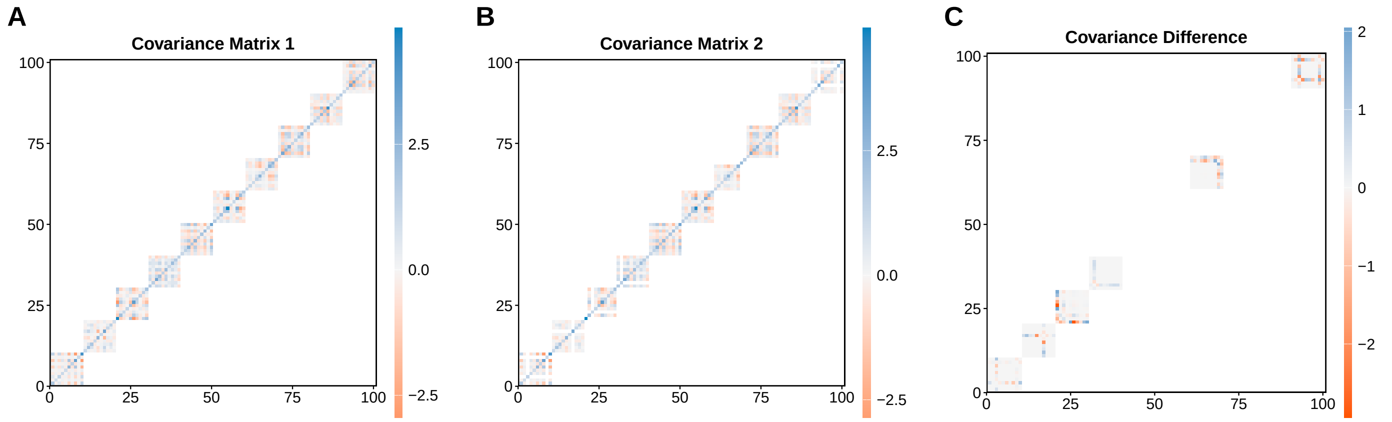

Supplementary Figure 43: Example covariance matrices generated with covariance 1 method multiple block icf. A: Covariance matrix for condition 1 $\Sigma_1$, B: Covariance matrix for condition 2 $\Sigma_2$, C: Differential covariance matrix $\Sigma_\Delta$.

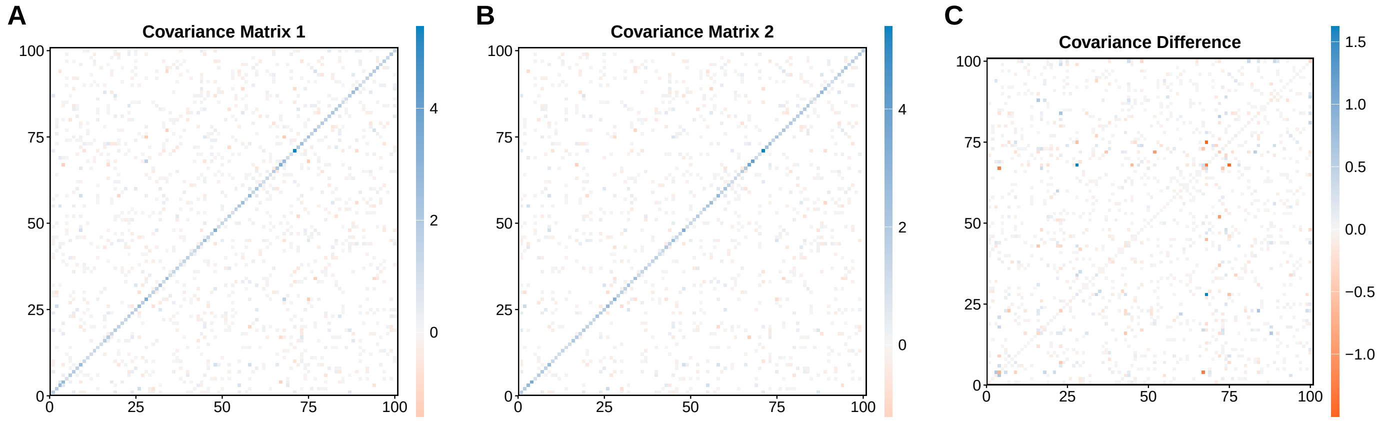

Supplementary Figure 44: Example covariance matrices generated with covariance 1 method random graph icf. A: Covariance matrix for condition 1 $\Sigma_1$, B: Covariance matrix for condition 2 $\Sigma_2$, C: Differential covariance matrix $\Sigma_\Delta$.

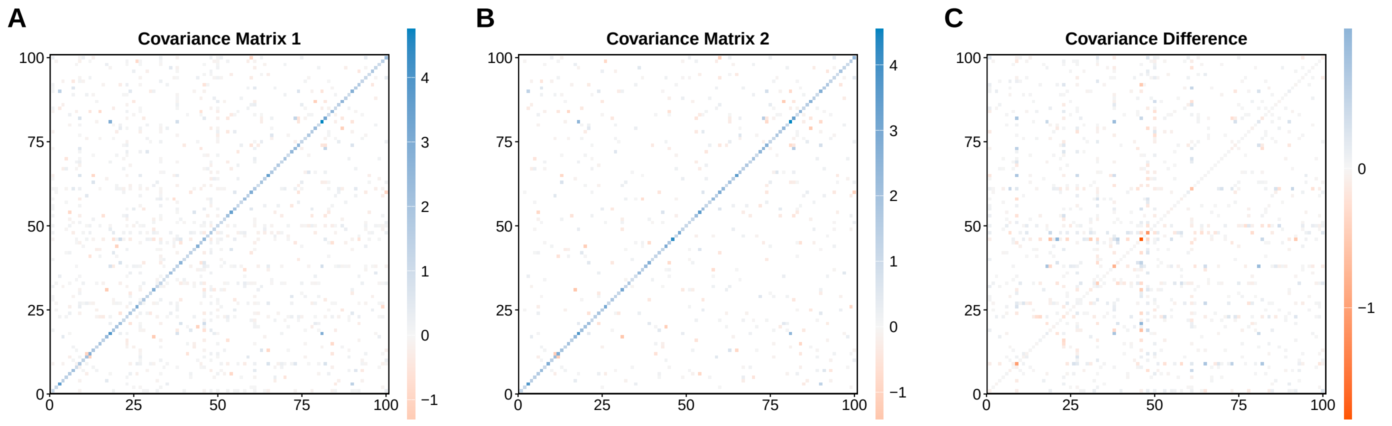

Supplementary Figure 45: Example covariance matrices generated with covariance 1 method random hubs icf. A: Covariance matrix for condition 1 $\Sigma_1$, B: Covariance matrix for condition 2 $\Sigma_2$, C: Differential covariance matrix $\Sigma_\Delta$.

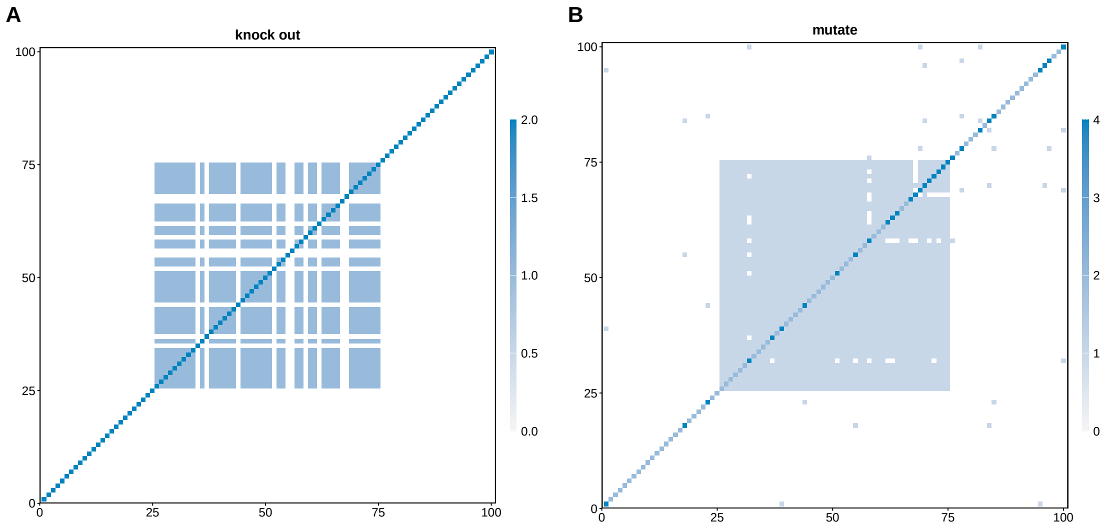

Supplementary Figure 46: Covariance 2 methods. A: knockout, B: mutate.

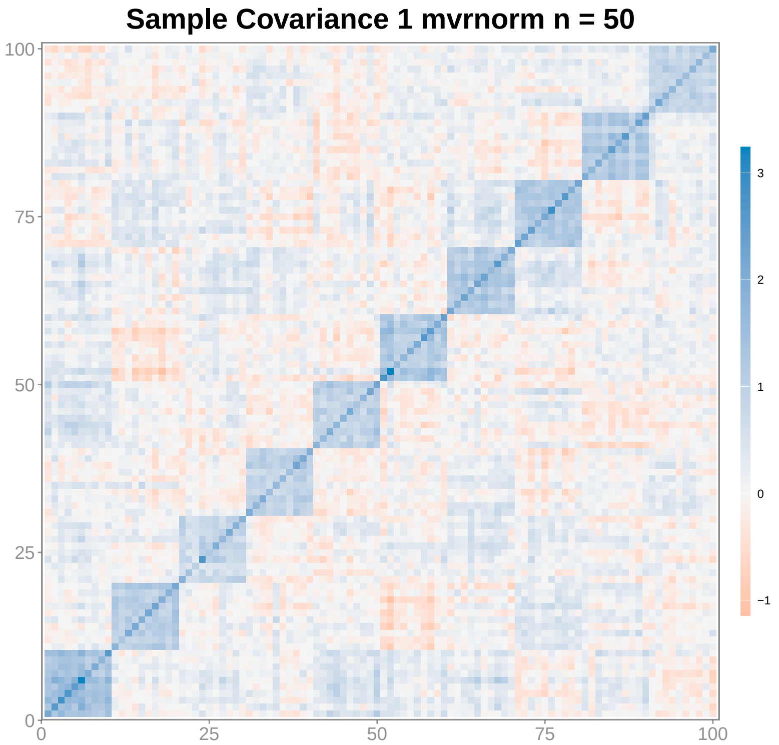

Supplementary Figure 47: Example sample covariance matrix $S_1$, generated with the sampling method mvrnorm with $n = 50$.

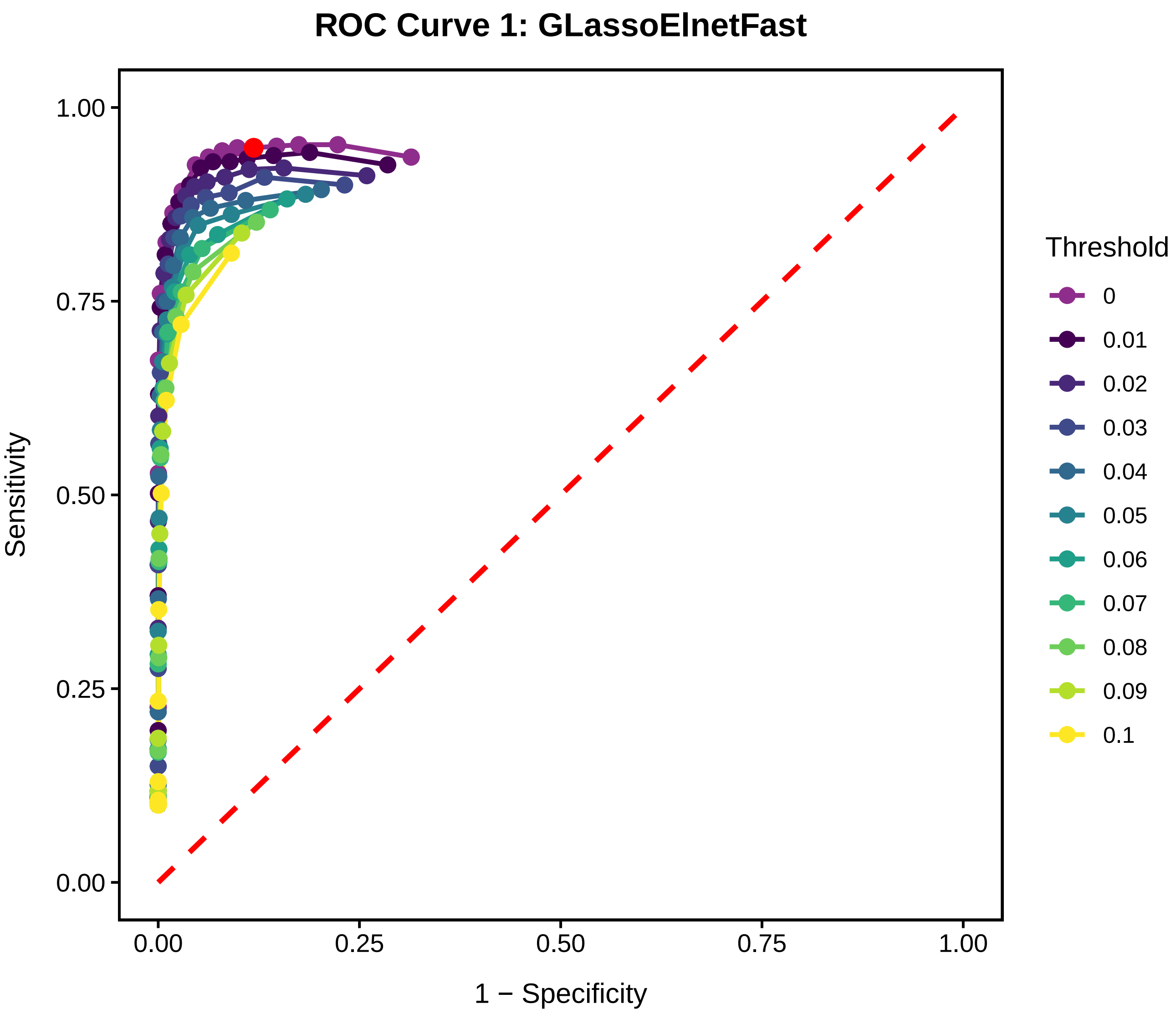

Supplementary Figure 48: Example ROC curve for estimated precision matrices obtained with *GLassoElnetFast* for condition one. Each coloured line represents one set of precision matrices, obtained by the PMEM with different regularization parameter values, with each point corresponding to one estimated precision matrix. A single precision matrix is selected as the final estimate, either via an internal regularization parameter selection or through external cross-validation. The sensitivity and 1-specificity of the selected precision matrix are indicated by a larger red dot. Different coloured lines correspond to different threshold levels applied to the estimated precision matrices. For each threshold, entries with absolute values below the threshold are set to zero, and the sensitivity and specificity of the resulting thresholded matrix are computed.

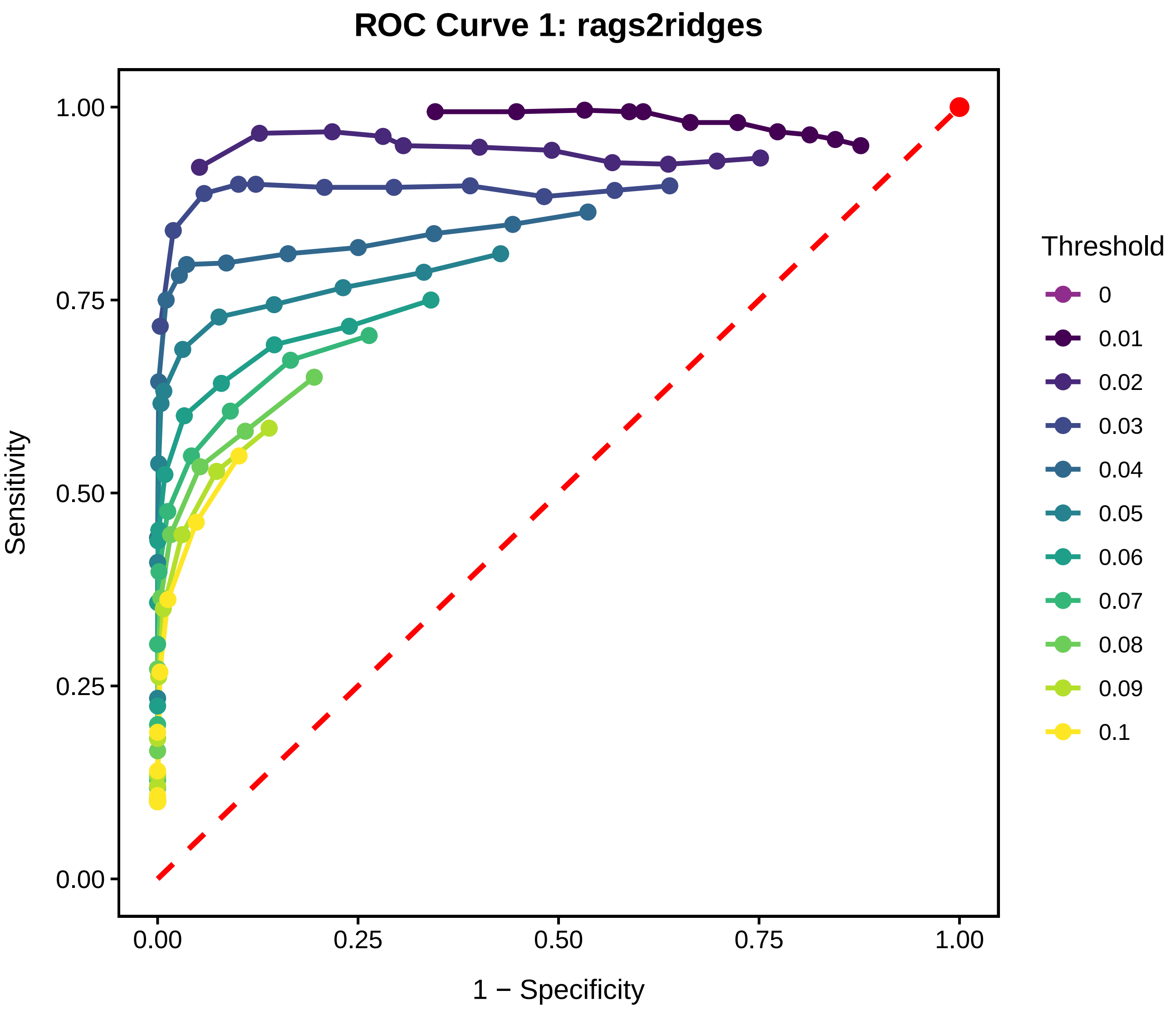

Supplementary Figure 49: Example ROC curve for estimated precision matrices obtained with *rags2ridges* for condition one. Each coloured line represents one set of precision matrices, obtained by the PMEM with different regularization parameter values, with each point corresponding to one estimated precision matrix. A single precision matrix is selected as the final estimate, either via an internal regularization parameter selection or through external cross-validation. The sensitivity and 1-specificity of the selected precision matrix are indicated by a larger red dot. Different coloured lines correspond to different threshold levels applied to the estimated precision matrices. For each threshold, entries with absolute values below the threshold are set to zero, and the sensitivity and specificity of the resulting thresholded matrix are computed.
